# Receptor tyrosine kinase inhibition leads to regression of acral melanoma by targeting the tumor microenvironment

**DOI:** 10.1101/2024.06.15.599116

**Authors:** Eric A. Smith, Rachel L. Belote, Nelly M. Cruz, Tarek E. Moustafa, Carly A. Becker, Amanda Jiang, Shukran Alizada, Tsz Yin Chan, Tori A. Seasor, Michael Balatico, Emilio Cortes-Sanchez, David H. Lum, John R. Hyngstrom, Hanlin Zeng, Dekker C. Deacon, Allie H. Grossmann, Richard M. White, Thomas A. Zangle, Robert L. Judson-Torres

## Abstract

Acral melanoma (AM) is an aggressive melanoma variant that arises from palmar, plantar, and nail unit melanocytes. Compared to non-acral cutaneous melanoma (CM), AM is biologically distinct, has an equal incidence across genetic ancestries, typically presents in advanced stage disease, is less responsive to therapy, and has an overall worse prognosis. Independent analysis of published genomic and transcriptomic sequencing identified that receptor tyrosine kinase (RTK) ligands and adapter proteins are frequently amplified, translocated, and/or overexpressed in AM. To target these unique genetic changes, a zebrafish acral melanoma model was exposed to a panel of narrow and broad spectrum multi-RTK inhibitors, revealing that dual FGFR/VEGFR inhibitors decrease acral-analogous melanocyte proliferation and migration. The potent pan-FGFR/VEGFR inhibitor, Lenvatinib, uniformly induces tumor regression in AM patient-derived xenograft (PDX) tumors but only slows tumor growth in CM models. Unlike other multi-RTK inhibitors, Lenvatinib is not directly cytotoxic to dissociated AM PDX tumor cells and instead disrupts tumor architecture and vascular networks. Considering the great difficulty in establishing AM cell culture lines, these findings suggest that AM may be more sensitive to microenvironment perturbations than CM. In conclusion, dual FGFR/VEGFR inhibition may be a viable therapeutic strategy that targets the unique biology of AM.

## INTRODUCTION

Acral melanoma (AM) arises from the volar surfaces and nail units of the hand and feet, and it is biologically distinct from non-acral cutaneous melanoma (CM)^1–3^. Since AM arises in partially sun-protected areas, it frequently lacks ultraviolet DNA damage signatures and instead demonstrates complex genomic rearrangements and copy number variations. Further reinforcing the genomic differences between subtypes, the pathogenic point mutations common in CM, such as in the BRAF, NRAS, KIT, and NF1 genes, are absent in up to 45-58% of AM^4–6^. From an epidemiologic perspective, CM is most frequently observed in the individuals of European descent, and, while AM has equal incidence across all genetic ancestries, AM represents the majority of melanomas in those of African, Asian, and Hispanic descent^3^. Diagnosing early AM is challenging since patients often present with advanced stage disease, and early tumors can mimic benign lesions, leading to delayed diagnoses^2,7,8^. This leads to a greater proportion of patients progressing to or presenting with metastatic disease. At the time of metastatic progression, frontline immune checkpoint inhibitors (ICI) demonstrate poorer overall response rates and median progression free survival for AM as compared to CM^5,9,10^. Other targeted inhibitors are either not indicated, such as BRAF inhibitors due to low prevalence of BRAF mutations in AM^4,10^, or have poor clinical responses, such as KIT inhibitors^10^. Taken together, there is an unmet need to identify AM-specific therapy regimens that target its unique biology.

A major barrier to studying AM biology and identifying targeted therapies is the lack of clinically relevant and well-described model systems. While a PTEN knockout and constitutively active Braf transgenic mouse model (*Dct-CreERKI;Braf-CA;Pten-fx/fx*) has been shown to develop acral nevi and melanoma after exposure to ionizing radiation and tamoxifen, this system only models BRAF-mutated AM^11^, which comprises at most 10-20% of AM^1,3,5^. Recently, Weiss *et al.* successfully created several zebrafish ‘Fin’ melanoma systems that are driven by genes such as CRKL, GAB2 and NF1, analogous to human melanomas that lack BRAF/KIT/NRAS alterations. This zebrafish system provides a powerful tool for studying the underlying biology of acral melanocytes, progression to melanoma, and inherent drug sensitivity of premalignant melanocytes^6^.

In lieu of using transgenic organism models, many laboratories instead develop patient-derived cell lines and patient-derived xenograft (PDX) systems in immunodeficient mice for many cancer types. Unfortunately, establishing primary AM cells maintained through immortalization has proven challenging, and the field has been unable to culture the panoply of mutational backgrounds that are clinically observed^3^. Conversely, a handful of prior studies have demonstrated that PDX models maintain a broad range of AM genotypes and can be used for preclinical therapy testing. The largest AM cohort comprises 22 PDX models from Chinese patients. While these tumors have good clinical annotations, histology, and targeted single-nucleotide variant (SNV) mutations, no copy number variation (CNV) annotations were provided^12^. The second largest cohort was made commercially available but is limited by inconsistent characterization of the AM models^13,14^. Out of 15 AM PDXs, nine were confirmed to arise from acral sites, eight of which were genetically characterized with a targeted sequencing panel. Three other reports have generated small numbers of variably characterized AM PDX models that include: five Chinese AM with CDK4 pathway aberrations^15^, up to six potential AM with BRAF or KRAS mutations^16^, and a BRAF mutated AM of the heel^17^. Recent genomic observations have demonstrated that high copy number amplifications are unique to AM and may represent an additional method of classifying these tumors besides the traditional RAF/RAS point mutations^4,5,18,19^. However, most PDX models were published before these studies, and the lack of granular CNV data prevents the classification of these models by CNV pattern. To overcome these translational limitations, we have genomically, histologically, and clinically characterized 11 AM and 6 CM PDX tumors for use in AM drug discovery, and these tumor models will be available through the Preclinical Research Resource core at the Huntsman Cancer Institute.

In this report, we have developed a drug discovery pipeline that leverages published genetic and transcriptomic data from human AM tumors, the premalignant *mitfa*-CRKL zebrafish model, new and comprehensively characterized AM PDX tumor models, and transient PDX tumor cell culture to evaluate the efficacy of small molecule inhibitors against AM. The only inhibitor identified to promote stable disease or tumor regression across all AM models was Lenvatinib, a multiple receptor tyrosine kinase (RTK) inhibitor that is most potent against the VEGFR and FGFR families^20,21^. While this was a surprising finding considering the poor performance of Lenvatinib against all skin melanomas in the LEAP-003 study results (abstract O-031, 20^th^ Society Melanoma Research Congress^22^), it mirrors a small study wherein six AM patients were given the drug as a second-line therapy and four patients (66%) had an objective response^23^. Mechanistically, we demonstrate that Lenvatinib has minimal direct cytotoxicity against transiently cultured AM cells and instead induces tumor regression by remodeling the tumor vasculature. Taken together, these data provide a rationale for the clinical evaluation of Lenvatinib and other potent dual FGFR/VEGFR inhibitors against AM as a biologically distinct subset of skin melanomas.

## METHODS

### RTK-Associated Gene Mutation, Gene Expression, and Inhibitor Evaluation from Literature

A literature search performed in December 2023 identified eight studies that contained genomic, exomic, and/or transcriptomic data for AM^4,6,19,24–28^. Of these articles, four provided sufficient genomic data for the evaluation of changes in copy number, translocations, InDels, gene expression for RTK proteins, and/or structural variants^4,6,19,28^. CNV analysis was performed on WES data from 37 AM tumors available in Wang 2023^19^. Briefly, average ploidy was estimated in each tumor after removal of highly amplified ‘hailstorm’ CNV events. Fold change in copy number was determined by dividing the locus copy number by average ploidy, and the calling cut-offs for CNV events are as follows: allele loss as ≤0.5 fold change, allele amplification as ≥1.5 fold change, and high allele amplification as ≥4.0 fold change. Large InDels and translocations/structural variants were identified from 121 tumors in Liang 2022^28^ and Newell 2020^4^. Fold changes in gene expression between AM and CM were extracted from Weiss 2022^6^, and multi-RTK inhibitor cell-free IC50 and estimated IC50 based on percent inhibition studies were collated from peer-reviewed literature, public FDA documents, and, in the case of Anlotinib, company promotional material.

### Patient-Derived Xenograft (PDX) Generation and Maintenance

Tumor tissue was obtained from patients who provided written informed consent according to a tissue collection protocol (University of Utah IRB 89989 and 10924) approved by the Huntsman Cancer Institute (HCI) Institutional Review Board.

NRG mice (JAX: NOD.Cg-Rag1tm1Mom Il2rgtm1Wjl/SzJ, Strain #:007799) were maintained in a pathogen-free facility at the HCI. All animal experiments were performed in accordance with protocols approved by the University of Utah Institutional Animal Care and Use Committees, and we have complied with all relevant ethical regulations. Mice were kept in a temperature-controlled facility on a 12/12-hour light/dark schedule with standard food and water supplies.

Patient and PDX tumor tissue fragments (~15mg) were subcutaneously implanted into NRG mice. Mice were placed under inhaled isoflurane anesthesia; the incision site was prepared by alternating alcohol and betadine scrubs. For the duration of surgical procedures, mice were kept on a water-circulated heated mat at 37°C. Three minutes prior to the procedure, a local anesthetic of 5mg/kg of Lidocaine was administered subcutaneously at the incision site. A small (3-4mm) incision was made with scissors and a tumor fragment (1-3mm^3^) was implanted under the skin. The incisions were closed with a single 9-mm wound clip that was removed 7-10 days after surgery. The procedures lasted ~2 minutes. Mice were allowed to recover on a 37°C warm pad before being returned to their cage. Following growth, PDX tumors were resected under sterile conditions and biobanked as 1) formalin fixed paraffin embedded blocks, 2) flash frozen tissue, and 3) viably cryopreserved tissue^29^. For cryopreservation, tumor tissue was cut into ~15mg fragments. Tumor fragments are placed in a cryovial with tissue freezing medium (95% FBS, 5% DMSO) and frozen at −80^◦^C overnight before transferring the vials to liquid nitrogen cryotanks for long term storage.

PDX models will be made available through the Huntsman Cancer Institute Preclinical Research Resource. Please contact for additional information.

### PDX Drug Studies

PDX fragments were implanted into male NRG mice. Tumor-bearing mice were randomly enrolled into treatment groups based on tumor size. These mice were subjected to various treatment regimens to assess anti-tumor efficacy. The treatment groups included control (vehicle-treated) mice and experimental groups receiving different drugs outlined in **Table S2**. Tumor size was measured twice weekly in two dimensions using calipers, and the volume was expressed in mm^3^ using the formula: V = 0.5 a x b^2^ where a and b are the long and short diameters of the tumor, respectively.

### Chart Review and Clinical Slide Imaging

Clinical chart review, slide procurement, and slide imaging were performed under the ARUP umbrella IRB protocol (#00091019) for general pathology specimens and the Dermatopathology umbrella IRB protocol (#00076927) for dermatopathology specimens. General case information such as gender, self-identified ethnicity, age at diagnosis, primary tumor origin, stage at diagnosis, PDX tumor origin, patient alive/dead status as of 3/2024, and treatment history were collected through the Huntsman Cancer Institute’s Research Informatics Shared Resource (RISR). Where possible, this information was confirmed through independent chart review and expanded to include clinically identified mutations and source of molecular pathology tissue (Table S1). All diagnoses were independently confirmed and imaged by a pathologist using an Olympus BX53 microscope equipped with Olympus PLAN N (2x/0.06 ∞/-/FN22) and UPlanFL N (4x/0.13 ∞/-/FN26.5, 10x/0.30 ∞/-/FN26.5, 20x/0.50 ∞/0.17/FN26.5, 40x/0.75 ∞/0.17/FN26.5) optics, a DP74 camera, and Olympus cellSens Entry 1.18 (Build 16686) software.

### Histologic Drift Score Development and Validation

A histology-based tool was developed to evaluate the changes in PDX tumor cytology and histological architecture over time compared to the parental clinical tumor. A pathologist, EAS, developed the scoring system and assigned ‘development’ scores to each PDX passage based on the criteria in Supplemental Data 3 as part of a longitudinal review of PDX tumor cases. For score validation, two cytopathologists, TAS and MB, independently compared and scored each clinical case, low passage PDX tumor, and high passage PDX tumor. One slide of each PDX tumor at low passage (passage 1-2) and high passage (passage 3-5) was evaluated and scored according to feature concordance with the clinical tumor of origin. Full scoring details are listed in Supplemental Data 3, but a brief description of scores are: Score 0 – identical to clinical tumor; Score 1 – minimal changes secondary to pigment, vascularity, stroma, necrosis, and/or shift between related nodular/alveolar and fascicular/storiform architectures; Score 2 – a partial shift in cytology, a partial change in architecture, or a secondary clinical architecture/cytology now predominates; Score 3 – partial shift in cytology and architecture, complete change in cytology, or complete change to a novel architecture; Score 4 – tumor is unrecognizable compared to clinical tumor.

### Zebrafish husbandry

The zebrafish transgenic strains used were casper MiniCoopR *mitfa*:EGFP and casper MiniCoopR:EGFP, *mitfa*:CRKL stable lines. Fish stocks were kept at 28.5 °C under 14:10 light:dark cycles, pH (7.4), and salinity-controlled conditions. The fish were fed a standard diet consisting of brine shrimp followed by Zeigler pellets. The animal protocols were approved by the Memorial Sloan Kettering Cancer Center (MSKCC) Institutional Animal Care and Use Committee (IACUC), protocol number 12–05-008. Individual mating pairs were crossed and collected embryos were incubated in E3 medium (5mM NaCl, 0.17mM KCl, 0.33mM CaCl_2_, 0.33mM MgSO_4_) at 28.5°C. Anesthesia of embryos was performed using Tricaine-S (MS-222, Syndel) with a 4g/L, pH 7.0 stock diluted in E3 medium to a final concentration of 250mg/L.

### Pharmacological treatment of zebrafish embryos

Zebrafish embryos were treated with the following compounds (purchased from Selleck Chemicals) at the indicated concentrations: Anlotinib (S8726), Apatinib (S5248), Cabozantinib (S1119), Lenvatinib (S1164), Sunitinib (S7781). Groups of twenty 24 hr post-fertilization (hpf) embryos were randomly selected from a single clutch and placed in a 70-μm cell strainer (Falcon 352350) submerged in E3 medium. The strainers were then transferred to a 6-well dish (Fisher 08–772-1B) containing 6 mL of either 3μM or 1μM compound diluted in E3 medium. Treated zebrafish were imaged at 72 hpf using a Zeiss Axio Zoom.V16 stereomicroscope. Images were used to measure GFP melanophore cell area in the tailfin mesenchyme, as previously described^6^. Embryo body length and yolk sac area were measured from scale-calibrated images with ImageJ. Treatment experiments were performed on three separate occasions.

### Targeted ArcherDx Gene Panel Sequencing

A targeted massively parallel gene sequencing panel was used to identify single nucleotide variants (SNV) and copy number variants (CNV) for each CM and AM PDX tumor. DNA was extracted from fresh frozen PDX tissue using the QIAMP DNA mini kit (Qiagen 56304) or the Qiagen All Prep DNA/RNA kit (Qiagen 80204), and 150ng was used for a custom anchored multiplex PCR protocol from ArcherDx (IDT)]. This probe set contains primers for 34 melanoma-associated genes: ARID1A, NRAS, NOTCH2, RAF1, BAP1, PBRM1, MITF, PDGFRA, KIT, TERT, ARID1B, EGFR, MET, BRAF, CDKN2A, PTEN, HRAS, CCND1, GAB2, KRAS, ARID2, CDK4, MDM2, BRCA2, RB1, SPRED1, MAP2K1, MC1R, TP53, NF1, BRCA1, MAP2K2, SMARCA4, and CRKL. Eight additional genes only had hotspot mutation coverage: CTNNB1, EZH2, GNA11, GNAQ, PPP6C, RAC1, SF3B1, and STK19. Library preparation was performed according to the Archer VariantPlex HS/HGC protocol for Illumina sequencing, and libraries were quantified with KAPA library quantitation (Roche, KR0405) before sequencing on an Illumina NovaSeq 6000 with 25% of reads PhiX per manufacturer’s recommendations. FASTQ files were uploaded and analyzed on the Archer Analysis Unlimited (v7.1) website to a read depth of 12 million reads per sample. These samples were CNV normalized to a control data set containing eight foreskin samples performed with the same methodology.

Analysis of the targeted NGS genomic sequencing panel was performed by a molecular pathologist. Single nucleotide variants (SNV) were identified using the following filters: allele frequency >0.027, alternative observations ≥5, and unique alternative observations ≥3, depth ≥250, and variant allele frequency (VAF) ≥10%. SNV allele homozygosity was defined as ≥75% VAF, heterozygous as 33-75% VAF, and low VAF as <33% VAF. SNVs were classified according to the Association for Molecular Pathologists standards and guidelines^30^ with variant review in ClinVar, COSMIC, and PubMed databases. CNV were determined by evaluating log2 fold change within a gene with the following variant calling thresholds: deletion at <0.1 fold change, loss at 0.1-0.6 fold change, and amplification at >1.75 fold change. To call focal or partial gene CNV a stretch of ≥6 primers must be at or within the indicated threshold. Full gene CNV is called when the majority of primers are within the indicated threshold, and the remainder of primers approximate the threshold cut-off. Highly amplified genes (“Amplification ≥8 copies”) have >8 observed copies across multiple primers.

### Transient Dissociated PDX Cell Culture and Quantitative Phase Imaging

Cryopreserved PDX tumor tissue is implanted into athymic nude mice and allowed to grow to 500-1000mm^3^ prior to harvesting for cell dissociation. Tumors are dissected into 200mg chunks in 4.7ml of RPMI and treated with 300μl of tumor dissociation enzymatic mix (Milteny, 130-095-929) at 37⁰C using the 37C_h_TDK_1 protocol on a gentleMACS™ Octo Dissociator with Heaters (Milteny, 130-096-427). After a 10 min 500g centrifugation, the cell pellet is resuspended in cold RPMI, and the cells are quantified using a Countess® II FL Automated Cell Counter with trypan blue. A repeat 10min 500g centrifugation is performed, and the cell pellet it resuspended in 80μL of mouse cell depletion kit buffer (Milteny, 130-104-694) and 20µL of mouse cell depletion kit beads per 10^7^ cells. After incubating at 4⁰C for 15 minutes, the volume is adjusted to 500μL/10^7^ cells with autoMACS® Rinsing Solution (Milteny, 130-091-222), and the samples are loaded onto an autoMACS® Pro Separator (Milteny, 130-092-545). The DEPLETES program is ran with an autoMACS® column (Milteny, 130-021-101) to enrich for tumor cells prior to seeding a 96 well plate at 4000 cells/well. Cells were cultured in Mel-2 media (aka ‘Mel2%’)^31^ which contains 400ml MCDB153 media (Sigma M7403-10X1L), 100ml Leibovitz’s L-15 media (Gibco 11415-064), 2.5% fetal bovine serum (Denville FB5001-H), 1.68mM CaCl2, 5ml Insulin-Transferrin-Selenium X (Gibco 51500-056), 5ng/ml EGF (Sigma E-9644), 15µg/ml bovine pituitary extract (Gibco 13028-014), and 1x Pen/Strep (Gibco 15140122).

QPI was performed using differential phase contrast (DPC)^32^ on a custom-built microscope^33^ with 50ms exposure time, coherence parameters (σ) of 1.25, and regularization parameter of 4×10^-3^. Images were acquired using an 10x, NA = 0.25 objective (Olympus, Tokyo, Japan), captured on a monochrome 1920 × 1200 CMOS camera (FLIR imaging, OH, USA) with illumination from an 8×8 0.1” spacing LED array (Adafruit, NY, USA). Cells were segmented using a Sobel filter and morphological operators to create a mask and separated into single cells using a watershed algorithm using custom Matlab code (Mathworks, MA, USA). A refractive increment of 1.8×10^−4^m^3^/kg was assumed for calculation of cell mass^34^. Specific growth rate, normalized mass, depth of response, time of response, EC_50_, and heterogeneity were calculated as previously described^35^. The following drug concentrations in were evaluated: 0.5% DMSO vehicle control; Apatinib at 0.0064, 0.064, 0.32, 1.6, 8, and 40µM; Lenvatinib at 0.0016, 0.016, 0.08, 0.4, 2, and 20µM; and Sunitinib at 0.008, 0.08, 0.4, 2, 10, and 50µM.

### Immunohistochemistry (IHC)

IHC was performed on Leica Bond Rx instrument through the Huntsman Cancer Institute BMP Core Facility. Heat induced epitope retrieval was performed at 95°C using Bond Epitope Retrieval Solution 1 (Leica Biosystems AR9961) for transferrin receptor 1 (TfR1) and using Bond Epitope Retrieval Solution 2 (Leica Biosystems AR9640) for Ki67 and CD31. All slides were washed five times with 10% H2O2 at 55°C to reduce melanin content. Primary antibody incubation was performed as follows: Ki67 (Cell Signaling #12202S) was incubated at 1:300 for 15 minutes, CD31 (Invitrogen pa5-16301) was incubated at 1:50 for 60 minutes, and TfR1 (Abcam ab214039) was incubated at 1:1000 for 15 minutes. Both TfR1 and CD41 were stained with the Leica Bond Polymer Refine Detection Kit DS9800, and Ki67 was stained with the Leica Bond Polymer Refine Red Detection Kit DS9390.

A board-certified pathologist blindly reviewed all IHC stains and scored or quantified the stains as follows: Ki67 as percent positive cells within a tissue section of tumor, CD31 as number of positive non-continuous vessels per 10x field, and membranous TfR1 by average intensity (1+ = weak/blush; 2+ = moderate/strong staining of membrane, 3+ very strong staining) and percentage of staining non-necrotic tumor cells.

### Data Availability

Data used for the RTK meta-analysis were previously published as supplementary data files in their corresponding articles^4,19,28^, and the analysis used to generate the graphs in Figure 1 are available in Supplemental Data 1. Supplemental Data 4 and 5 contain all SNV calls and CNV probe results from the ArcherDx gene panel, and representative histology images of each PDX tumor and its corresponding clinical tumor are available in Supplemental Data 6.

**Figure 1:**
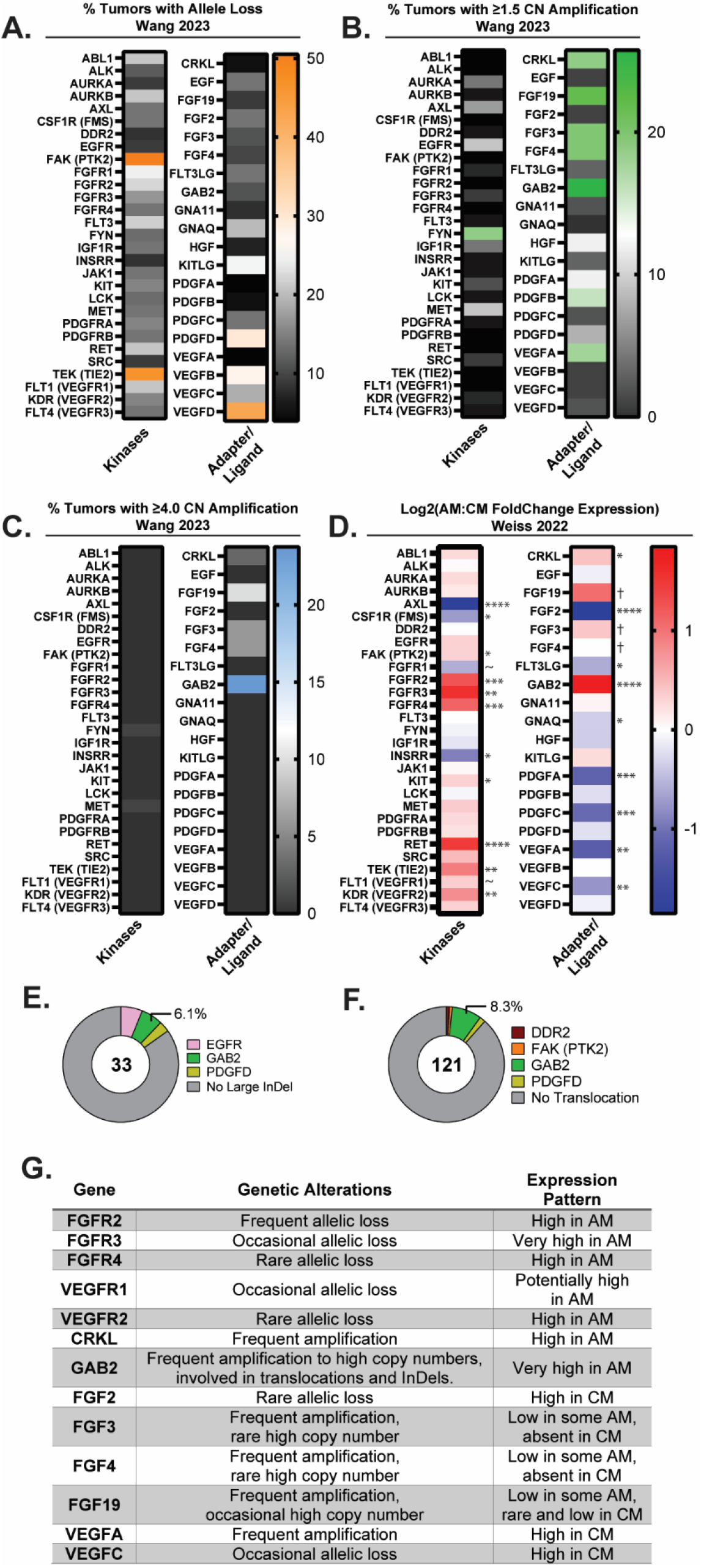
Acral melanoma (AM) tumors highly amplify and/or upregulate RTK adapter proteins, VEGFR, FGFR, and FGF ligands. **(A)** From 88 tumors in the Wang 2023 cohort, the percentage of tumors bearing RTK-associated allele loss, **(B)** amplification, or **(C)** high copy-number amplification above 4x background tumor ploidy are shown. **(D)** Translocation events involving RTK-associated genes occur in a minority of AM per Liang 2017^83^ and Newell 2020^4^. In comparison to CM, AM highly express RTK and intracellular RTK adapters (CRKL and GAB2) per Weiss 2022^6^. Certain ligands such as FGF3 and FGF19 were expressed at low levels in some AM tumors compared to lack of expression in CM. P-adjusted significance values are indicated as follows: ~ p=0.05-0.07, * p<0.05, **p<0.01, ***p<0.001, ****p<0.0001, † low expression gives error in p-value calculations. (**E)** Large insertion and deletion (InDel) events and (**F**) translocations infrequently involve RTK-associated genes. **(G)** A summary of published CNV and expression results for select genes are tabulated.

## RESULTS

### Receptor Tyrosine Kinase (RTK) associated proteins are highly amplified and expressed in AM

Candidate drug classes were first identified from published genomic and transcriptomic data (**Figure 1**). A meta-analysis of available single nucleotide variation (SNV), copy number variation (CNV), structural variation (SV), and transcriptomic data was performed for AM. Eight volar melanocyte and/or AM studies were identified to contain whole genome sequencing (WGS), whole exome sequencing (WES), and/or RNA-sequencing data (RNA-seq)^4,6,19,24–28^. Of these, three provided sufficient genomic data for the evaluation of InDels, CNV, and/or structural variants^4,19,28^, and one provided RNA-sequencing data^6^. CNV analysis was performed on WES data from 37 AM tumors available in Wang 2023^19^ and revealed that RTK-associated genes are variably lost (**Figure 1A**, Supplemental Data 1) and commonly amplified (**Figure 1B-C**, Supplemental Data 1). Of these genes, CRKL, FGF19, FGF3, FGF4, and GAB2 had high levels of amplification (≥4x the background ploidy) in ≥5-20% of AM. These genes are present within loci that commonly undergo tyfonas/hailstorm events, which can produce SV and high CNV on the order of 10-100x baseline ploidy as described in previous reports^18,19^. Interestingly, only the amplified genes present within tyfonas-affected loci had a corresponding fold change increase in RNA expression for AM compared to CM (**Figure 1D**). For the FGF3/4/19 ligands, most tumors expressed at or below the RNAseq limit of quantitation. Only 6 of 53 (11%) CM tumors expressed FGF19 above this limit, and no FGF3 or FGF4 transcripts exceeded this threshold. AM had a larger percentage of tumors expressing these ligands (20 of 61 cells, 33%) with 16 having increased FGF19 expression, 6 with increased FGF3 expression, 1 with increased FGF4 expression, and 3 with overlapping expression of multiple of these ligands. RTK-associated genes are infrequently involved in large insertion/deletion (InDel) or translocation/SV events, with GAB2 being the most commonly involved gene (**Figure 1E-F**, Supplemental Data 1). Compared to the adapters and ligands, certain families of kinase receptors, such as VEGFR and FGFR, were highly expressed in AM despite being rarely amplified or involved in SV, InDel, or tyfonas events (**Figure 1G**). Considering that the FGF-FGFR axis and VEGFR family are highly amplified and/or expressed, we hypothesized that a dual FGFR/VEGFR inhibitor could be uniquely efficacious in AM tumors.

### Multi-RTK inhibitors inhibit embryonic melanogenesis via FGFR and VEGFR blockade

Several FGFR and VEGFR inhibitors have been developed and used clinically, with each having a unique kinome inhibition profile^36^. Based on their cell-free biochemical 50% inhibitory concentration (IC_50_) and kinase inhibitory percentage profiles, we built a panel of RTK inhibitors to probe which pathways, or combination of pathways, are essential for AM tumor cell survival. For this drug panel (**Figure 2A**) we selected a variety of potent VEGFR inhibitors that include Apatinib (Rivoceranib) as an ultra-narrow-spectrum VEGFR2/RET inhibitor^36,37^, Anlotinib as a narrow-spectrum FGFR1/pan-VEGFR inhibitor (^38,39^ and company promotional materials), Lenvatinib as a potent pan-VEGFR and pan-FGFR family inhibitor^20,21^, Cabozantinib as a moderately broad inhibitor with no activity against FGFR (^40^ and FDA pharmacology review application number 208692Orig1s000, 12 Oct 2015), and Sunitinib as a broad kinome inhibitor with no FGFR activity (IC_50_ reported in FDA pharmacology review application number NDA 21-938 and NDA 21-968, 10 Aug 2005).

**Figure 2:**
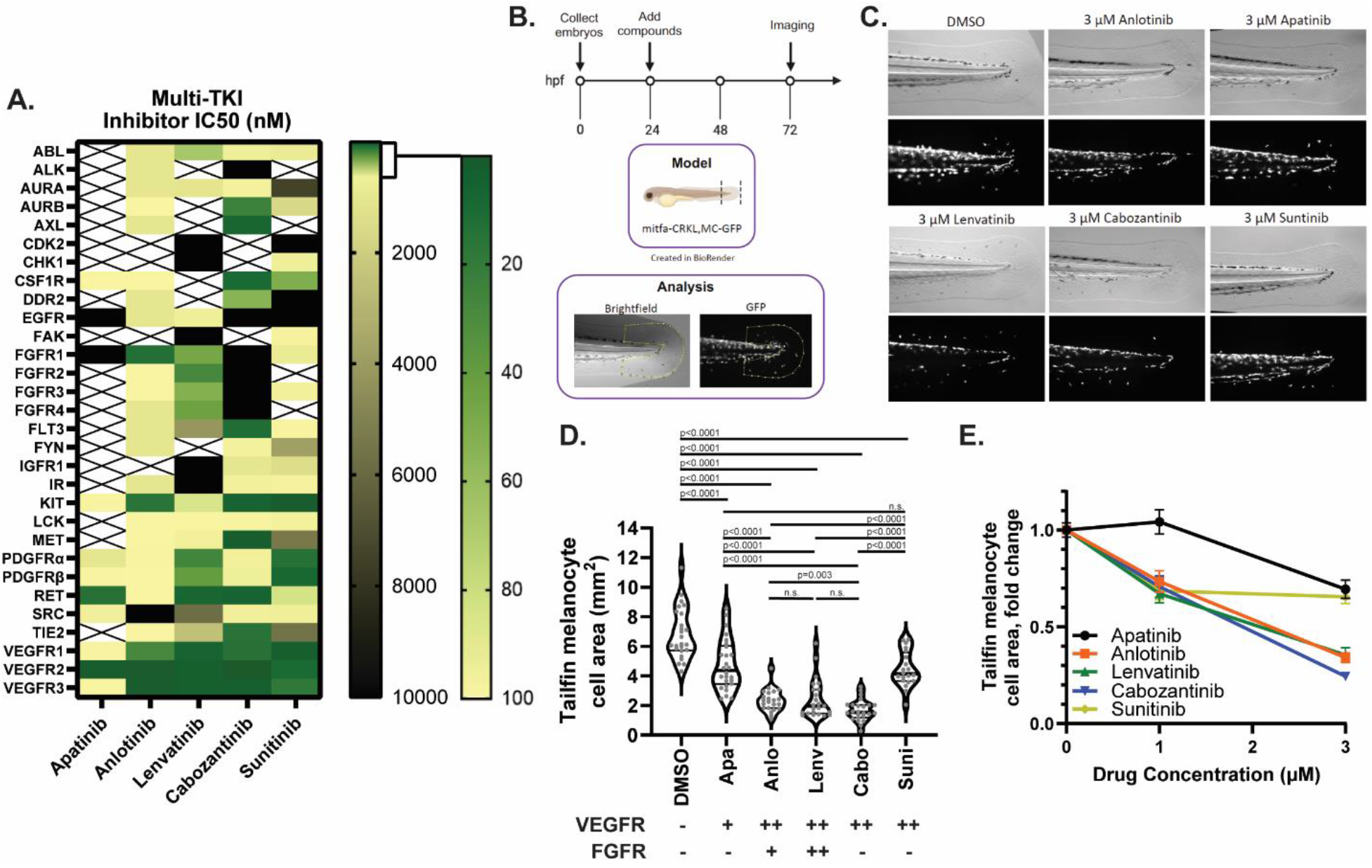
Blockade of FGFR/VEGFR receptors inhibits melanogenesis in a dose-dependent fashion. **(A)** Colormap representing the published IC50 values and inferred IC50 values from kinase inhibition studies for five separate multi-RTK inhibitors. **(B)** A premalignant mitfa-CRKL and MC-GFP zebrafish model of AM is utilized to test multi-RTK inhibitor effects against fin melanogenesis. **(C-D)** Phase contrast and GFP fluorescence imaging of multi-RTK treated zebrafish tails reveal that 3µM Anlotinib, Cabozantinib, or Lenvatinib induce a marked decrease in fin melanogenesis. Each grey dot in J represents an individual zebrafish, and the table underneath the summarizes each inhibitor’s potency against VEGFRs or FGFRs (+potent against only one FGFR or VEGFR protein, ++ potent against all FGFR or VEGFR proteins, - not potent). **(E)** Tailfin melanocyte cell area quantification at different drug doses in mitfa-CRKL MC-GFP zebrafish. Drug abbreviations: Apa – Apatinib, Anlo – Anlotinib, Lenv – Lenvatinib, Cabo – Cabozantinib, Suni - Sunitinib.

These inhibitors were evaluated for their ability to reduce acral-like fin melanogenesis in a premalignant *mitfa*-CRKL MC-GFP zebrafish embryo model described previously^6^ (**Figure 2B**). The embryos were treated with RTK inhibitors for 48 hours prior to brightfield and fluorescent imaging to quantify GFP+ fin melanocytes. While all inhibitors exhibited a modest to profound dose-dependent effect on tailfin melanocyte cell area, the most potent inhibitors were Anlotinib, Cabozantinib, and Lenvatinib (**Figure 2C-E**, **Figure S1A**). Similar results were seen in the wild-type MC-GFP control embryos with Anlotinib and Lenvatinib in addition to a general decrease of GFP area throughout the animal, indicating that the compounds also act as general inhibitors of normal melanogenesis in a dose-dependent manner (**Figure S1B-C**). Compound toxicity was assessed by evaluating zebrafish embryo morphology, length, and yolk sac area, and no significant toxicity was identified using 1µM and 3µM of the inhibitors (**Figure S1D**). By comparing the IC_50_ differences between drugs, it appears that melanogenesis can be disrupted by either direct FGFR/VEGFR inhibition with Anlotinib or Lenvatinib as well as through a combination of AURKB/AXL/MET/FLT3/TIE2/VEGFR inhibition with Cabozantinib. Based on these results, we elected to compare two of the FDA-approved agents in our preclinical PDX tumor models: Lenvatinib as a potentially efficacious dual FGFR/VEGFR therapeutic and Sunitinib as a VEGFR inhibitor control with modest effects on fish melanogenesis.

### Characterization of AM patient-derived xenograft (PDX) mouse tumor models

Lenvatinib and Sunitinib were tested against a panel of AM and CM PDX tumors. These tumor models were selected to represent a genetically diverse population of AM tumors that represent all TCGA genetic groups and a comparatively diverse complement of CM tumors. De-identified clinical information, pathological stage at presentation, patient treatment and survival, and details about tissue type used to generate PDX models are summarized in **Table 1** and detailed in **Table S1**. In total, five CM and nine AM patients were enrolled with a total of 17 PDX tumors generated. In general, the cohort mirrored clinical reports that demonstrate a propensity for AM to be identified at higher stages and to have worse patient outcomes^41–45^. The type of tissue used to generate AM PDX models included untreated primary specimens (46%) and regional/distant metastasis (54%). Comparatively, all CM PDX models were generated on metastatic tissue with most being previously treated with biologics or chemotherapies (67%). The site of primary disease was primarily in the upper body for CM tumors. For acral disease, both acral subungual melanoma (ASM) and melanomas at acral volar sites (AM) were collected at sites on the hands and feet (**Figure 3A**). The primary acral tumor sites were confirmed by chart review, review of pathologic records, and, where available, clinical photos of pre-biopsy and/or resected lesions. Two AM required additional effort to ensure correct classification. HCI-AM092 was a nodular lesion present on the ankle at the junction between glabrous and non-glabrous skin. Histologic review confirmed the presence of acral lentiginous melanoma in-situ radiating out from the main tumor on the glabrous skin portion of the specimen, confirming the diagnosis of AM. HCI-AM090 was not originally biopsied at the University of Utah, and no prebiopsy clinical photos were available. Pre-resection documentation and photos detail a tumor encompassing the dorsal skin and nail bed of the 4^th^ toe. Anecdotally, this lesion first arose on the dorsal skin, but we could not confirm this evidence with a high degree of certainty. Thus, HCI-AM090 was designated as an AM with the potential that it represents ASM or a rare triple-wild type CM.

**Figure 3:**
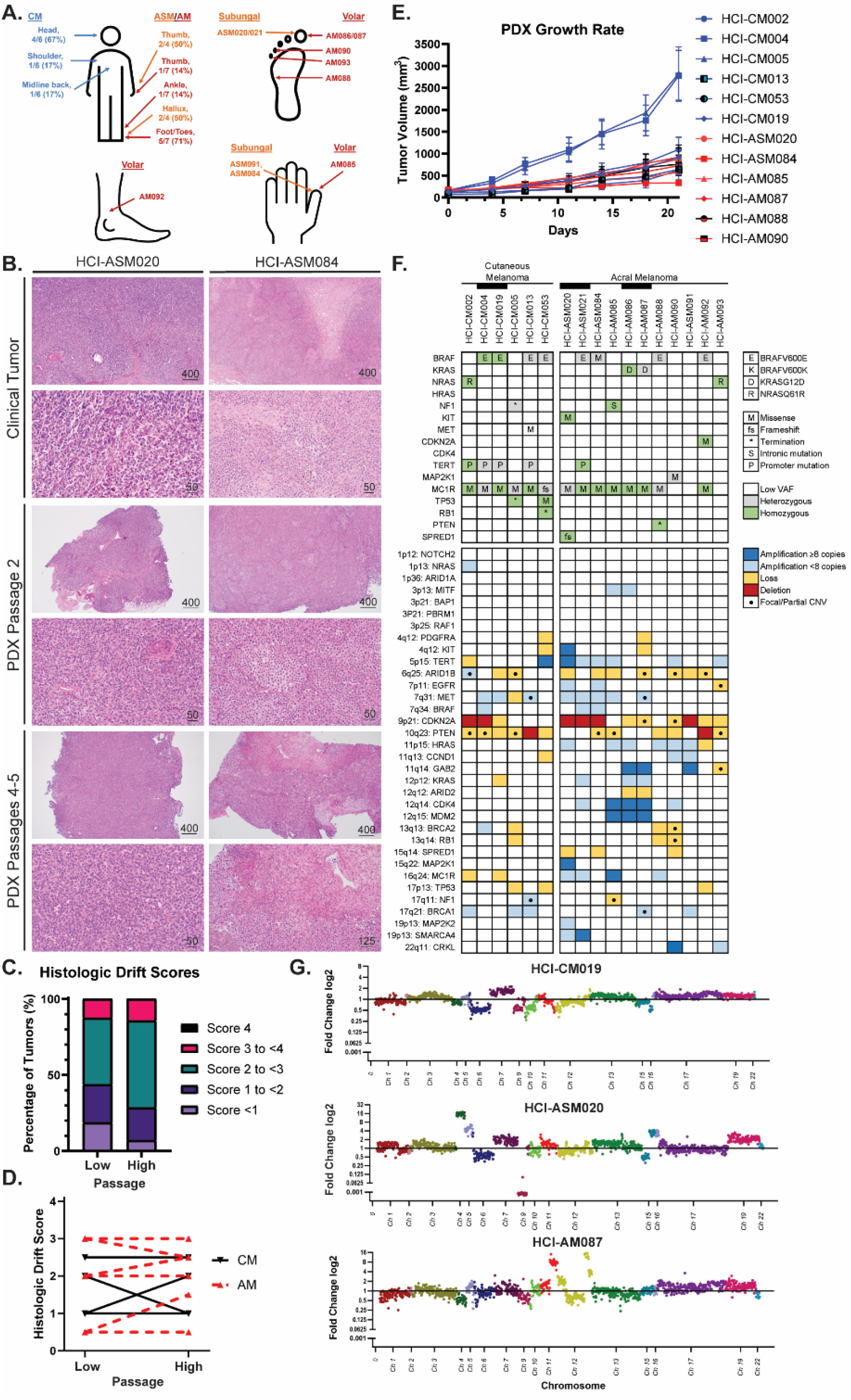
Histologic and genetic characterization of AM and CM PDX tumor models. **(A)** Anatomic location of AM and CM tumors. CM are labelled in blue text, acral subungal melanoma (ASM) in orange, and volar AM in red. **(B)** Representative clinical, low-passage, and high passage PDX tumor histology images. **(C-D)** Results from the Clinical Drift Score validation indicates that PDX tumors are histologically stable through multiple passages. Scores represent cytologic and histologic architecture drift from the original clinical tumor histology: none/identical (0), minimal (1), mild (2), moderate (3), and marked (4). Definitions and criteria for each classification are available in Supplemental Data 3. Four CM tumors and nine AM tumors are plotted in C-D. **(E)** Growth rates of representative PDX tumors. **(F)** Available AM PDX models encompass the spectrum of TCGA MAPK mutations. Relevant Tier 1 (pathogenic) and 2 (likely pathogenic) SNV mutations, MC1R mutations associated with melanoma predisposition, and CNV are represented in the tile plot. Black bars link PDX tumors that were collected from the same patient. **(G)** AM PDX models with localized highly amplified genomic regions are compared to a representative CM model.

**Table 1.** Overall clinical characteristics of patients and PDX models.

| **Table 1. Overall clinical characteristics of patients and PDX models** | **All Patients**<br>**(n = 14)** | **CM**<br>**(n = 5)** | **AM**<br>**(n = 9)** |
| --- | --- | --- | --- |
| **Age** |  |  |  |
| Median (range), years | 59<br>(24-85) | 62<br>(24-85) | 59<br>(53-83) |
| 65 and older, number (%) | 4 (29%) | 1 (20%) | 3 (33%) |
| **Gender** |  |  |  |
| Male | 8 (57%) | 3 (60%) | 5 (56%) |
| Female | 6 (43%) | 2 (40%) | 4 (44%) |
| **Patient Status (as of 3/2024)** |  |  |  |
| Alive | 8 (57%) | 4 (80%) | 4 (44%) |
| Dead | 6 (43%) | 1 (20%) | 5 (56%) |
| **Clinical Stage at Presentation (TNM)** |  |  |  |
| IA/B | 3 (21%) | 3 (60%) | 0 |
| IIA/B/C | 4 (29%) | 2 (40%) | 2 (22%) |
| IIIA/B/C/D | 7 (50%) | 0 | 7 (78%) |
| IV | 0 | 0 | 0 |
| **Total number of PDX collected** | 17 | 6 | 11 |
| Paired primary and metastasis | 1 (6%) | 0 | 1 (9%) |
| Paired regional and distant metastasis | 1 (6%) | 1 (20%) | 0 |
| Paired untreated and treated | 1 (6%) | 0 | 1 (9%) |
| **Tissue type for PDX tumor** |  |  |  |
| Primary tumor | 5 (29%) | 0 | 5 (46%) |
| Local Recurrence | 0 | 0 | 0 |
| Local Metastasis | 2 (12%) | 2 (33%) | 0 |
| Regional Metastasis | 5 (29%) | 2 (33%) | 3 (27%) |
| Distant Metastasis | 5 (29%) | 2 (33%) | 3 (27%) |
| **Treatment (biologicals, chemo-therapy, and/or radiation)** |  |  |  |
| Untreated prior to PDX collection | 6 (35%) | 1 (17%) | 5 (45%) |
| Treated prior to PDX collection | 9 (53%) | 4 (67%) | 5 (45%) |
| Treated after PDX collection | 12 (71%) | 3 (50%) | 9 (82%) |
| Unknown treatment history | 2 (14%) | 1 (17%) | 1 (9%) |

All PDX tumors were histologically reviewed by a pathologist to confirm clinical diagnosis and to assess if passaged PDX tumors continued to represent their clinical counterparts (**Figure 3B**, Supplemental Data 2 and 3). During longitudinal histologic review, a qualitative tool was designed to compare the histologic drift of PDX tumors from the original clinical tumor. A detailed breakdown of the score criteria is listed in Supplemental Data 3 with a summary in the methods section. After developing the tool, two independent pathologists with a specialization in cytology reviewed and scored low passage (passage 1 and 2) and high passage (passage 3-5) PDX tumors for tool validation (**Figure 3C-D**, Supplemental Data 3). The validation cohort had a 92.9% concordance rate for calls within +/- 1 score variation at low PDX passage, and 92.3% at high PDX passage. Breakdown of the average histologic drift scores between high and low passages from the validation cohort reveals an overall decrease in 0-1 score tumors in the high passage tumor group, with a corresponding increase in the 2-3 score categories. More than half of the tumors maintained their low passage score or decreased at higher passages (8/13, 62%) with the remaining increasing in score at higher passages (5/13, 38%). No tumor increased greater than 1 score between low and high passages. The stability of histologic appearance mirrors a previous report where clinical tumor CNV was stably maintained in passaged PDX tumors^46^.

The growth rates of most acral and cutaneous PDX tumors are similar (**Figure 3E**), with the notable exceptions of HCI-CM004 and HCI-CM005.

Genomic characterization was performed by targeted gene sequencing, with all pathogenic (Tier I) and likely pathogenic (Tier II) point mutations, as classified by the 2017 AMP/ASCO/CAP criteria^30^, shown in **Figure 3F** and Supplemental Data 4. The only variants of uncertain significance (Tier III mutations) presented in these figures are high-incidence MC1R variants that have been proposed as risk modifiers for developing CM^47^. The driver mutations in six CM models include two homozygous BRAFV600E, two heterozygous BRAFV600E, one NRASQ61R, and a NF1 termination mutation. TERT promoter mutations were common in 4/6 of the models, and all CM models had MC1R Tier III variants. Comparatively, the AM models had a more diverse range of RAS pathway driver mutations, including two TCGA ‘triple wild-type’ tumors that lacked driver mutations common to CM. Driver mutations in the other nine models include three heterozygous BRAFV600E mutations, one BRAFG469A, two KRASG12D from the same patient, one NRASQ61R, one hemizygous NF1 intronic mutation producing a pathogenic splicing variant, and a homozygous amplified KIT mutant. Pathogenic TERT promoter mutations were very rare (1/11, 9%). Per published reports, AM typically have greater numbers of copy number variation (CNV) events compared to CM, which was also seen in our cohort (**Figure 3F-G**, Supplemental Figure 2, Supplemental Data 5). These include highly amplified genes (≥8 copy number) at loci previously observed to be impacted by tyfonas/hailstorm events^18,19^ in 7/11 (64%) of AM compared to only one CM tumor (17%).

A summary of each PDX tumor histology, clinical history, and genetics are available in Supplemental Data 6: PDX Model Data Sheets.

### Dual FGFR/VEGFR inhibition induces growth arrest or regression in all tested AM PDX models

The effects of broad-spectrum RTK inhibition with Sunitinib and targeted dual FGFR/VEGFR inhibition with Lenvatinib on tumor growth kinetics were compared using our well-charactered PDX tumor models. In order to plot all tumor models on the same graph, individual tumor plots were Log transformed and the slope of growth (‘Tumor Growth Velocity’) for each tumor was determined according to published Rate-based T/C methods^48^ (more details in **Figure S3A**). Each dot in **Figure 4A** and **4C** represents the slope of tumor size change for an individual tumor. Sunitinib treatment over 21 days significantly slowed tumor growth rate in half of the tested CM and AM models, and inhibited tumor growth in one of four AM. Overall, there was no significant change in growth rates between the CM tumors and all AM with Sunitinib (**Figure 4A-B**, **Figure S3B**). Over similar timescales, Lenvatinib also slowed CM tumor growth rates, with one of five tested CMs achieving net-zero tumor growth. Comparatively, Lenvatinib was more efficacious in AM: with an oncostatic response in two tumors and tumor regression in the remaining four AM tumors (**Figure 4C-F**, p<0.0001, two-way ANOVA).

**Figure 4:**
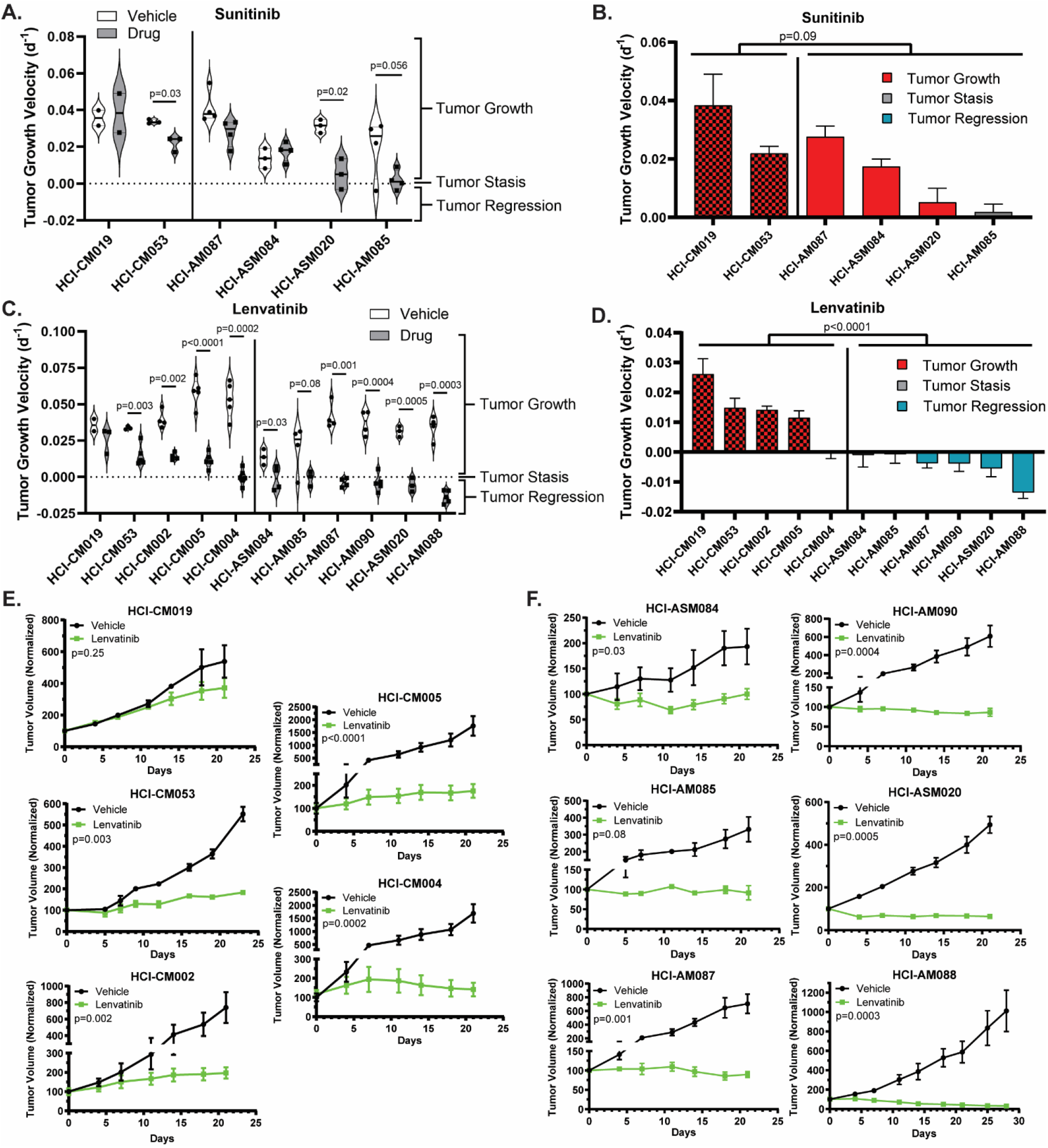
Dual FGFR/VEGFR inhibition with Lenvatinib induces tumor stasis or regression in all AM PDX tumors. Tumor growth velocity for individual Sunitinib (daily, 40mg/kg) and Lenvatinib (daily, 50mg/kg) treated PDX tumors are represented as violin plots **(A, C)**. The Lenvatinib-treated tumors from (A, C) are shown as bar plots that are color-coded based on depth of response **(B, D).** CM tumors are indicated by a checkerboard pattern. Average tumor growth of vehicle and Lenvatinib treated CM **(E)** and AM **(F)** are shown for each PDX model. Two-way ANOVA was used to determine differences between CM and AM therapy response (B-D), and student T-tests were used to compare vehicle and treatment differences within a PDX model (A, C, E-F).

### AM PDX models had variable responses to rationally selected non-FGFR/VEGFR inhibitors

In parallel with the RTK inhibitor drug study, several other narrow-spectrum kinase or enzyme inhibitors were screened based on promising literature findings. Both melanocytes and melanoma have high EGFR phosphorylation activity^49^, which makes it a promising target for EGFR inhibitors such Dacomitinib. While there was a statistically significant change in tumor growth velocity in one of the AM models, stable disease or regression was not achieved (**Figure S3C**). ERBB2/HER2 is another potential target as it is amplified or mutated in 5% of AM by MSK-IMPACT NGS testing^50^ and is the most common dimerization partner of ERBB3^49^. Despite ERBB3 being identified as a marker of poor prognosis in melanoma^51^ and enriched in volar melanocyte transcriptomes^27^, no antitumoral activity was identified with the HER2 inhibitor Lapatinib (**Figure S3D**). TRKB/NTRK2 was similarly enriched in volar melanocytes^27^ and can be activated by EGF or neurotrophin^52^. While the ANA12 antidepressant is a specific NTRK2 inhibitor^53^, it had no anti-tumoral activity on our screening AM cohort (**Figure S3E**).

Our interest in studying Olaparib, a PARP inhibitor that induces single-strand DNA breaks which are synthetic lethal in the presence of homologous recombination repair deficiency, stems from early promising results in CM studies^54^ and the observation of tyfonas^18^ and hailstorms^19^ in AM, which imply the presence of genomic instability. Unfortunately, all tumors grew on Olaparib therapy and there was no synergy with the addition of other inhibitors (**Figure S3F-H**). This lack of efficacy mirrors a recent case study wherein a patient with a BRCA1-mutated AM achieved stable disease with Olaparib treatment for only four months^55^. A second synthetic lethal approach where the complementary non-homologous end-joining DNA repair pathway was inhibited with NU7441 was also ineffective (**Figure S3I**).

### Dual FGFR/VEGFR inhibition is not directly cytotoxic to transiently cultured AM PDX tumor cells

Out of the tested therapeutics, Lenvatinib was the most effective in inducing oncostasis or tumor regression in the AM PDX models, but, in light of the poor performance of Lenvatinib in CM in the LEAP-003 study, we sought to determine the mechanism by which AM specifically respond to the drug since it could direct second-generation targeted AM drug development. Lenvatinib has multiple potential mechanisms of action described in the literature: inhibition of angiogenesis^20,56–58^, induction of ferroptosis^59,60^, inhibition of the cell cycle^61^, and stimulation of the adaptive immune response^62–64^. Since the adaptive immune system is absent in our PDX tumor models, we sought to delineate which of the other processes could have provided the robust tumor response.

AM researchers have anecdotally stated the difficulty in establishing AM cell lines^3^, and we have confirmed this challenging reality as only one of five AM PDX tumors continued to propagate in long-term cell culture versus all three attempted CM PDX tumors (**Figure 5A**). To circumvent this challenge, fresh PDX tumors were harvested, dissociated into individual cells, and depleted of mouse stroma before plating onto a 96 well plate for 72 hours of quantitative phase imaging (QPI) with RTK inhibitors (**Figure 5B**). This strategy was chosen as 48 hours of monitoring growth rate has been shown to provide equivalent endpoint data to traditional metabolically-activated dyes such as Cell-Titer-Glo while also 1) measuring the kinetics of cell growth and 2) capturing cell images in real-time^35^. Considering that a subset of cultured AM PDX cells survive only a few days in culture (HCI-AM088 and HCI-AM090), we opted to use QPI to evaluate changes in growth kinetics and cell mass over a short time period. For each drug, a dose titration was performed and individual cell specific growth rate (SGR, **Figure S4A-C**) and whole image ‘normalized mass’ were calculated (**Figures 5C-F**). Changes in population size and distribution of these characteristics are followed over time (**Figure S4C**) to derive GR_50_ curves and values (**Figure S5A-B**).

**Figure 5:**
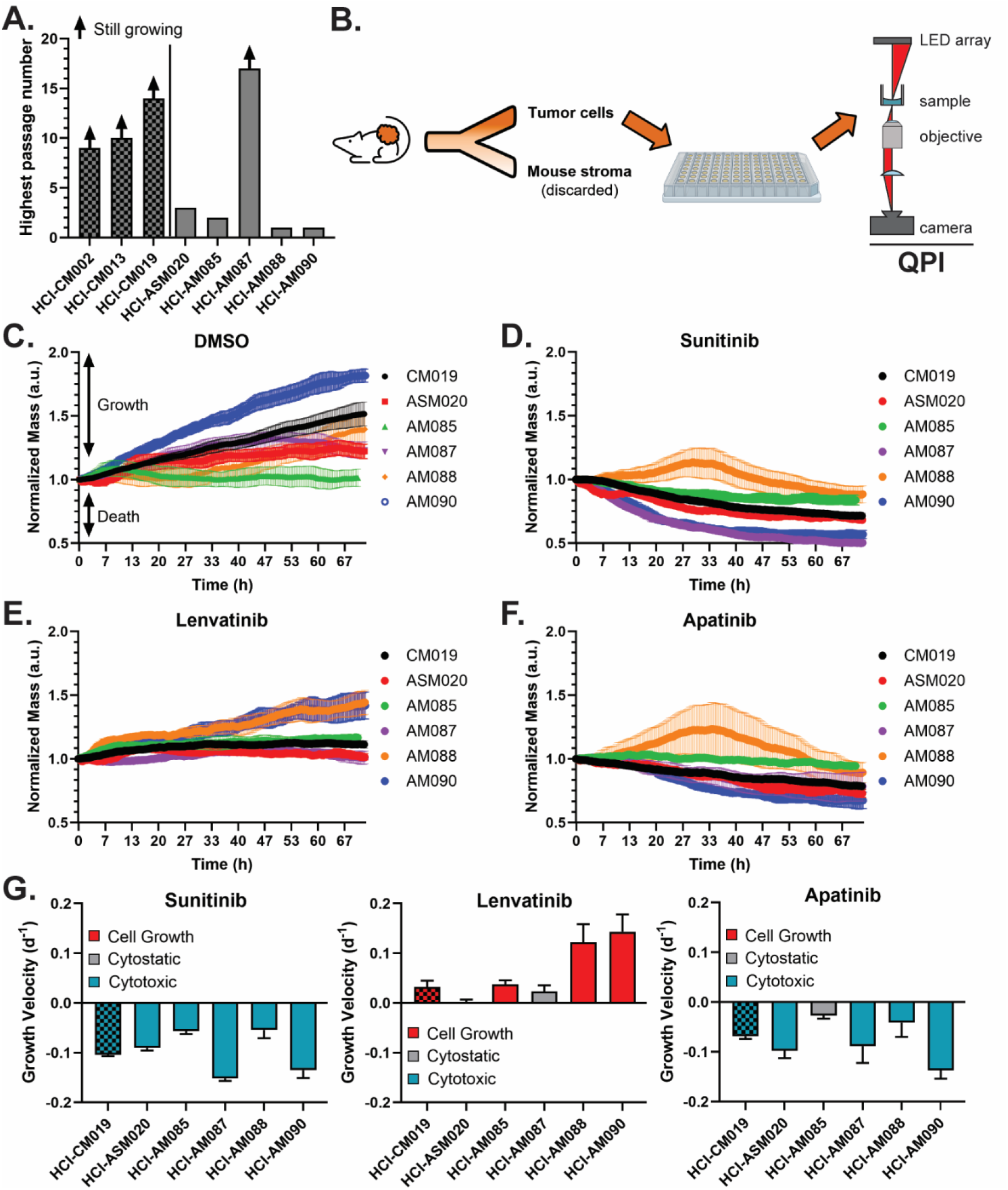
Dual FGFR/VEGFR is not directly cytotoxic in dissociated AM PDX tumor cells. **(A)** Most AM cells cultured from PDX tumors do not grow in extended culture conditions. **(B)** Fresh PDX tumors were collected, dissociated into individual cells, and depleted of mouse stroma before plating for immediate and short term culture drug studies using quantitative phase imaging (QPI). Growth of AM and CM cells in **(C)** 0.5% DMSO; **(D)** 50µM Sunitinib, **(E)**, 20µM Lenvatinib, and **(F)** 40µM Apatinib. **(G)** To facilitate comparison to PDX tumor growth velocity in Figures 4B and 4D, growth velocity was calculated by linear regression of normalized mass.

Under 0.5% DMSO control conditions, all dissociated cells grew except for HCI-AM085 (**Figure 5C**). While no tumor regression was noted with Sunitinib with *in vivo* PDX-tumor studies, 50µM Sunitinib uniformly induced cell death in all transient cell cultures (**Figure 5D, 5G**). This observation indicates that Sunitinib has direct cytotoxic and/or cytostatic activity against melanoma cells in culture, but *in vivo* tumors are able to evade this method of cell killing. Conversely, the *in vivo* efficacy that we observed with Lenvatinib against AM was not observed in transient tissue culture. Three AM cultures grew despite 20µM Lenvatinib and the remaining two experienced no net cell growth or death, indicating that Lenvatinib is at best directly cytostatic and may be indirectly cytotoxic *in vivo* (**Figure 5E, 5G**). To determine if the cytostatic response was secondary to FGFR inhibition vs VEGFR-specific inhibition, Apatanib was evaluated against these cells (**Figure 5F**). At 40µM of Apatanib, one CM and four AM cultures demonstrated direct cytotoxicity, and one AM culture was cytostatic. This degree of response was unexpected given the intermediate effects of Apatanib on zebrafish melanogenesis (**Figure 2C-E**) and the drug’s ultra-narrow specificity for VEGFR2 and RET. The observed efficacy was likely due to the significant and very high RET expression in AM compared to CM (**Figure 1F**) and indicates that RET/VEGFR dual inhibition should be considered for future preclinical studies.

While GR_50_ could be calculated for most cells and conditions, this was not possible for conditions in which the melanoma cells had at least transient growth, such as Lenvatinib in HCI-AM085/088/090 cultures and Sunitinib against HCI-AM088 (**Figure S5A-B**). In general, Apatanib tended to have a higher GR_50_ when compared to the other two drugs but had at least a transient effect on growth for every melanoma culture. When Lenvatinib inhibited cell growth, it was often at a lower concentration than the other agents.

### Tumor vasculature decreases with dual FGFR/VEGFR inhibition

Based on the *in vivo* tumor regression (**Figure 4**) and the *in vitro* QPI experiments (**Figure 5**) we hypothesized that Lenvatinib has an indirect cytotoxic effect on *in vivo* tumor cells. In an immunocompromised setting, Lenvatinib’s main extrinsic effect should be disruption of tumor vasculature^20,56–58^. To test this hypothesis, assessment of tumor vasculature, proliferation, necrosis, and ferroptosis was conducted on FFPE tumors collected after 21 days of therapy. Most of the AM PDX tumors had a complete response by gross exam and could not be collected for FFPE. The two exceptions were HCI-ASM084, which had an oncostatic response, and HCI-ASM087, which partially regressed with treatment. These tumors were compared to HCI-CM004, which also had an oncostatic response, and HCI-CM005, which had the slowest growth rate in the CM cohort on Lenvatinib therapy (**Figure 3C-D**). A pathologist measured the percentage of necrosis within the tumors on H&E staining, percentage of proliferating cells by Ki67 IHC staining, ferroptosis score by multiplying the average membranous stain intensity and percentage of tumors cells with TfR1 IHC membranous staining, and the number of CD31 IHC positive blood vessels per 10x field (**Figure 6A**). While the CM tumors had a significantly lower number of proliferating cells following Lenvatinib treatment, the effect of treatment on the replication rate in the AM models was moderate, variable, and statistically insignificant (**Figure 6B**). No significant difference was observed in tumor necrosis (**Figure 6C**) or ferroptosis (**Figure 6D**) for any Lenvatinib-treated tumors. Comparatively, a consistent, dramatic, and biologically significant decrease in CD31+ blood vessel density was observed across all PDX tumor models (**Figure 6E** and **Figure S6**) with a qualitative decrease in lumen diameter and wrapping around tumor cell nests. These blood vessel changes provide a mechanism for how FGFR/VEGFR dual inhibition can lead to tumor regression or stasis in AM when cell culture-based proliferation assays demonstrate no difference (**Figure 5**).

**Figure 6:**
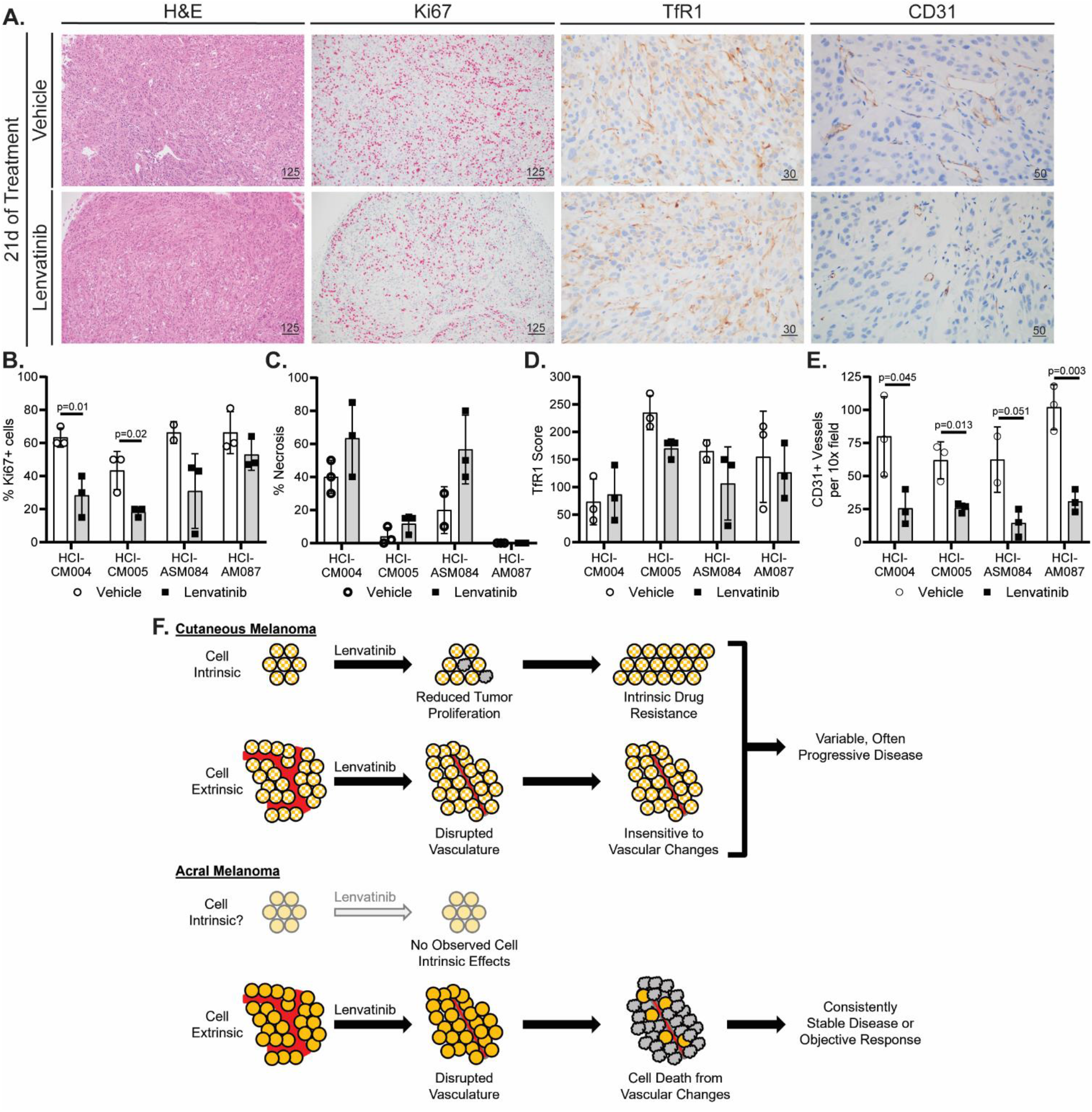
Lenvatinib halts AM tumor growth or induces regression by remodeling tumor vasculature. **(A)** Representative histology (H&E, 10x objective) and immunohistochemistry (IHC) images of vehicle and Lenvatinib HCI-AM087 are shown. CD31 (20x objective) and transferrin receptor 1 (TfR1, 40x objective) were stained with DAB, and MiB/Ki67 (10x objective) was stained with red chromogen. Scale bars are in microns. A pathologist quantified the **(B)** Ki67 IHC positive cell percentage, **(C)** percent necrosis from H&E-stained sections, **(D)** membranous TfR1 IHC scores, and **(E)** CD31 positive IHC vessels. Significance was determined using the Student’s t-test. **(F)** While CM has reduced tumor proliferation and diminished blood vessel quantity and quality on Lenvatinib therapy, these PDX tumors often continue to grow. There are no observed direct cytotoxic effects of Lenvatinib on AM cells. Instead, tumor regression or stable disease is achieved in AM tumors by reducing the blood vessel quantity and quality.

## DISCUSSION AND CONCLUSIONS

Our data indicate that AM is highly reliant on tumor vasculature for survival and growth, and inhibition via Lenvatinib is sufficient to halt growth or induce regression. Comparatively, while CM suffers both the intrinsic effects of decreased cell proliferation and extrinsic effects of decreased vascularization with Lenvatinib, it appears less sensitive to the extrinsic effects and is capable of overcoming the intrinsic barriers to growth (**Figure 6F**). The amplification of genes in the FGF/FGFR/CRKL/GAB2 signaling axis in AM has previously been described^6,19,65^, and our work presents the first attempt to drug this pathway in preclinical models. The therapeutic efficacy of Lenvatinib across all tested AM models reveals an important observation: amplification of genes in the FGF signaling pathway is not a pre-requisite for Lenvatinib sensitivity. For instance, HCI-AM088 had the greatest response to Lenvatinib (**Figure 4C-D**) despite having no amplification of the CRKL, GAB2, or CCND1 loci (the latter contains the FGF3, FGF4, and FGF19 genes), and HCI-ASM084 and HCI-AM085 experienced stable disease without amplifications at these loci (**Figure 3F**). These data combined with the general suppression of fin melanogenesis in zebrafish (**Figure 2** and **S1**) indicate that dual FGFR/VEGFR inhibition targets an essential signaling axis in acral-type melanocytes and AM.

Due to the difficulty of establishing stable AM cell lines^3^ (**Figure 5A**), animal and PDX models were leveraged to screen potential drug candidates. This *in-vivo* approach to drug discovery provided the benefit of incorporating drug effects on tumor cells, stroma, and vasculature within the context of a living organism. The degree of incongruence observed between PDX tumor treatment and transient *ex-vivo* cell culture was remarkable considering that Lenvatinib was the most efficacious agent against AM PDX tumors but was either oncostatic or ineffective against dissociated tumor cells. Since mouse stroma is present within the PDX tumors, it is possible that these findings may not translate to human tumor neovasculature, and it is not possible to assess the potentiation of ICIs by Lenvatinib in PDX models due to lack of an adaptive immune system. Conversely, Sunitinib would have been identified as a promising therapeutic agent *ex-vivo* due to intrinsic cytotoxicity, but would have failed *in-vivo* testing in preclinical PDX tumors. While direct anti-melanoma cell cytotoxic responses in 2D culture often translate to PDX tumor response^66^, drugs that target supporting stroma cells and vasculature, such as Lenvatinib, are indirectly cytotoxic and are not expected to provide a response in a pure tumor cell culture. Loss of genetic heterogeneity, genetic drift, and epigenetic changes from culture conditions and media are appropriate concerns for long-term melanoma cell culture but, like primary patient cell cultures, are not expected to impact transient 2D culture from PDX tumors^35,66,67^. The discrepancies observed between *in-vivo* and *ex-vivo* model systems highlights the importance of considering potential indirect or stroma/vasculature-targeted cytotoxic effects of drugs when deciding on a drug screening method.

From a translational standpoint, Lenvatinib has been FDA approved for hepatocellular carcinoma^68^, renal cell carcinoma^69,70^, endometrial carcinoma^71^, and iodine-resistant differentiated thyroid carcinomas^72^. It has also been evaluated in combination with immune checkpoint inhibitors (ICI) for melanoma in the LEAP-003 (presented at the 20^th^ Society for Melanoma Research Congress) and LEAP-004 studies^73^. While these studied showed no survival benefit with Lenvatinib across all skin melanomas, it is important to highlight that acral melanomas were not considered a separate melanoma subtype in these studies. Based on a small cohort study where 4 of 6 acral melanomas responded to second-line Lenvatinib plus ICI^74^ and an early clinical study where Anlotinib, another dual FGFR/VEGFR inhibitor, provided a significant survival benefit when used with ICIs^75^, targeting AM with FGFR/VEGFR inhibitors are expected to provide a survival benefit for patients. Based on these studies and our promising preclinical data, we recommend conducting a case series where patients with AM are treated with ICI and Lenvatinib, or another regionally available dual FGFR/VEGFR inhibitor, as a second- or third-line therapy to further establish clinical benefit.

Due to the genetic instability of melanoma, use of PARP inhibitors such as Olaparib have been investigated as a potential synthetic lethal therapy approach^54,76,77^. These inhibitors have been successfully used in homologous-recombination deficient cancers that include breast, ovarian, and prostate carcinomas, and, while preclinical data was promising for the use of Olaparib in melanoma, a case study of a patient with BRCA1-deficient metastatic AM derived 4 months of stable disease on Olaparib^55^. While beneficial for the patient, it is important to note that BRCA deficiency often provides a longer progression-free survival in breast^78^, ovarian^79^, and prostate^80,81^ cancers. Further reinforcing concerns that AM has a reduced response to PARP inhibitors despite genomic instability, no AM PDX models achieved regression or stable disease despite Olaparib dose-escalation and combination therapy regimens (**Figures S3F-H**).

While PDX models are well positioned to aid researchers in identifying potential disease modifying genes and/or phenotypes, the race and ethnicity of patients enrolled in Western^13,14,16^ and Chinese^12,15^ AM PDX cohorts were not described. Towards the goal of developing an international and multiracial cohort of PDX tumors, we are contributing 11 clinically, histologically, and genetically characterized AM PDX from self-identified white non-Hispanic patients. However, to accurately capture the heterogeneity of AM in preclinical testing, there is a critical need for additional AM models generated from underrepresented groups, such as Africans, Hispanic and Latino/a/x, and Indigenous populations. Since individuals from these groups experience worse outcomes relative to other groups^42,82^, understanding the biology of AM in these contexts is essential and represents an underserved societal need.

In summary, we have demonstrated an efficient pipeline to identify potential therapeutics by 1) using available informatic datasets, 2) screening agents in zebrafish, 3) confirming anti-tumoral activity in a preclinical PDX model, and 4) assessing mechanism of action using transient cell culture and histology techniques. With this method, we identified Lenvatinib as a promising clinical agent against AM, and, based on available clinical evidence, encourage clinician to consider using this, or a regionally available dual FGFR/VEGFR inhibitor, for the treatment of acral melanoma.

## AUTHOR RESPONSIBILITIES

Hypothesis generation was performed by EAS, RLB, HZ, and RLJ-T. Project leadership was spearheaded by EAS, RLB, and RLJ-T. Writing, figure generation, literature informatics analysis, histology review, IHC evaluation, and slide imaging were conducted by EAS. Patient chart review was performed by EAS and DCD. Zebrafish studies were designed and performed by NMC and RMW. Histologic clinical drift scoring was developed by EAS and validated by MB and TAS. DNA library preparation and sequencing were performed by CAB and AJ, and genomic analysis was conducted by EAS, CAB, AJ, and DCD. PDX tumor development, passaging, drug testing, and data analysis was a collaborative effort by EAS, RLB, JRH, AHG, DHL, TYC, RLJ-T and the Preclinical Research Resource core at the Huntsman Cancer Institute. Quantitative phase imaging of dissociated PDX tumor cells was a collaborative effort between EAS, DHL, ECS, TEM, SA, TAZ, and the Preclinical Research Resource core. Immunohistochemistry development was conducted by EAS and the research BMP IHC core at the Huntsman Cancer Institute. Funding for this project was secured by EAS, TAZ and RLJ-T. EAS drafted the original manuscript and EAS, RLB, DCD, RMW, TAZ, and RLJ-T substantially contributed to edits.

## ACKNOWLEDGMENTS

This work was supported by a National Cancer Institute R01 (R01CA276653) to RLJ-T and TAZ, the Harry J Lloyde Charitable Trust Melanoma Research Grant to RLJ-T, the 5 for the Fight Fellowship to RLJ-T, the ARUP Research Histology Lab Pathology Research Support Fund to EAS and pilot funds from the Huntsman Cancer Institute Cell Response and Regulation Program and Melanoma Center. We utilized the Shared Resources for Research Informatics, High-Throughput Genomics, Preclinical Research Resource, Immuno Oncology Network Core, and Biorepository and Molecular Pathology at Huntsman Cancer Institute at the University of Utah supported by the National Cancer Institute of the National Institutes of Health under Award Number P30CA042014. The content is solely the responsibility of the authors and does not necessarily represent the official views of the NIH.

## SUPPLEMENTAL FIGURE LEGENDS

**Supplemental Figure 1:**
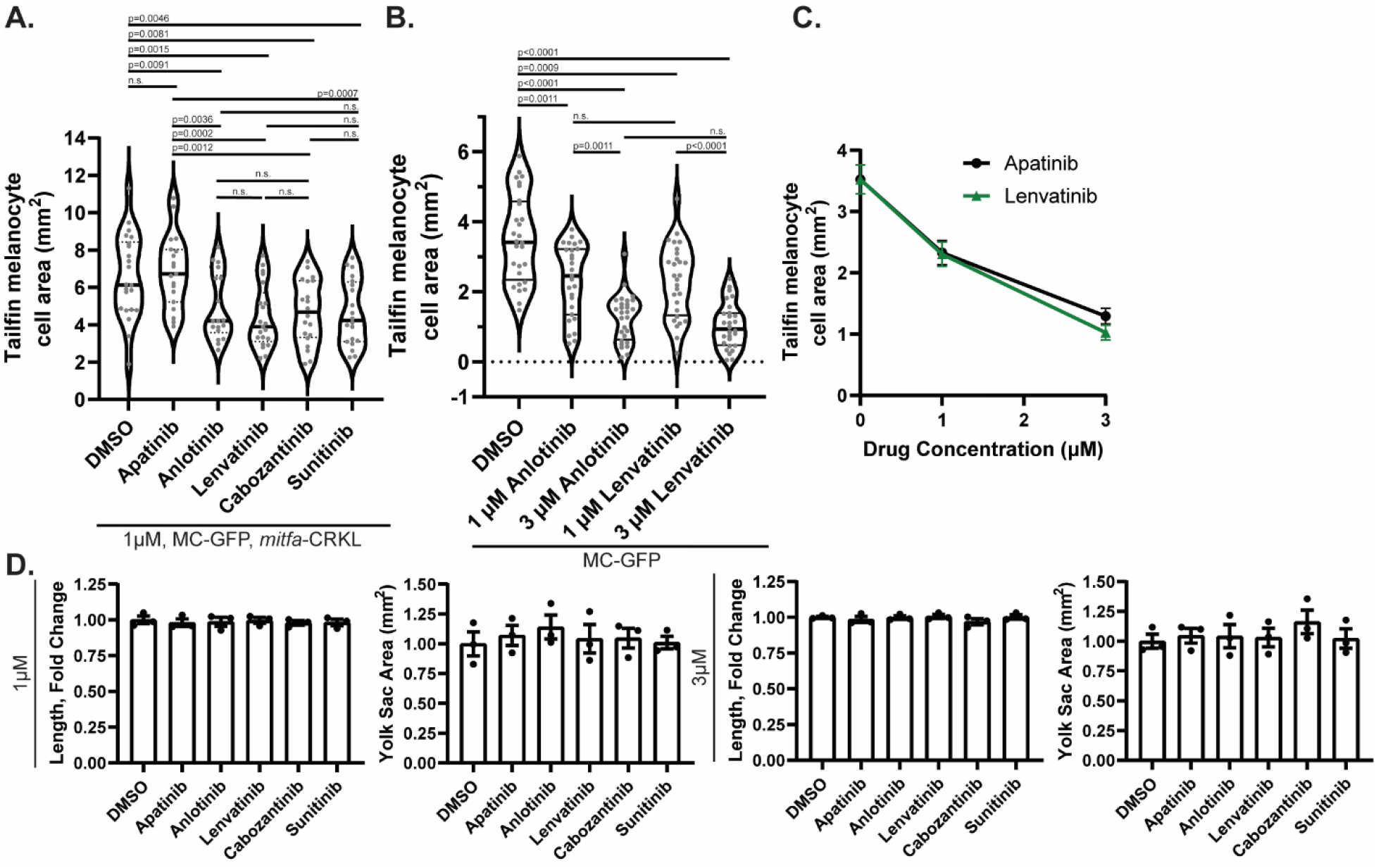
Multi-RTK inhibitors prevent melanogenesis in wild-type and mitfa-CRKL premalignant zebrafish in a dose-dependent manner without organism-level toxicity. **(A)** Tailfin melanocyte cell area at 1µM dose of multi-RTK inhibitors in the mitfa-CRKL MC-GFP model. **(B-C)** Tailfin melanocyte cell area with treatment of Anlotinib and Lenvatinib at 1 and 3µM in CRKL wild-type MC-GFP cells. **(D)** Organism-level toxicity evaluation by length and yolk-sac area for each multi-RTK inhibitor in mitfa-CRKL MC-GFP zebrafish.

**Supplemental Figure 2:**
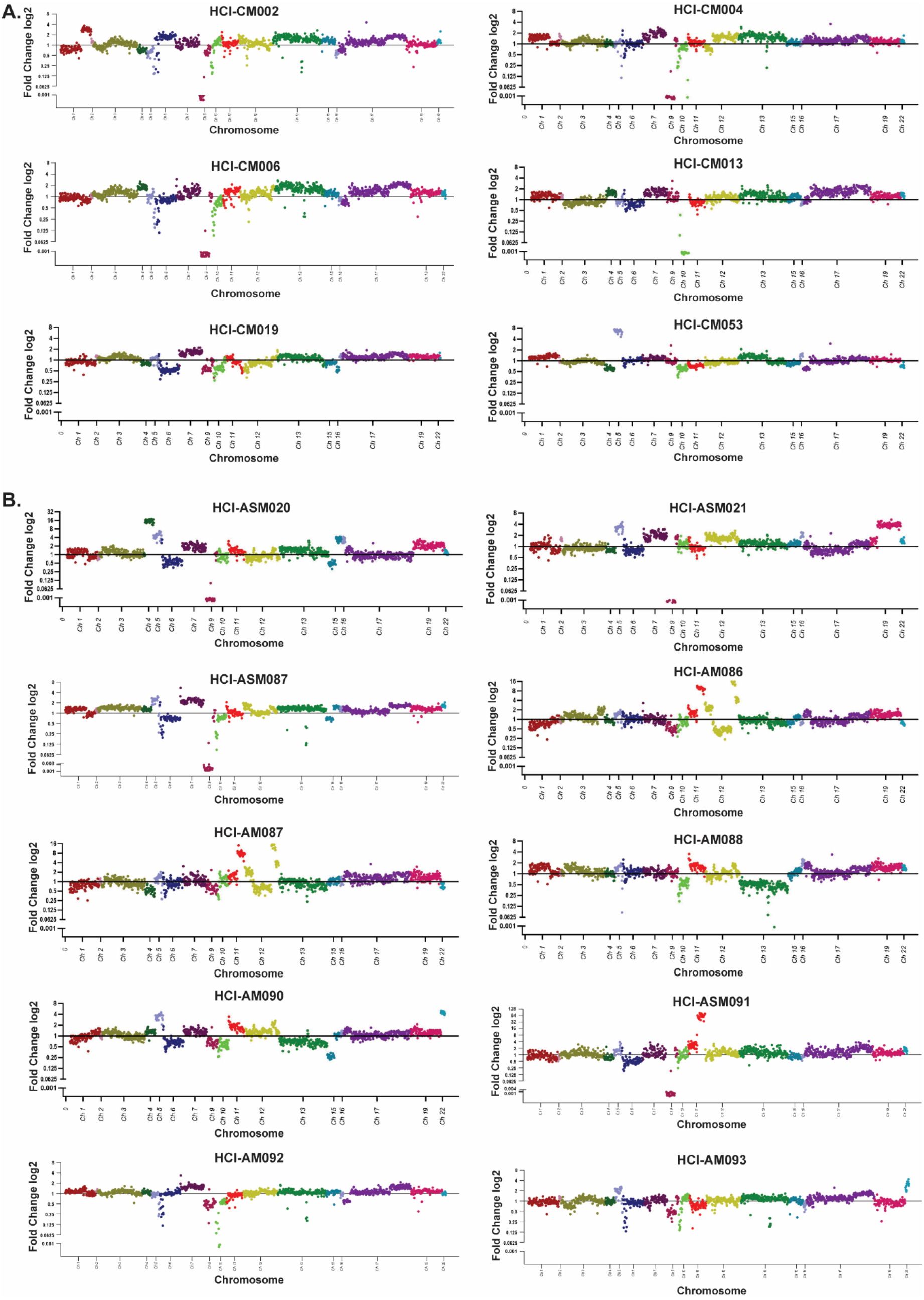
Raindrop plots representing CNV for each PDX model. **(A)** CM CNV plots and **(B)** AM CNV plots. Paired PDX models developed from the same patient include HCI-CM004 and HCI-CM019, HCI-ASM020 and HCI-ASM021, and HCI-AM086 and HCI-AM087.

**Supplemental Figure 3:**
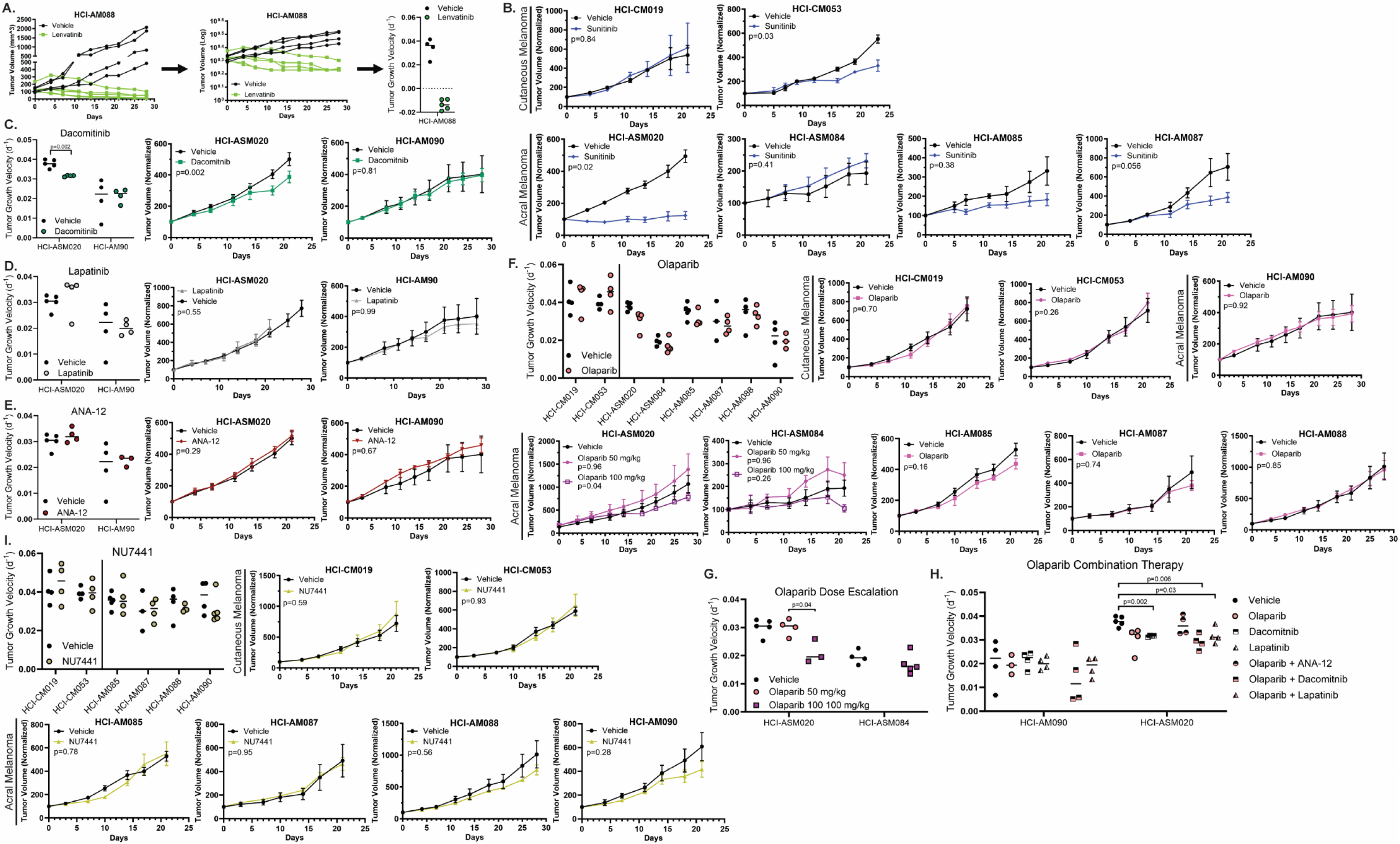
Drug response in PDX tumor models. **(A)** An example of the transformations for creating tumor growth velocity and HCI-AM088 is shown per Hather 2014^48^. In the left panel, individual tumor sizes are shown in the HCI-AM088 Lenvatinib and vehicle cohorts. These values undergo Log10 transformation using an absolute minimum tumor size of 50mm^3^ to avoid exponential data skewing from small tumor volumes (middle panel). In the right panel, the logarithmic slope of each tumor is plotted as a single dot in the tumor growth velocity plot. Negative average growth rates represent regression and positive growth rates indicate tumor growth. Near-zero average growth rates are oncostatic. **(B)** Average normalized tumor sizes are shown for each PDX model treated with Sunitinib. Tumor growth velocities and averaged normalized tumor sizes are shown for **(C)** Dacomitinib, **(D)** Lapatinib, **(E)** ANA-12, **(F)** Olaparib alone and with **(G)** dose escalation and **(I)** combination drug studies, and **(J)** NU7441. P-values are calculated with the student T-test per Hather 2014^48^.

**Supplemental Figure 4:**
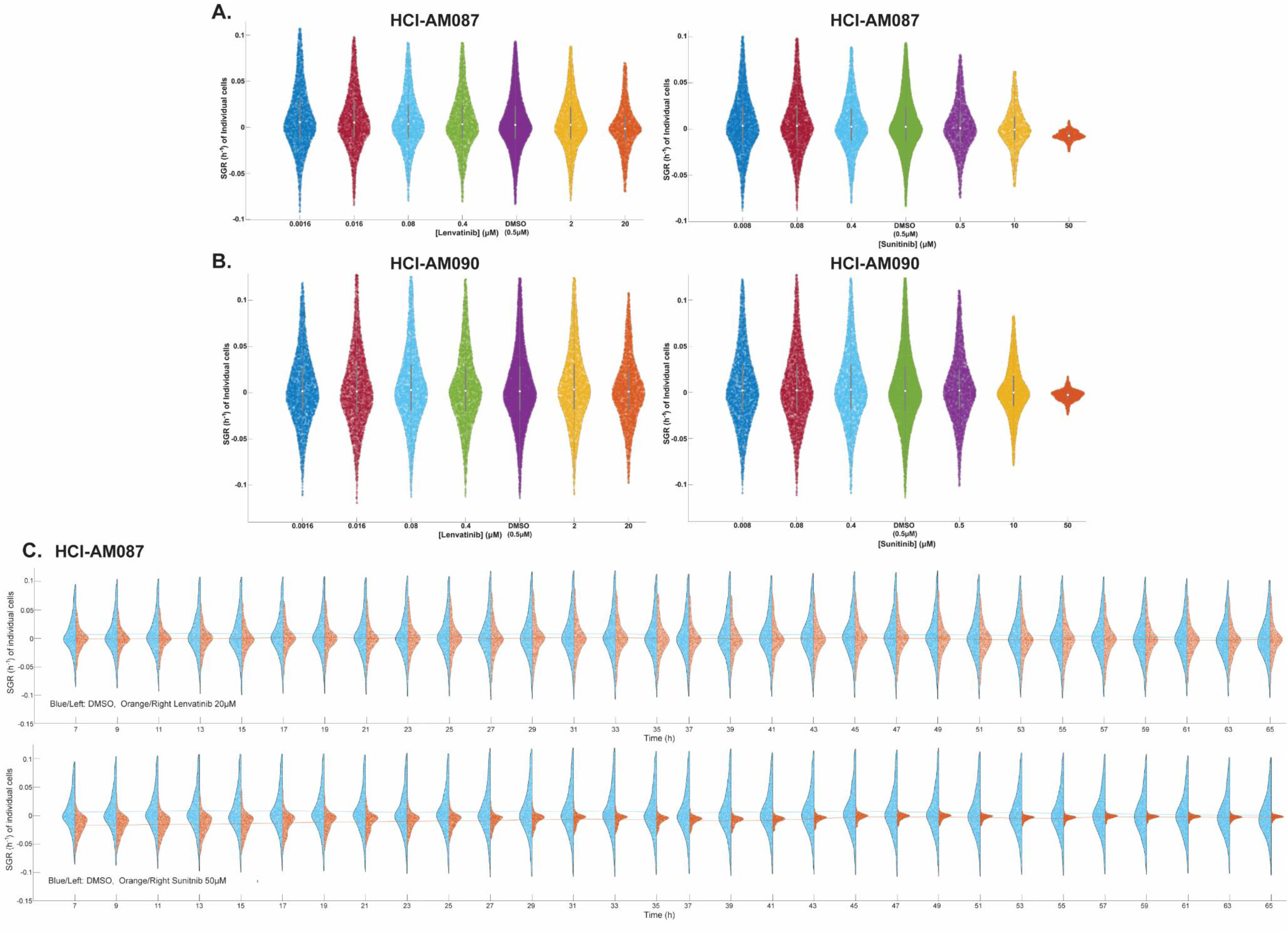
QPI representative SGR population dynamics. Representative SGR violin plots of all analyzed **(A)** HCI-AM087 and **(B)** HCI-AM090 cells are shown at tested concentrations of Lenvatinib and Sunitinib. The white central dot represents the average, grey bars represent standard deviation, and individual colored dots represents individual cells. Higher concentrations of drugs often have fewer cells, leading to smaller violins. **(C)** Representative two-sided violin plots show differences in SGR for DMSO (blue, left) and either Lenvatinib or Sunitinib (orange, right). Colored lines represent mean of each population, x-axis represents time (h), and y-axis represent SGR (h^-1^)

**Supplemental Figure 5:**
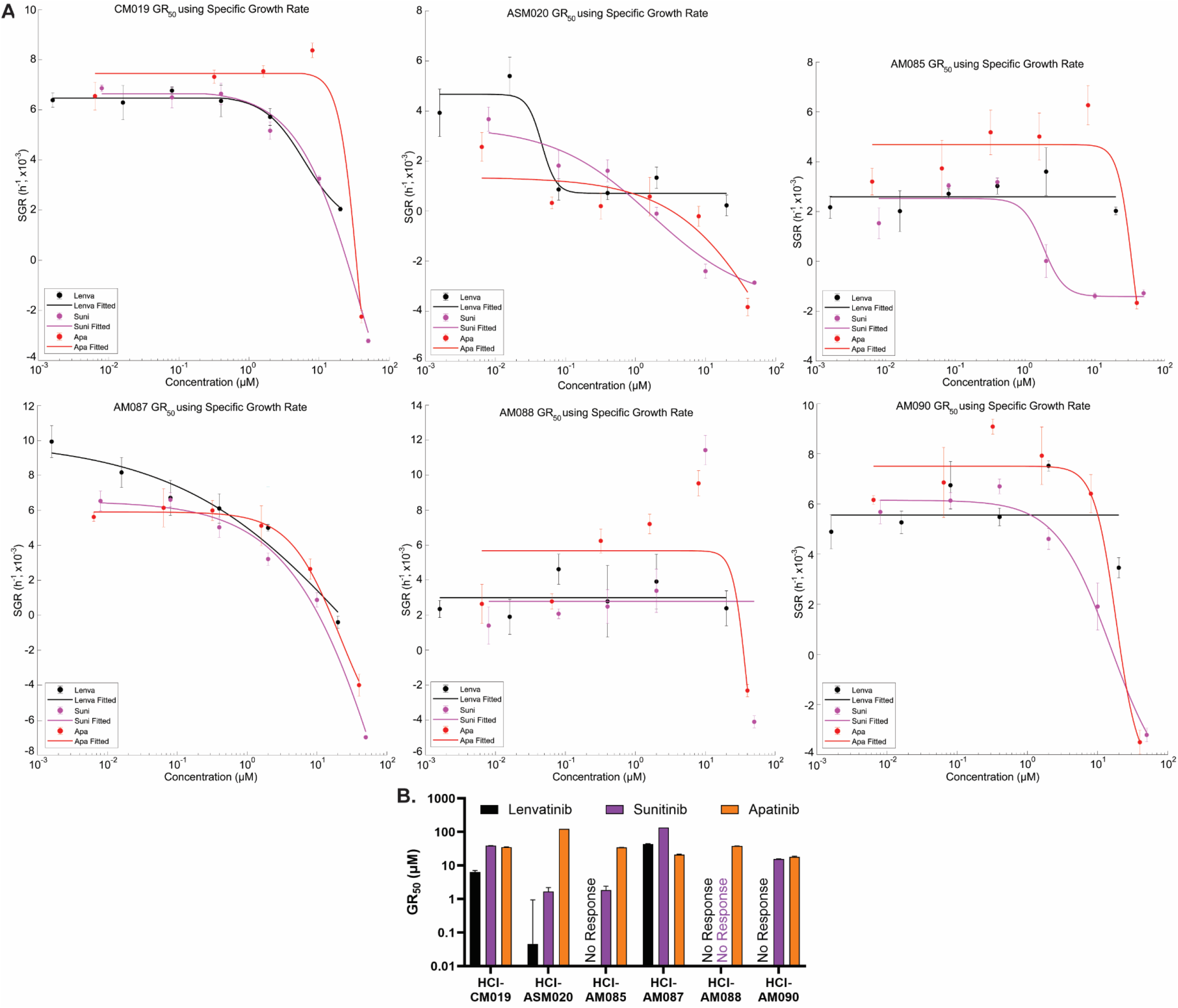
QPI GR_50_ best-fit curves. **(A)** Best fit curves used to identify GR_50_ in the different PDX cell cultures. Horizontal lines indicate failure to calculate or extrapolate a GR_50_ value. **(B)** Comparison of GR_50_ in each cell line for Sunitinib and the dual FGFR/VEGFR inhibitors Lenvatinib and Apatanib.

**Supplemental Figure 6:**
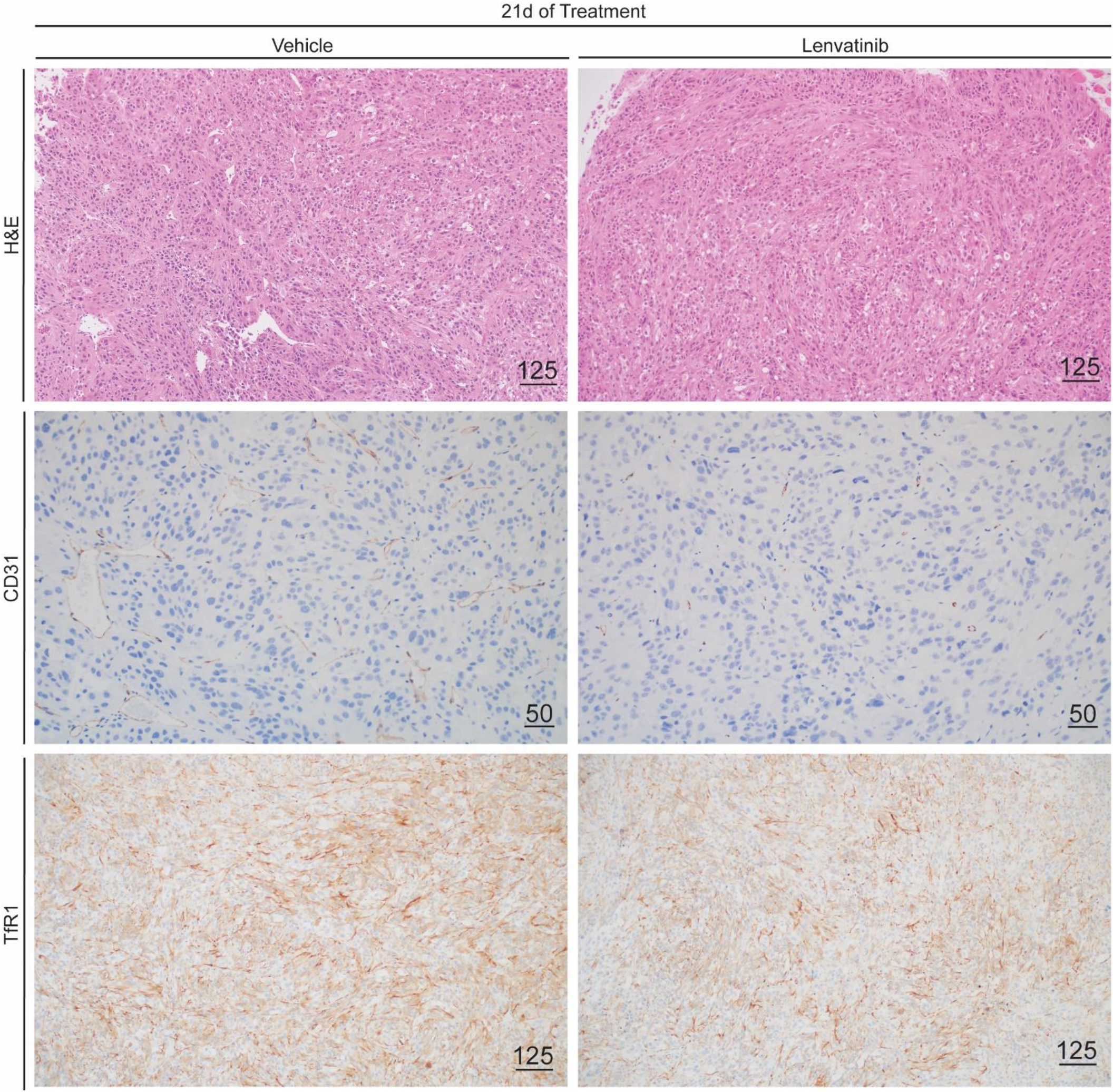
Larger images of TfR1 and CD31 immunohistochemistry stains. H&E and Tfr1 stains were imaged with a 10x objective, and CD31 was imaged with a 20x objective. Scale bars indicate length in microns.

