## Supplemental Data 6 for "Receptor tyrosine kinase inhibition leads to regression of acral melanoma by targeting the tumor microenvironment"

### HCI-CM002, Cutaneous Melanoma

| Primary Tumor Location | PDX Tumor Location | Other Known Metastatic Locations |
| --- | --- | --- |
| Right Cheek Skin | Right Superficial Parotid Gland | Lung |

Clinical Tumor Histology

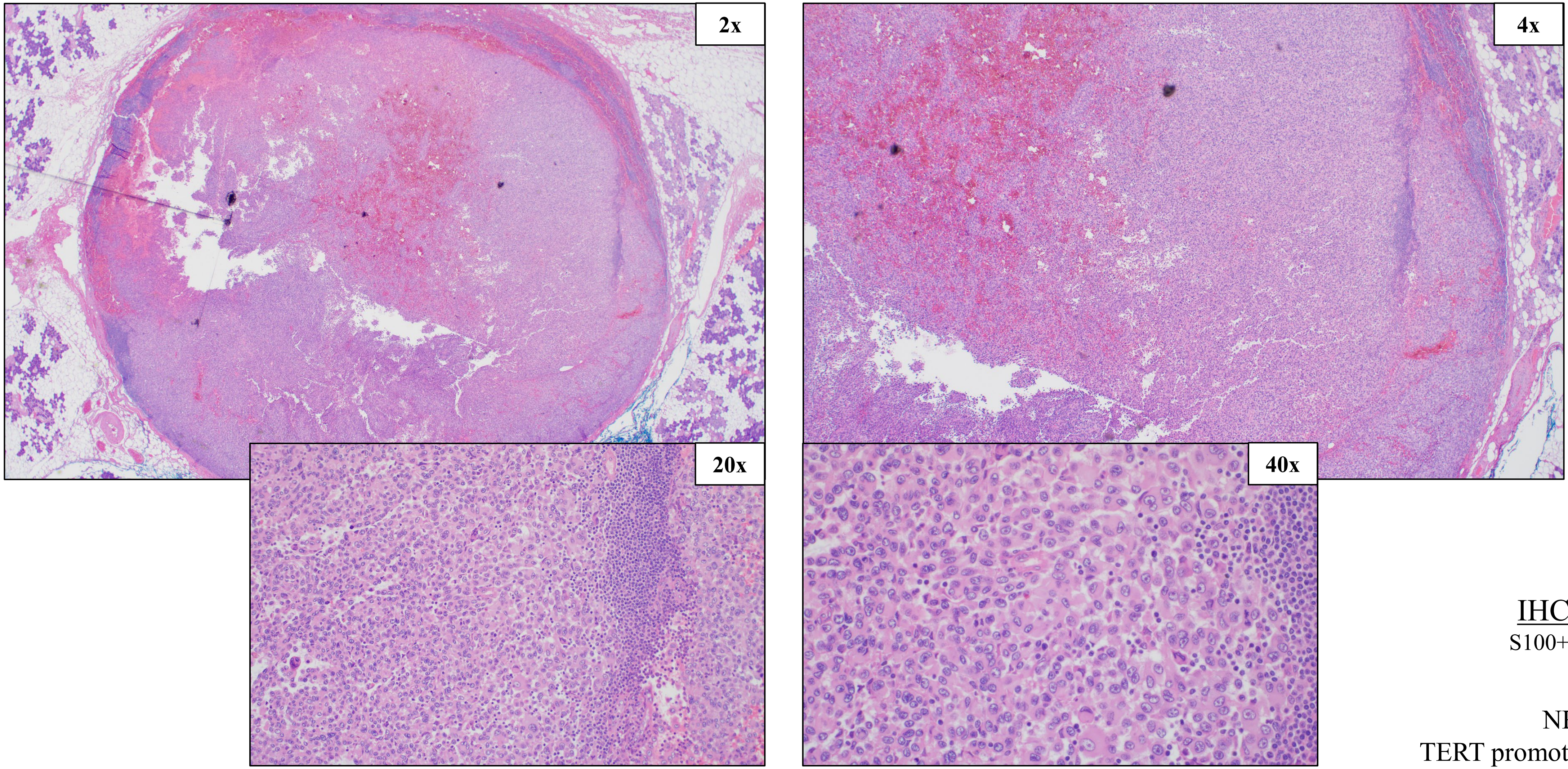


IHC Stains:
S100+, MelanA+

NRAS Q61R,
TERT promoter mutation

### HCI-CM002, Cutaneous Melanoma

#### SNV (%VAF)

NRAS c.182A>G p.Q61R (87%)
TERT promoter, chr5:1295242GG>AA (94%)

#### CNV

Amplification: NRAS, BRCA1
Loss: PTEN, TERT, MC1R
Deep deletion: CDKN2A

#### PDX Tumor Histology, Passage 2

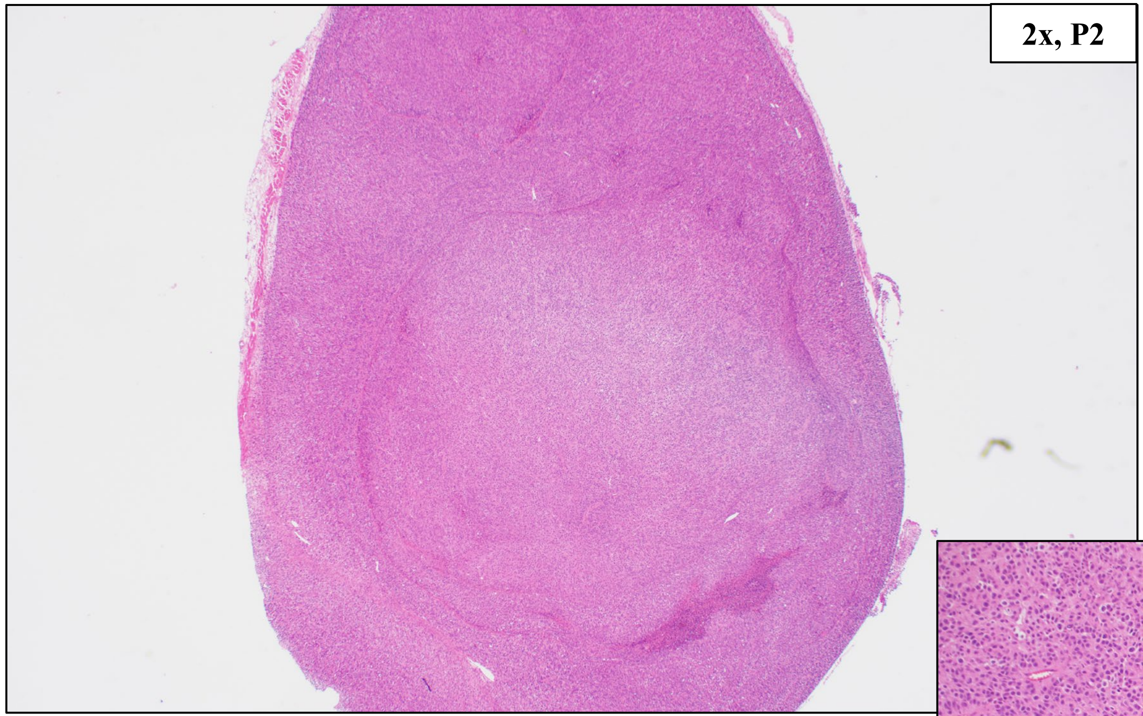


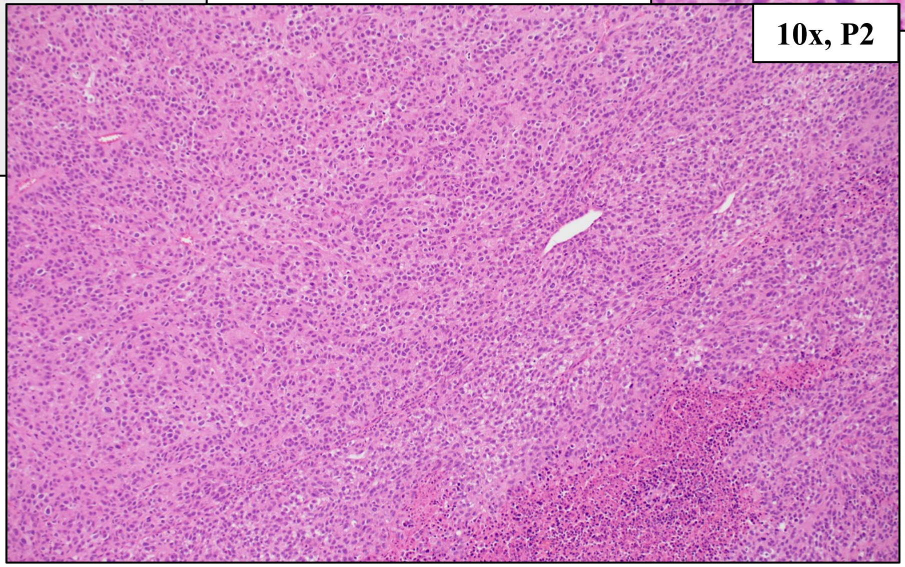


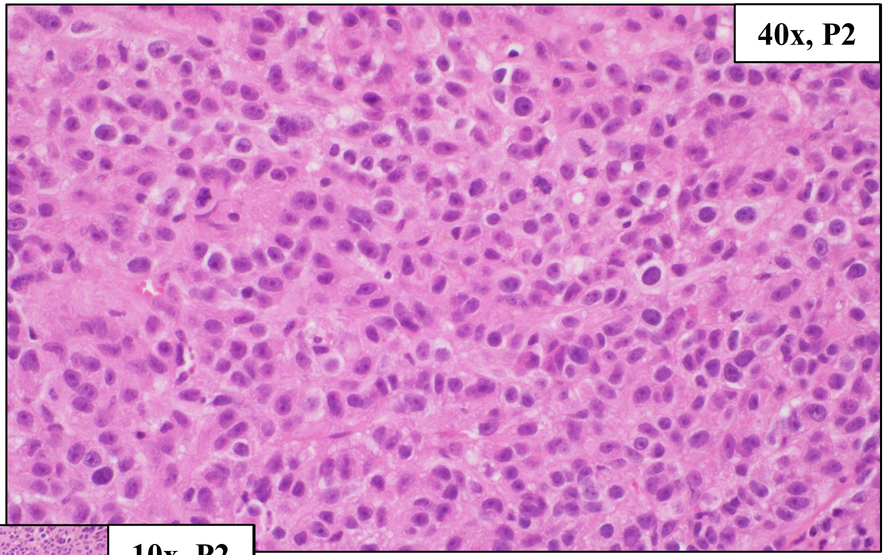


NRAS Q61R, TERT promoter mutation

### HCI-CM004, Cutaneous Melanoma (from same patient as HCI-CM019)

| Primary Tumor Location | PDX Tumor Location | Other Known Metastatic Locations |
|---|---|---|
| Scalp | Omentum | Brain, Bilateral Lungs, Back Skin, Supraclavicular Lymph Nodes, Brain |

Clinical Tumor Histology

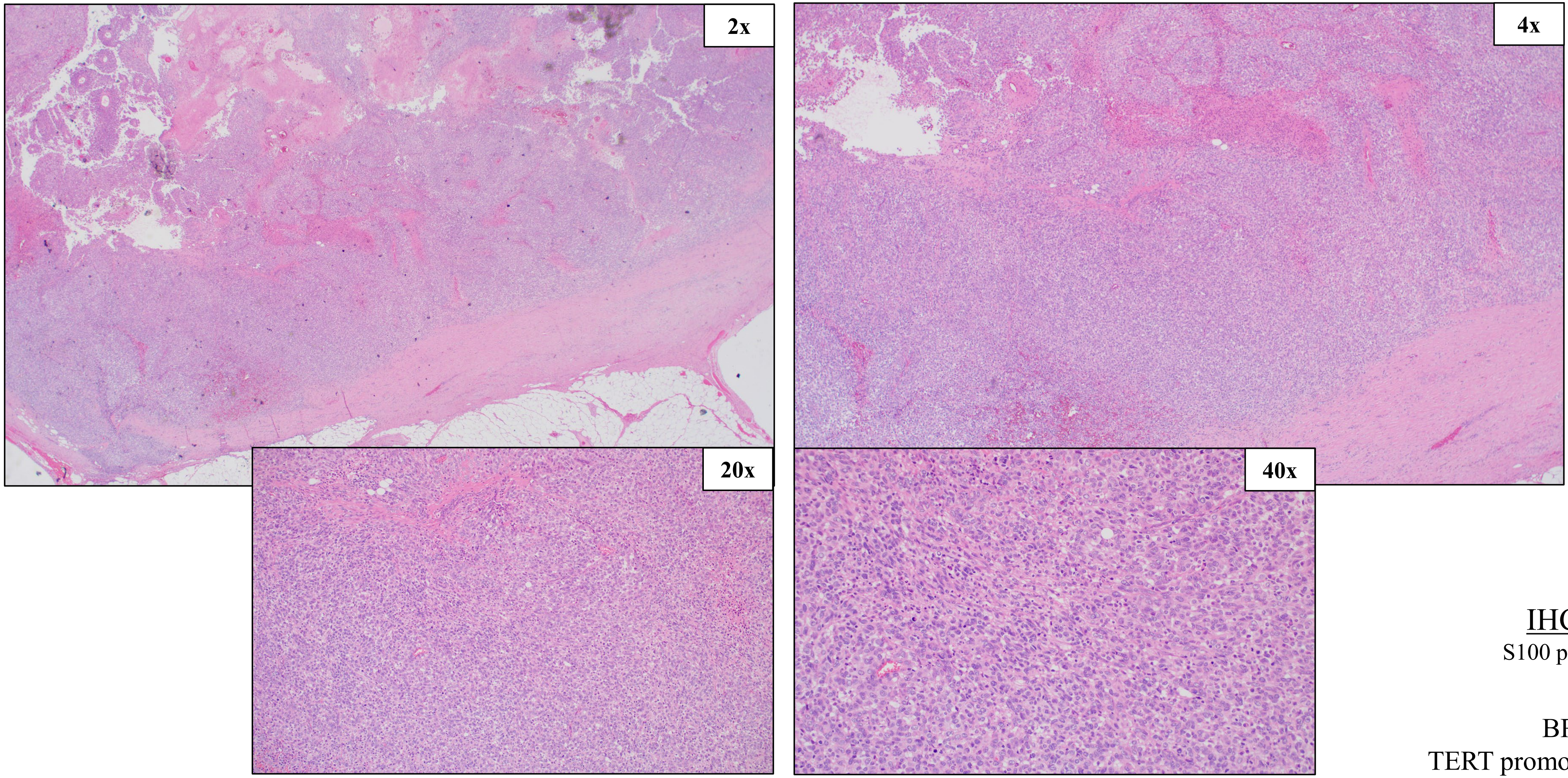


IHC Stains:
S100 partial positive

BRAF V600E,
TERT promoter mutation

### HCI-CM004, Cutaneous Melanoma (from same patient as HCI-CM004)

<u>SNV (%VAF)</u>

BRAF c.1799T>A p.V600E (83%),
TERT chr5:1295250G>A (66%)

<u>CNV</u>

Amplification: MET, BRAF, BRCA2
Loss: PTEN
Deep deletion: CDKN2A

<u>PDX Tumor Histology, Passages 2 and 4</u>

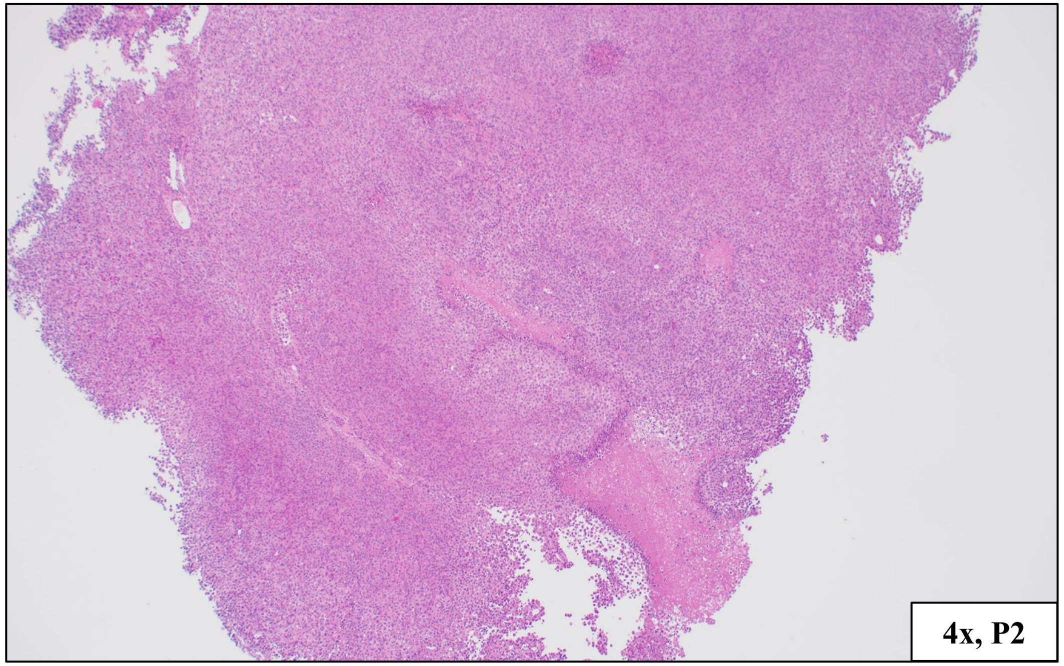


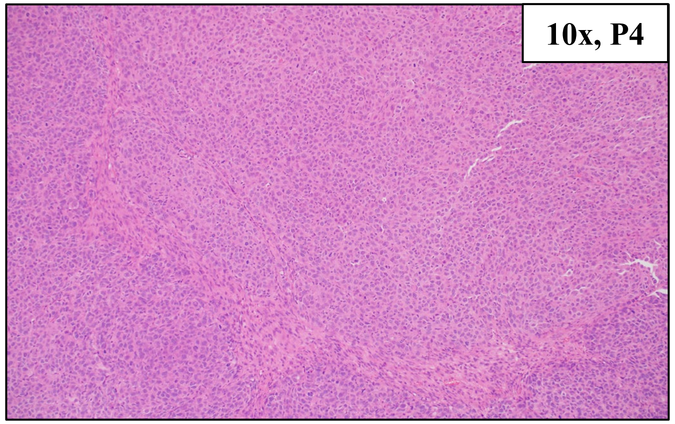


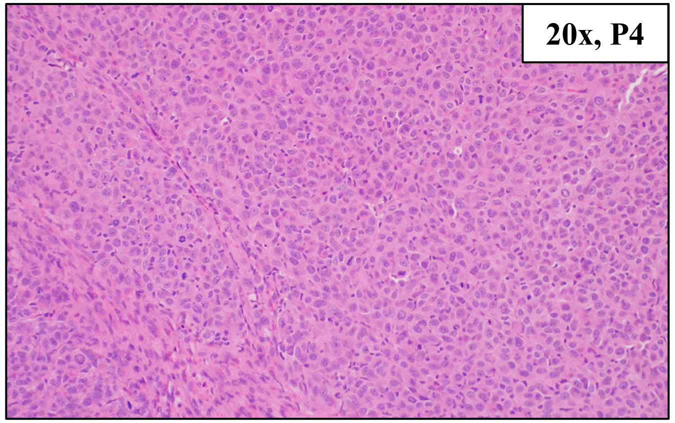


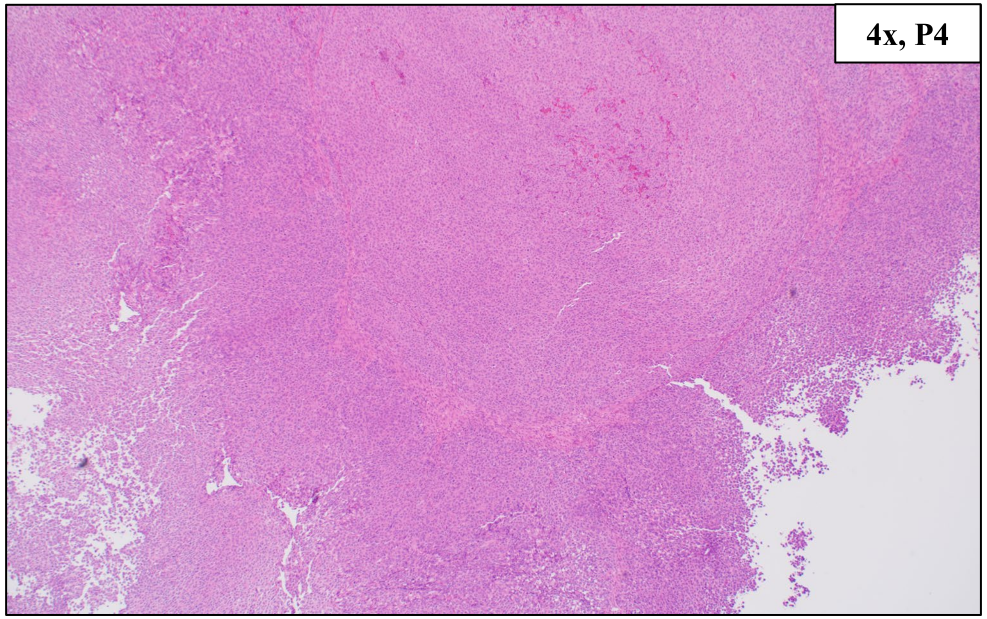


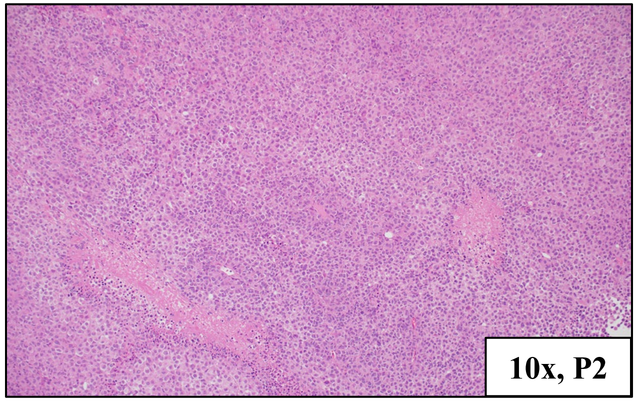


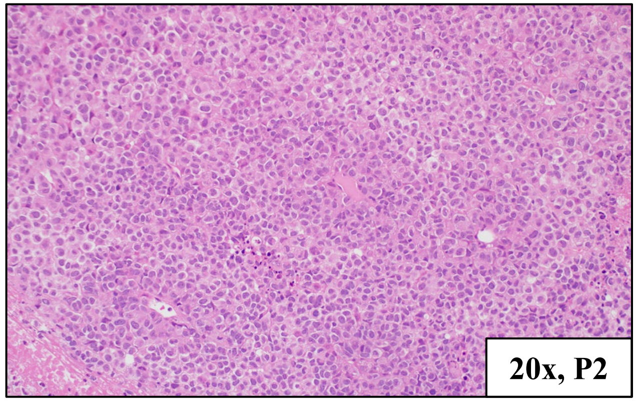


BRAF V600E, TERT promoter mutation

### HCI-CM005, Cutaneous Melanoma

| Primary Tumor Location | PDX Tumor Location | Other Known Metastatic Locations |
|---|---|---|
| Left Temporal Face Skin | Left Parotid | None |

Clinical Tumor Histology

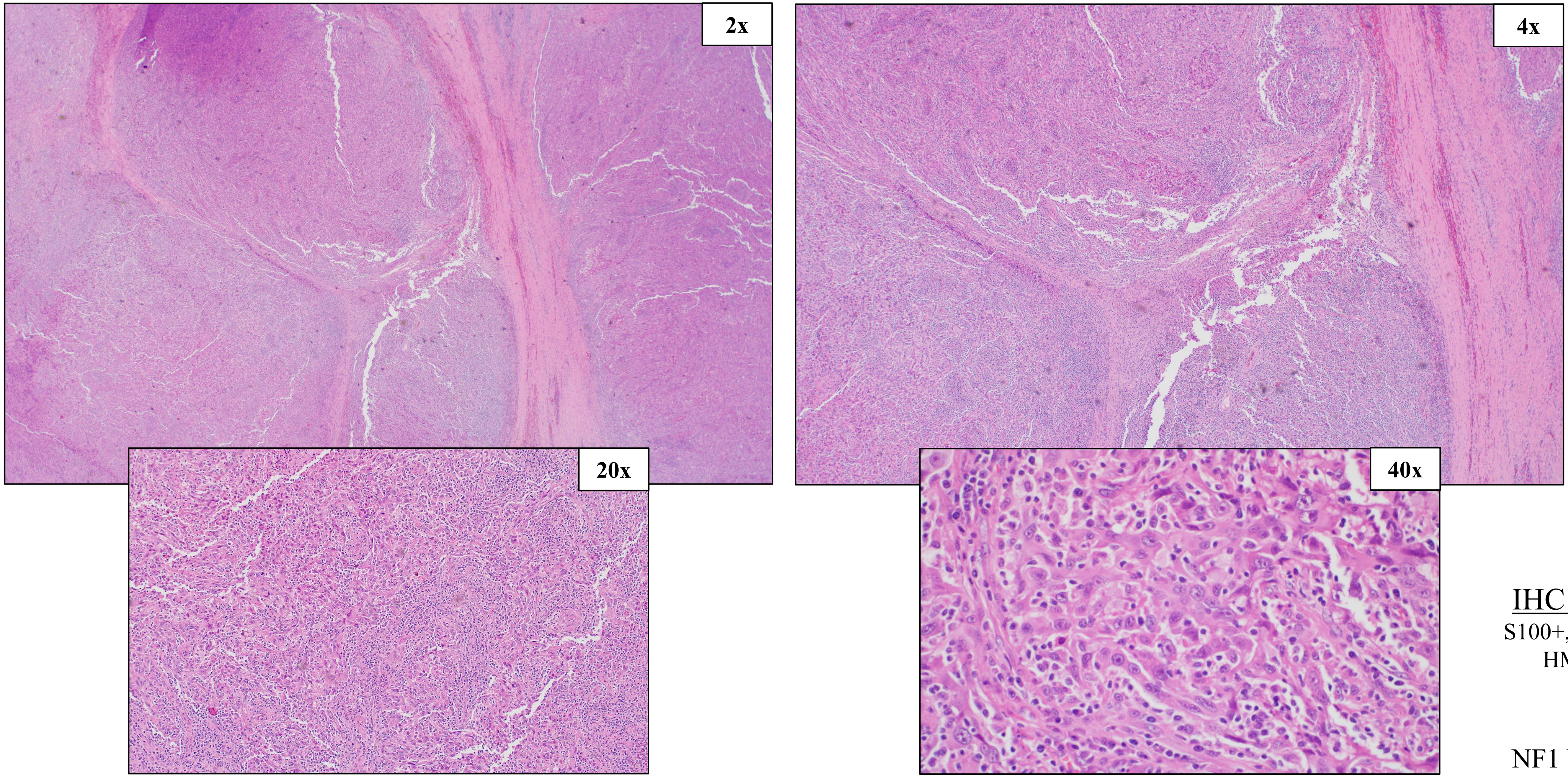


IHC Stains:
S100+, MelanA-, HMB45-

NF1 W221/Q282

### HCI-CM005, Cutaneous Melanoma

#### SNV (%VAF)

NF1 c.662G>A, p.W221* (47%)
NF1 c.844C>T, p. Q282* (49%)
TP53 c.637C>T, p.R213* (99%)

#### CNV

Amplification: BRCA1
Loss: ARID1B, MET, PTEN, BRCA2, RB1, TP53

#### PDX Tumor Histology

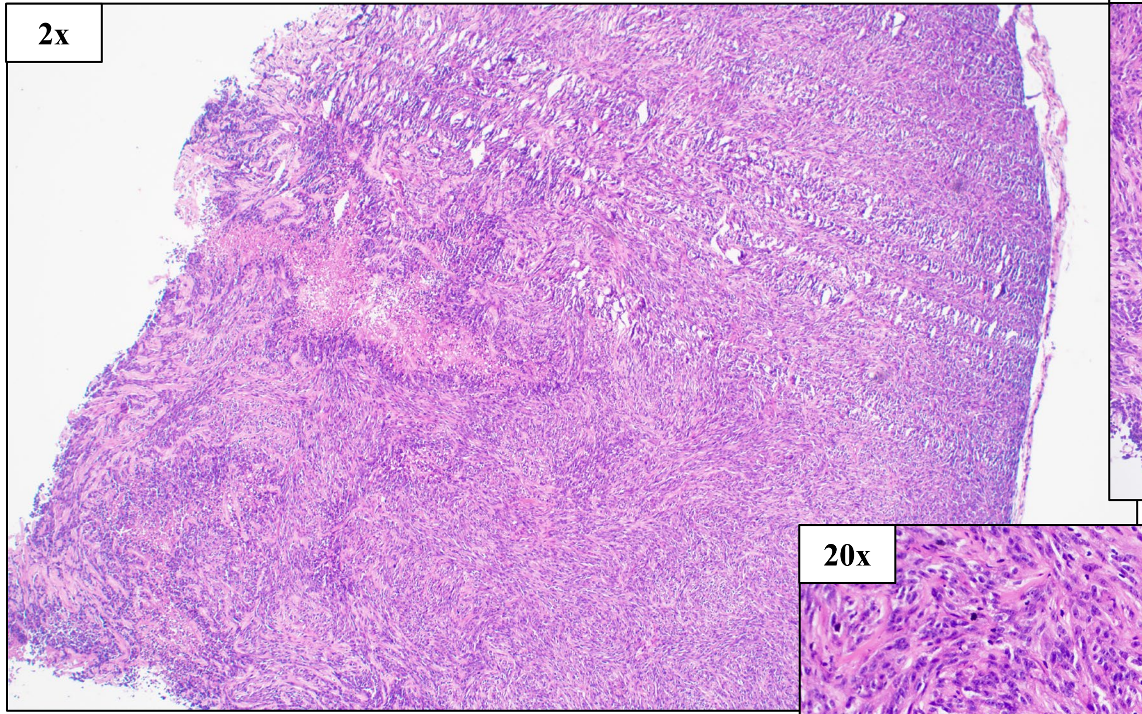


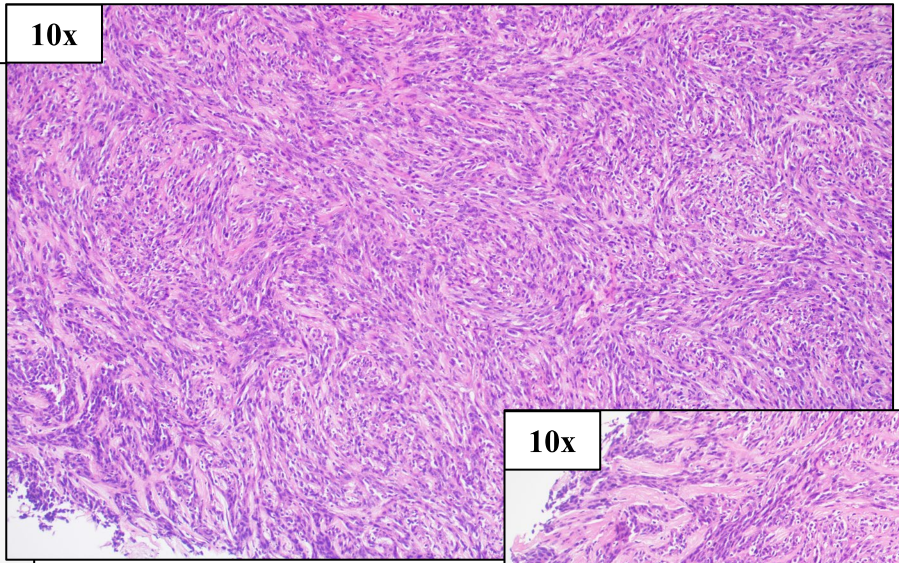


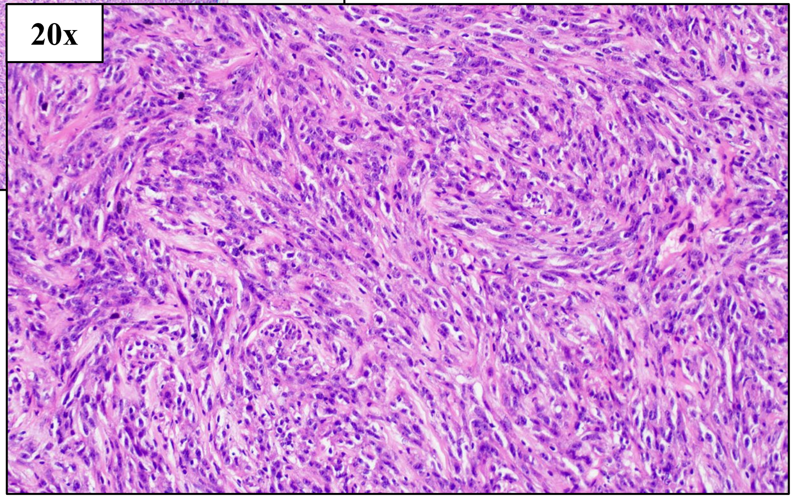


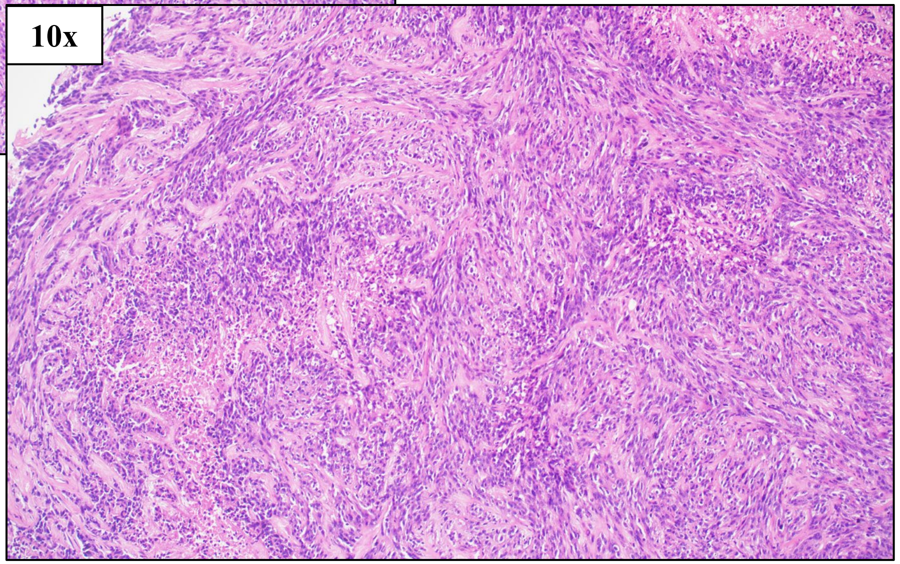


NF1 W221/Q282

### HCI-CM019, Cutaneous Melanoma (from same patient as HCI-CM004)

| Primary Tumor Location | PDX Tumor Location | Other Known Metastatic Locations |
|---|---|---|
| Scalp | Right Posterior Back Lymph Node | Omentum, Right and Left Lower Lung Lobes, Supraclavicular and Posterior Neck Lymph Nodes |

Clinical Tumor Histology

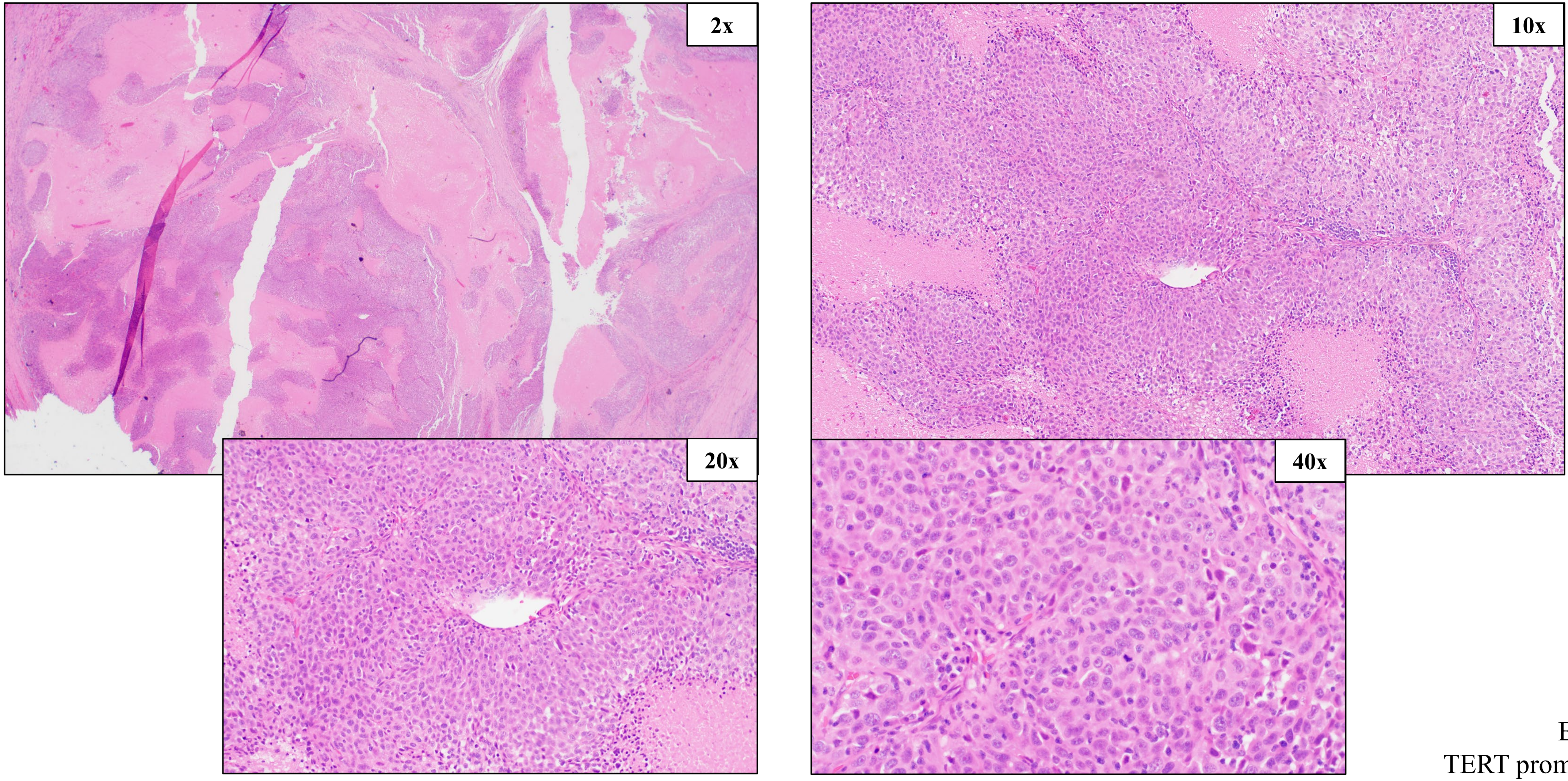


BRAF V600E,
TERT promoter mutation

### HCI-CM019, Cutaneous Melanoma (from same patient as HCI-CM004)

#### SNV (%VAF)

BRAF c.1799T>A p.V600E (83%),
TERT chr5:1295250G>A (52%)

#### CNV

Amplification: MET
Loss: ARID1B, CDKN2A, PTEN, KRAS, MC1R

#### PDX Tumor Histology, Passages 2 and 4

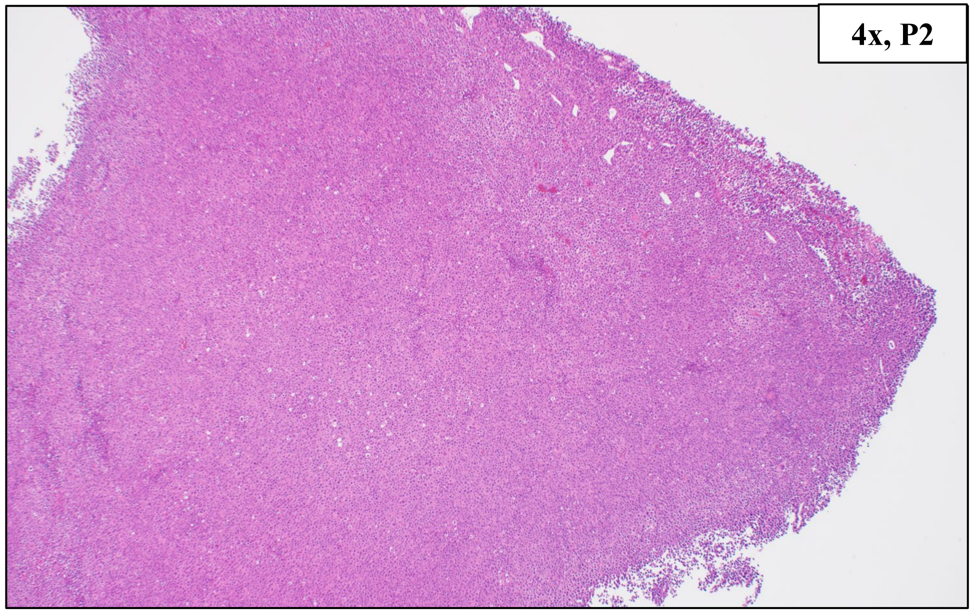


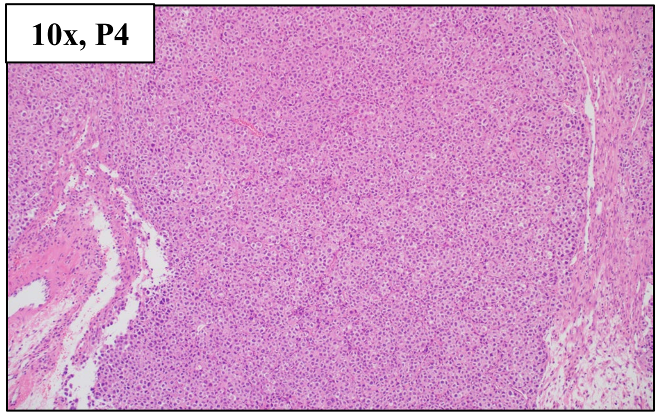


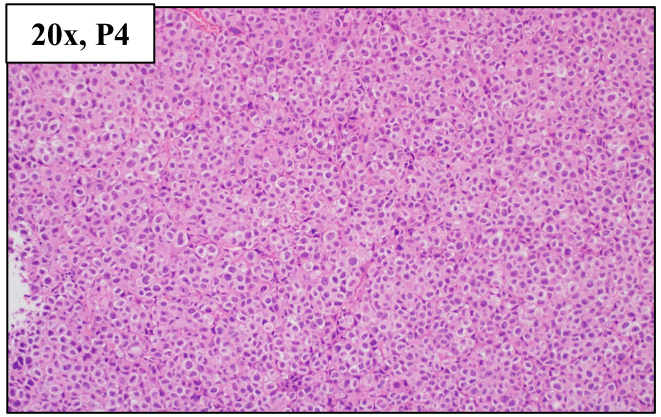


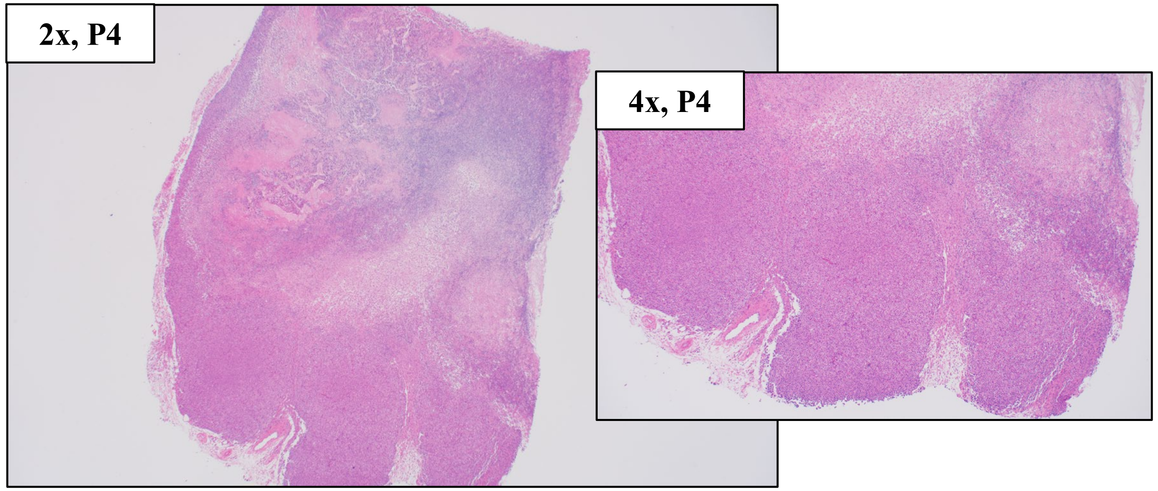


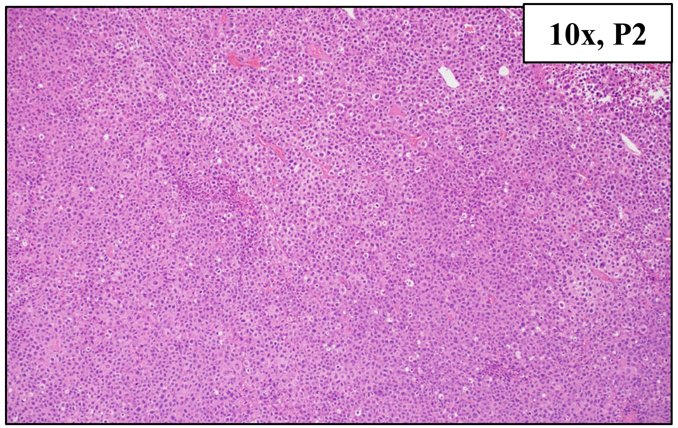


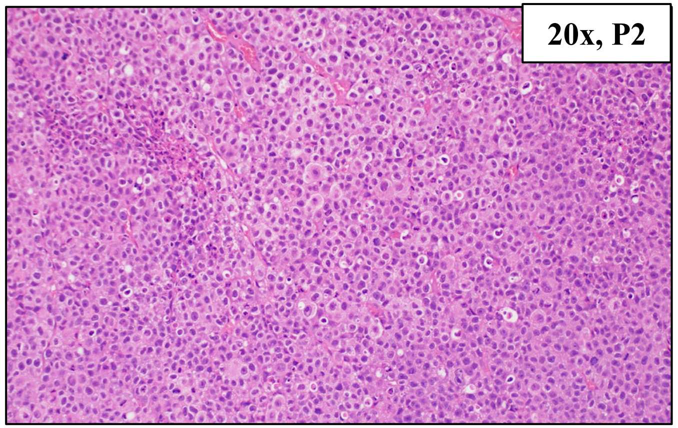


BRAF V600E, TERT promoter mutation

### HCI-ASM020, Acral Melanoma (from same patient as HCI-ASM021)

| Primary Tumor Location | PDX Tumor Location | Other Known Metastatic Locations |
|---|---|---|
| Left Hallux Toenail | Lower Abdominal Wall Lymph Node | Left Inguinal Lymph Nodes, Iliac Lymph nodes, Left Iliac Wing |

Clinical Tumor Histology

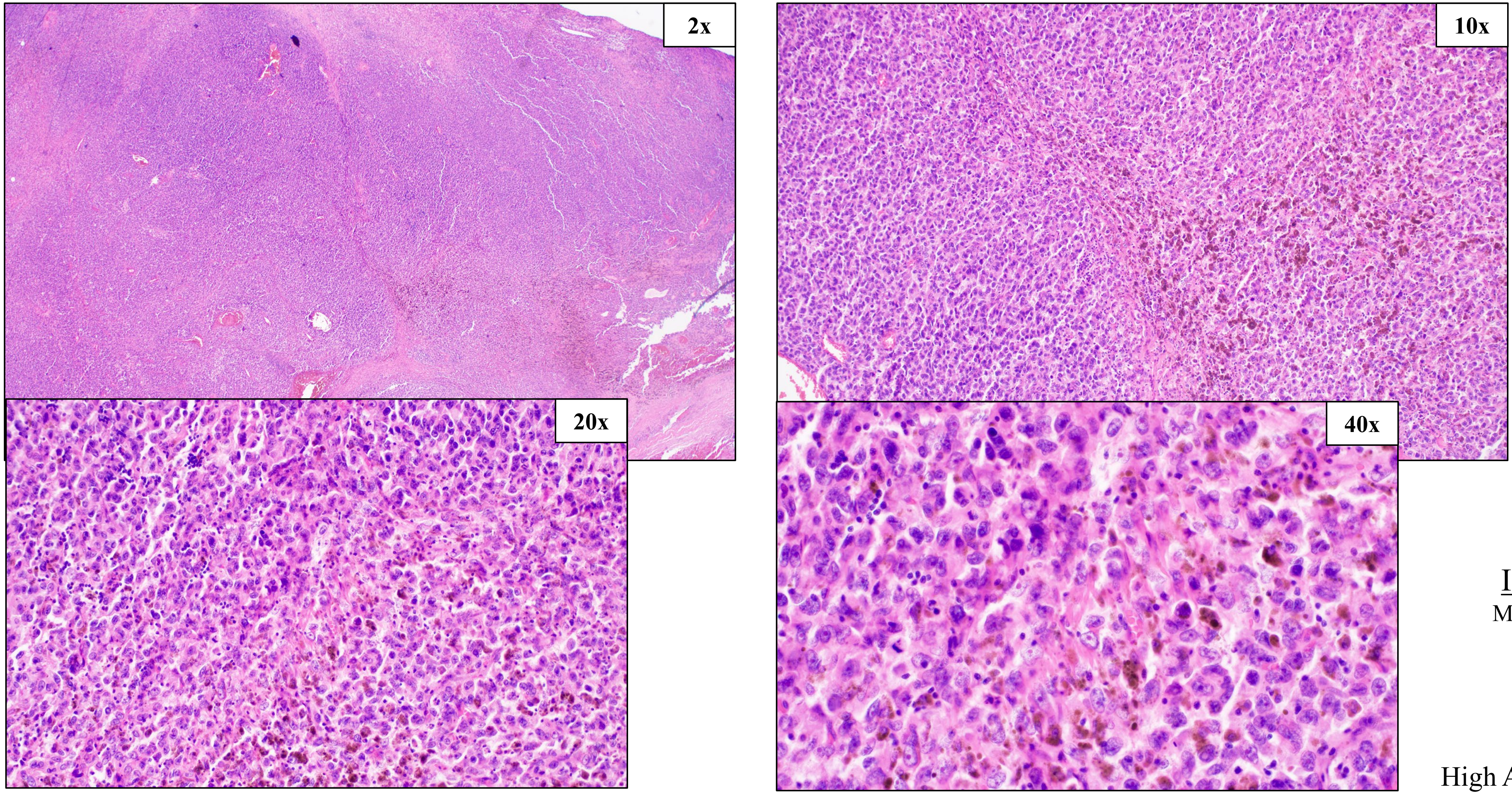


IHC Stains:

MelanA+, S100+

KIT N822K

High AMP: KIT, TERT

### HCI-ASM020, Acral Melanoma (from same patient as HCI-ASM021)

#### SNV (%VAF)

KIT c.2466T>G, p.N822K (89%)
SPRED1 c.867dup, p.S290IfsTer10 (95%)

#### CNV

Deep Deletion: CDKN2A
Loss: ARID1B, SPRED1
Amplification: EGFR, MET, HRAS, MC1R, MAP2K2, SMARCA4
Amplification ≥8 copies: KIT, TERT, MAP2K1

#### PDX Tumor Histology, Passages 2 and 5

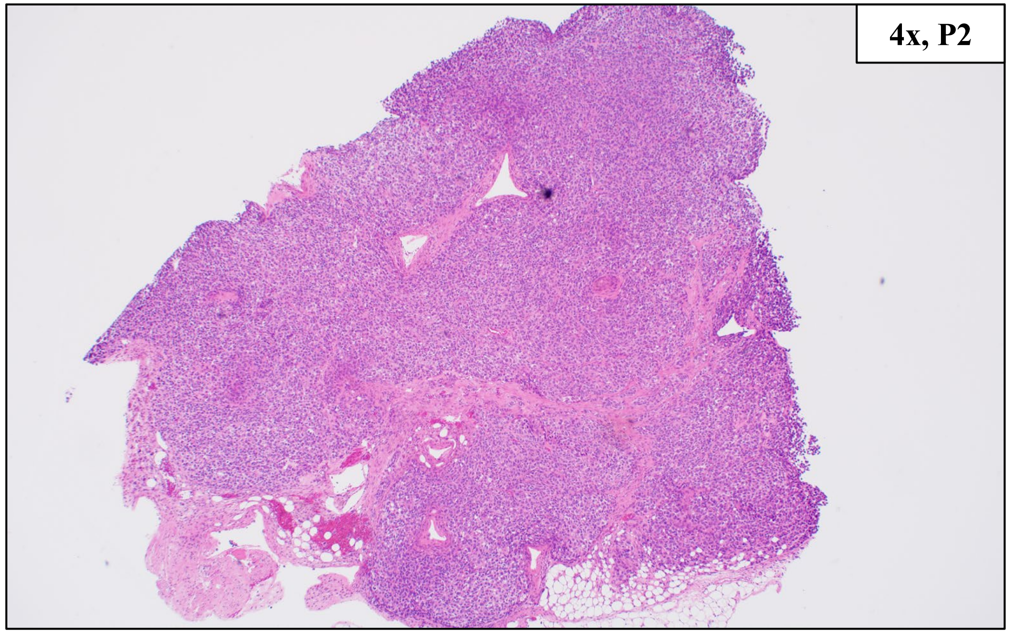
4x, P2

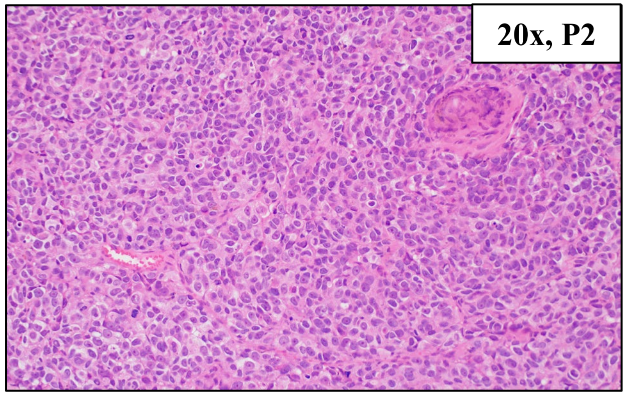
20x, P2

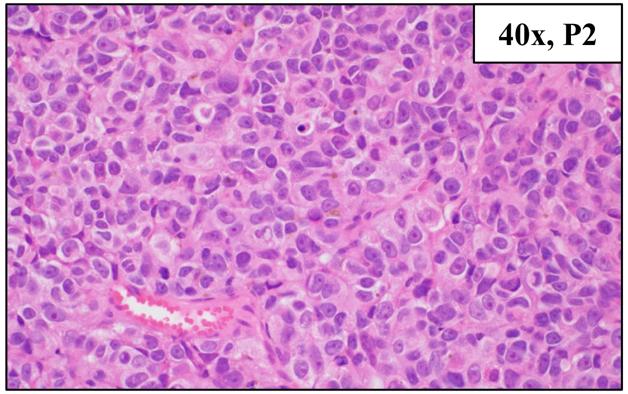
40x, P2

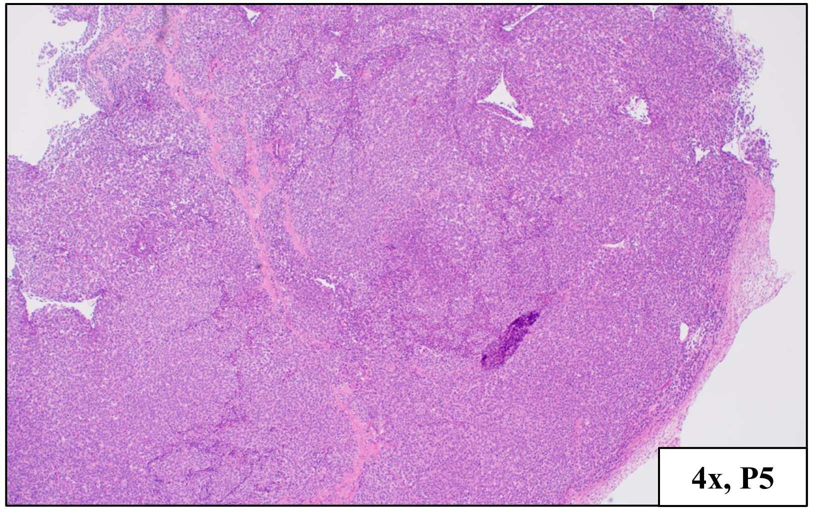
4x, P5

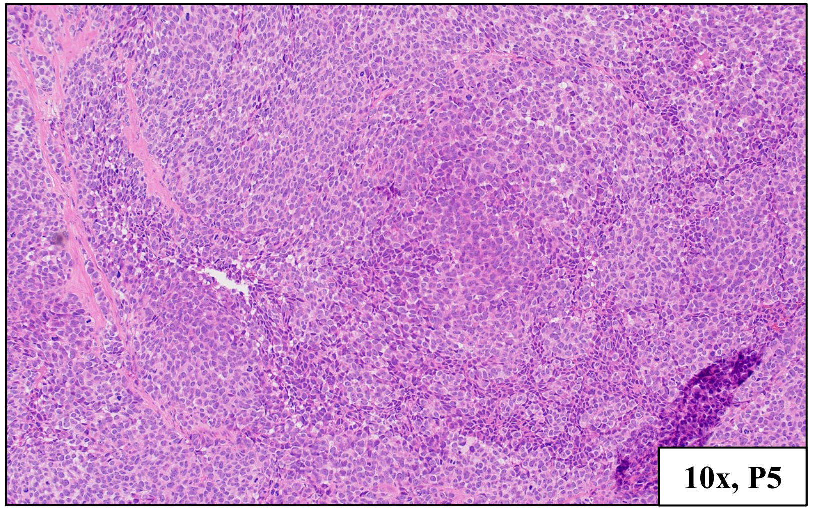
10x, P5

KIT N822K
High AMP: KIT, TERT

### HCI-ASM021, Acral Melanoma (from same patient as HCI-ASM020)

Primary Tumor Location

Left Hallux Toenail

PDX Tumor Location

Left Great Toe

Other Known Metastatic Locations

Left Lower Abdominal Wall and Inguinal Lymph Nodes, Iliac Lymph nodes, Left Iliac Wing

Clinical Tumor Histology

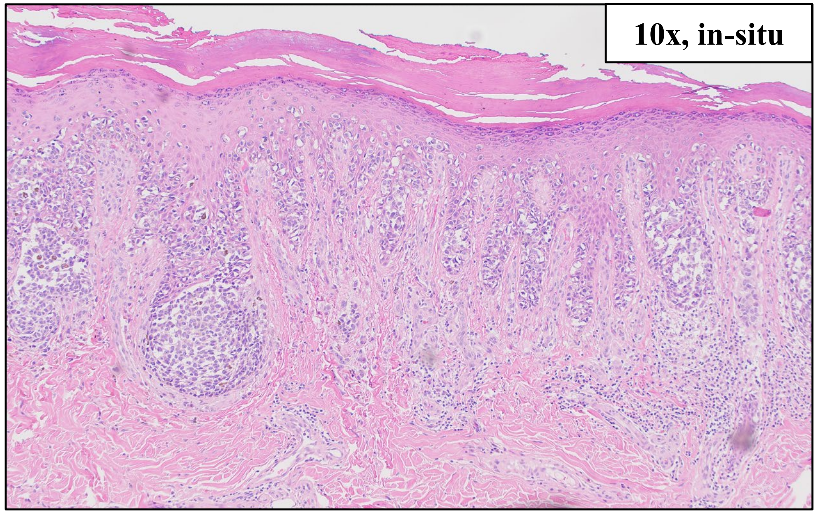


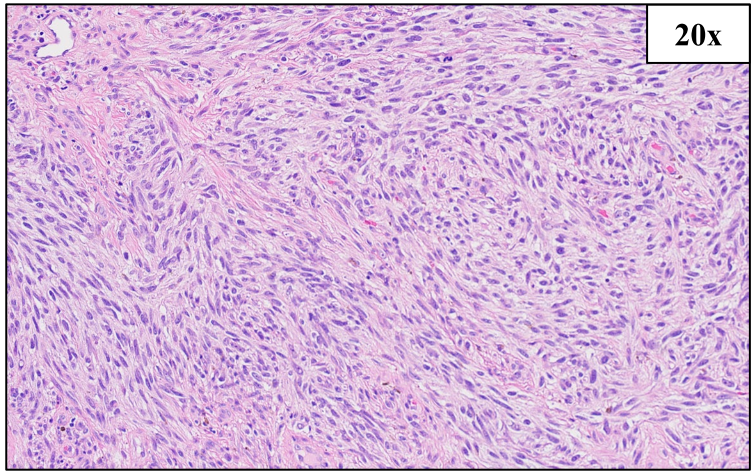


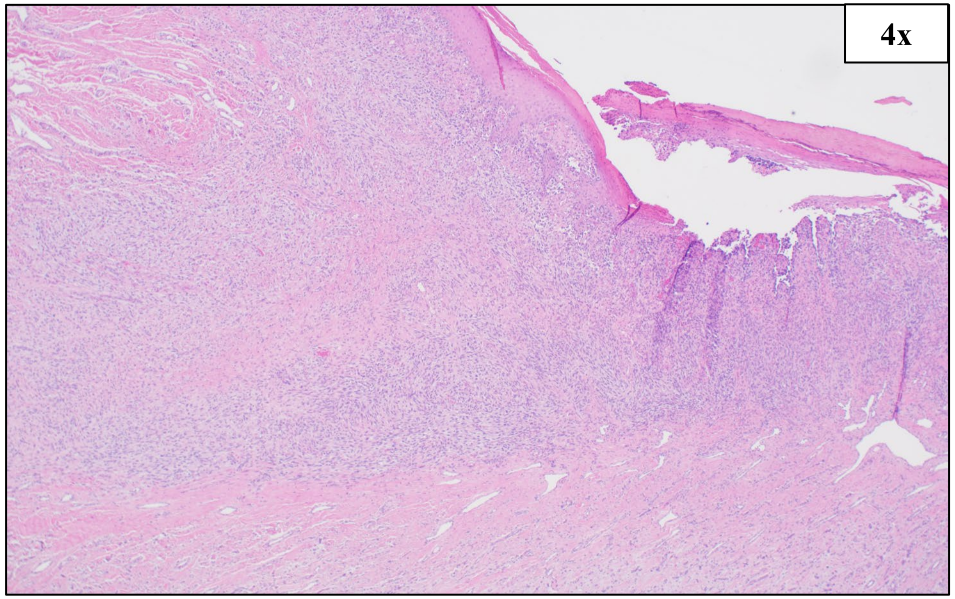


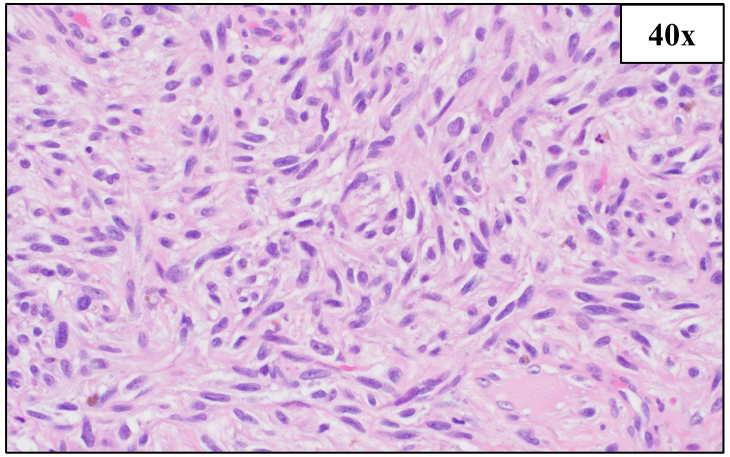


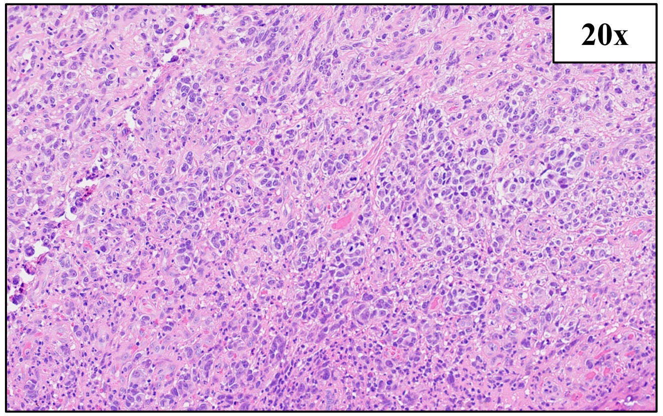


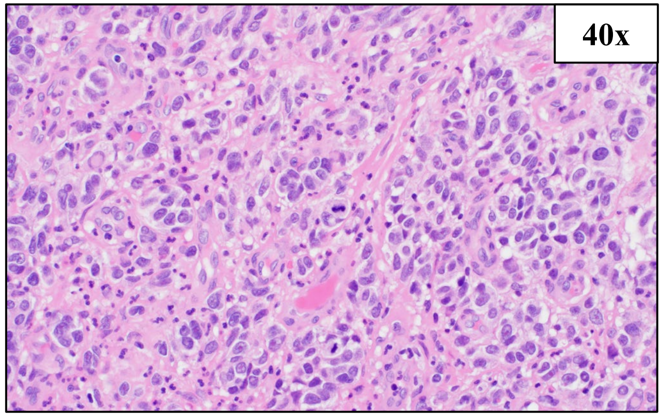


IHC Stains:

MelanA+, S100+

BRAF V600E, TERT promoter mutation,
High Amp: SMARCA4

### HCI-ASM021, Acral Melanoma (from same patient as HCI-ASM020)

#### SNV (%VAF)

BRAF c.1799T>A, p.V600E (63%)
TERT chr5:1295250G>A (99%)

#### CNV

Amplification ≥8 copies: SMARCA4
Amplification: TERT, MET, BRAF, KRAS, CDK4, MC1R
Deep Deletion: CDKN2A

PDX Tumor Histology, Passage 3

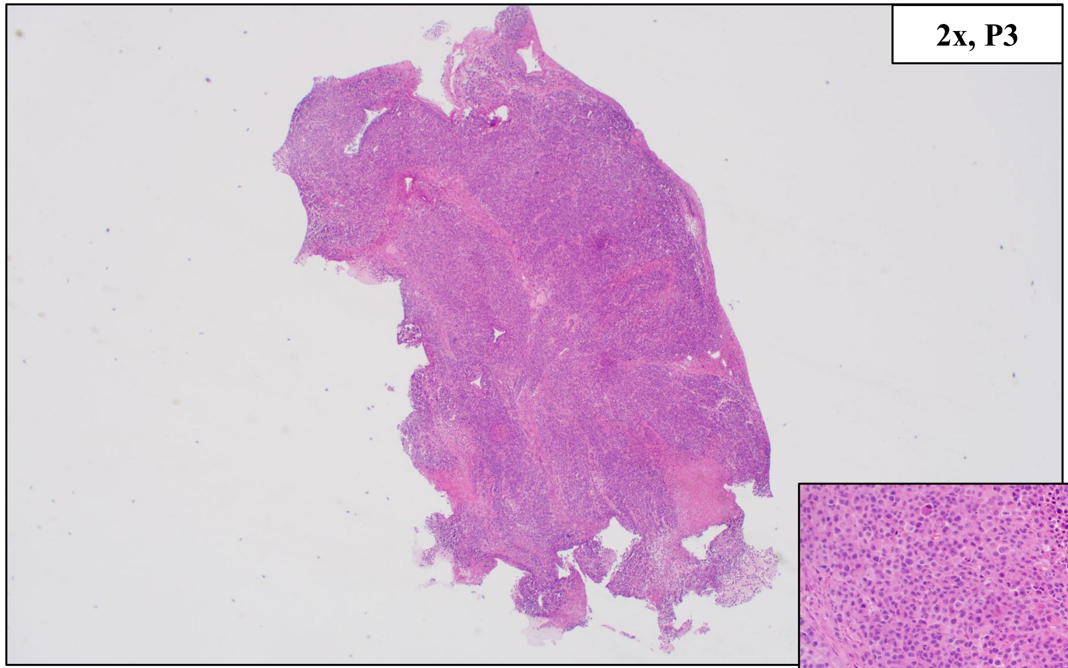


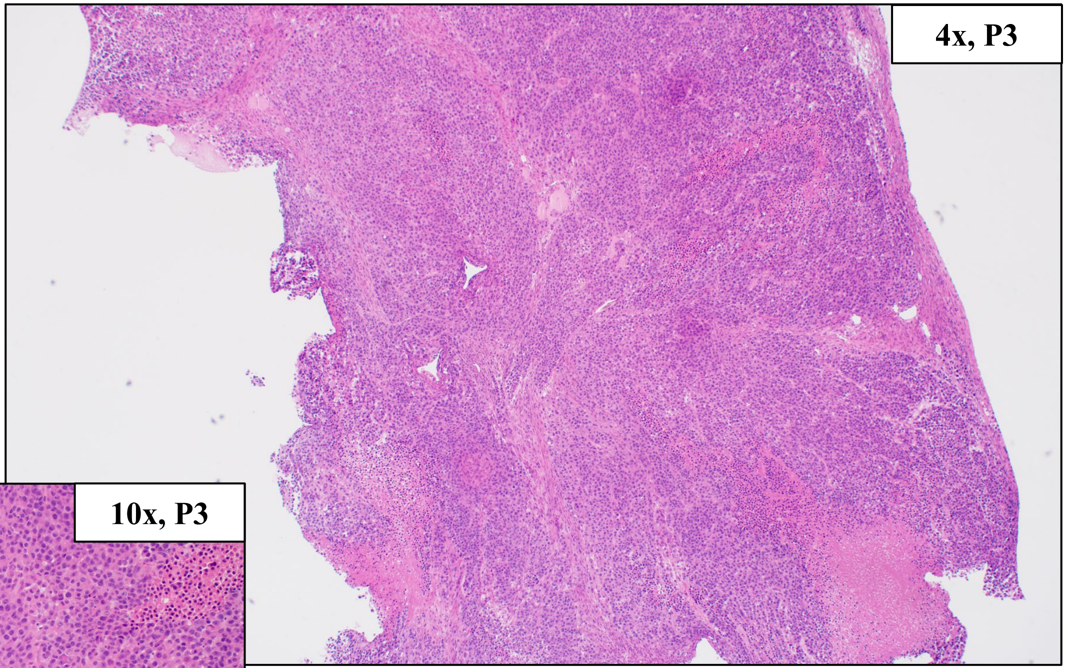


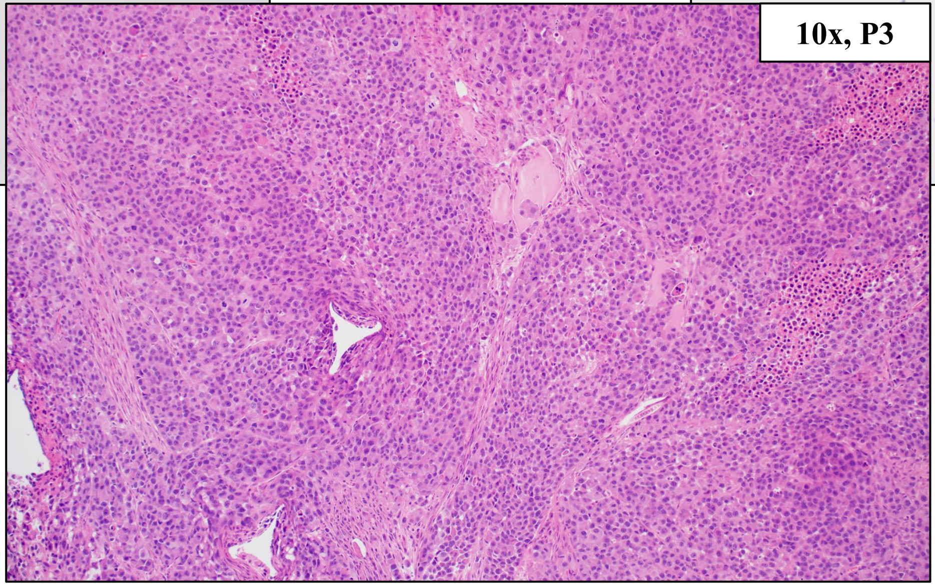


BRAF V600E, TERT promoter mutation,
High Amp: SMARCA4

### HCI-CM053, Cutaneous Melanoma

| Primary Tumor Location | PDX Tumor Location | Other Known Metastatic Locations |
| --- | --- | --- |
| L Shoulder | L Axillary Lymph Node | None |

Clinical Tumor Histology

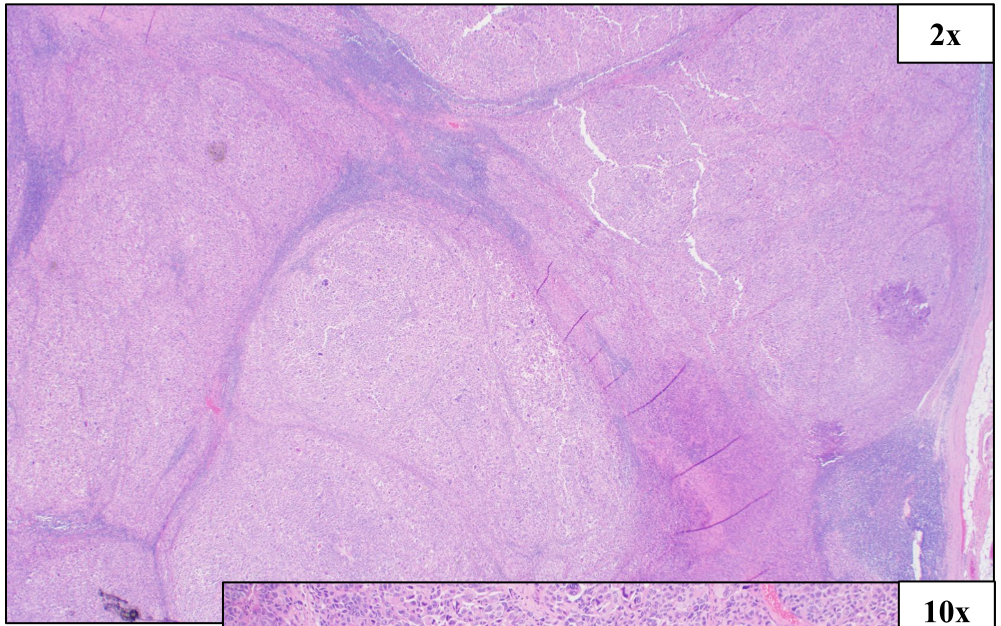


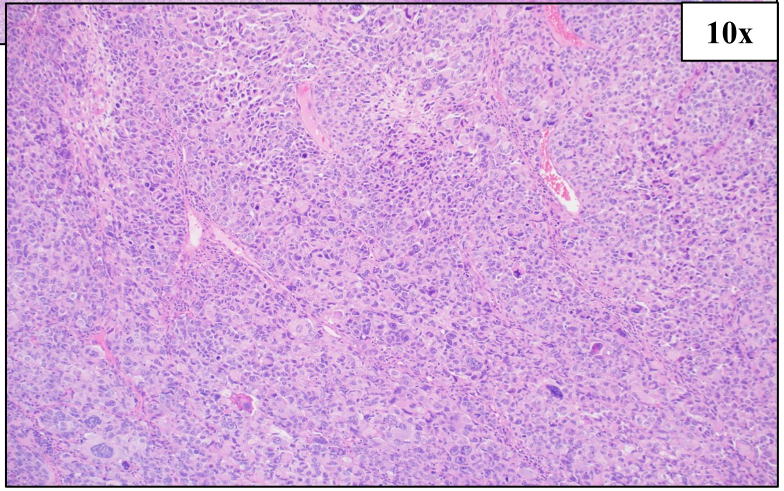


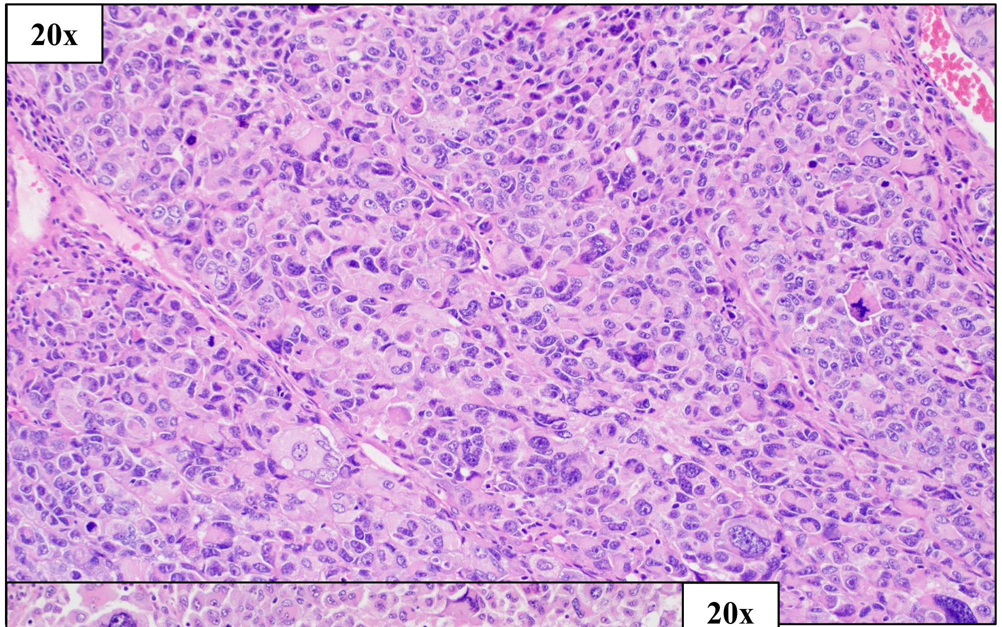


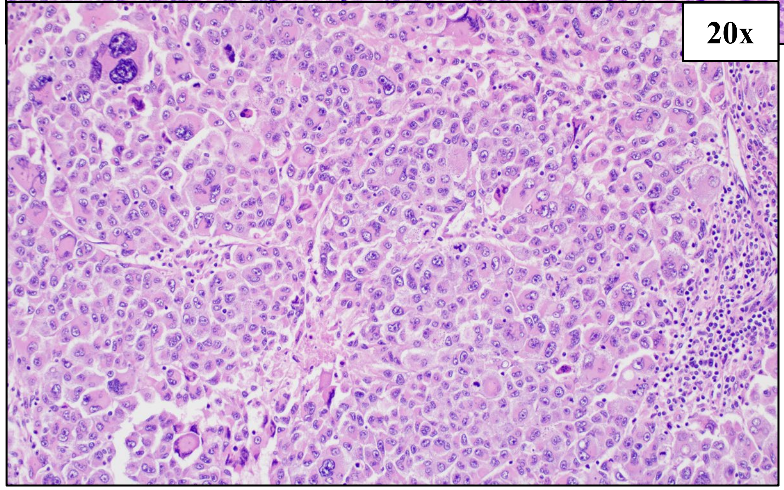


IHC Stains:
S100+

BRAF V600E, High Amp: TERT

### HCI-CM053, Cutaneous Melanoma

#### SNV (%VAF)

BRAF c.1799T>A, p.V600E (50%)
MC1R c.86dup, p.N29KfsTer14 (35%)
TP53 c.722C>T, p.S241F (99%)
RB1 c.802G>T, p.E268* (99%)

#### CNV

Amplification ≥8 copies: TERT
Loss: PDGFRA, KIT, PTEN, CCND1, TP53

#### PDX Tumor Histology, Passages 2 and 4

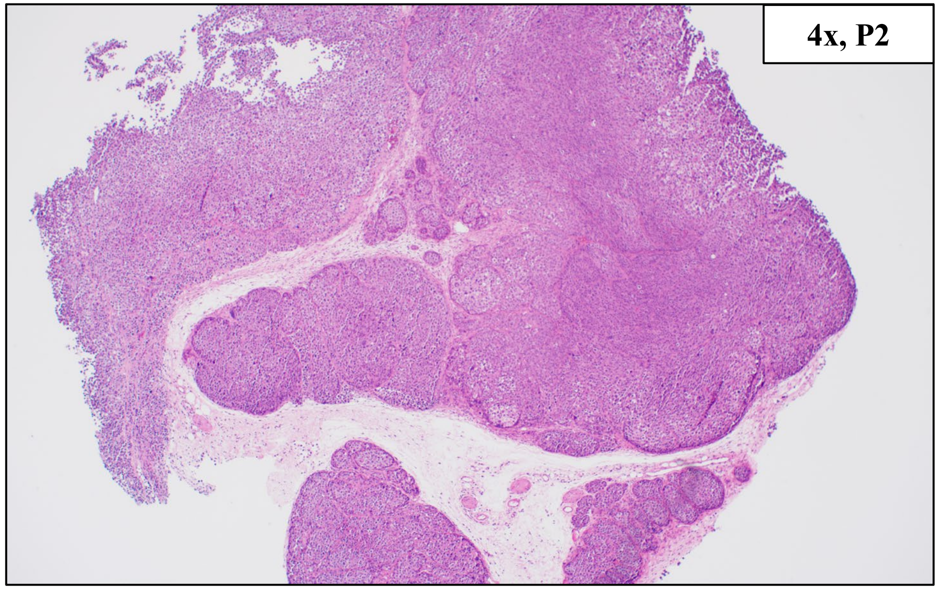


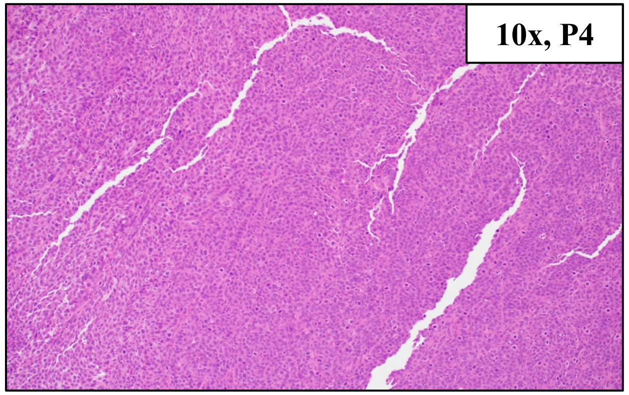


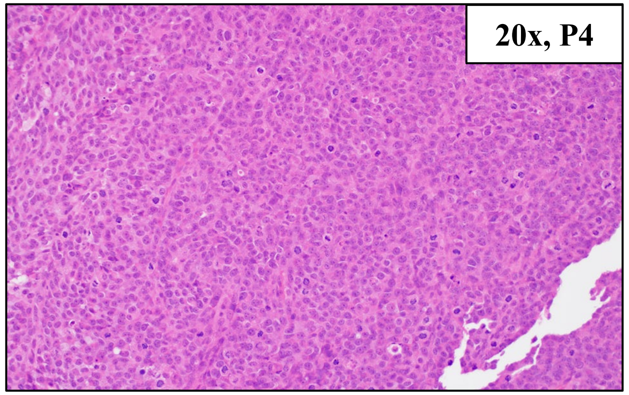


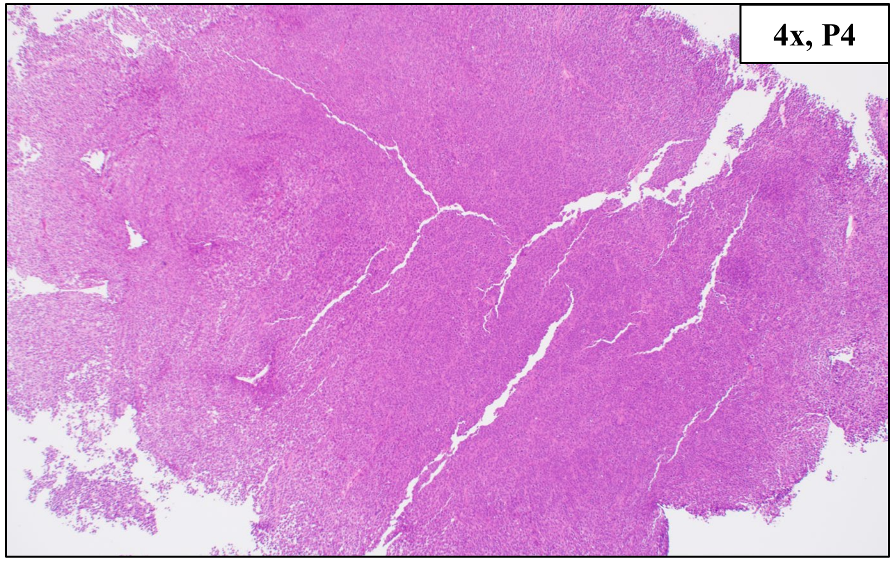


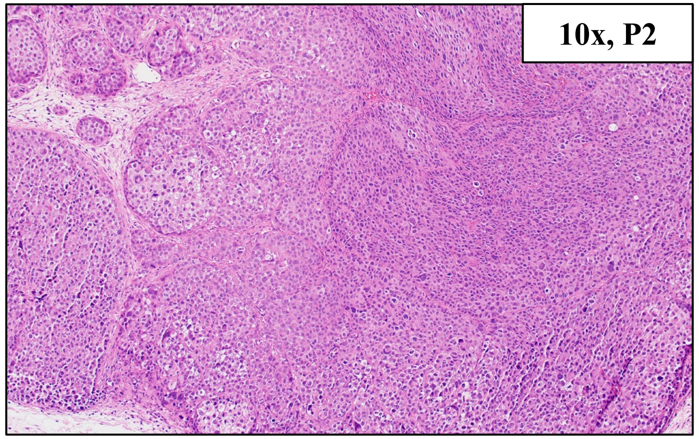


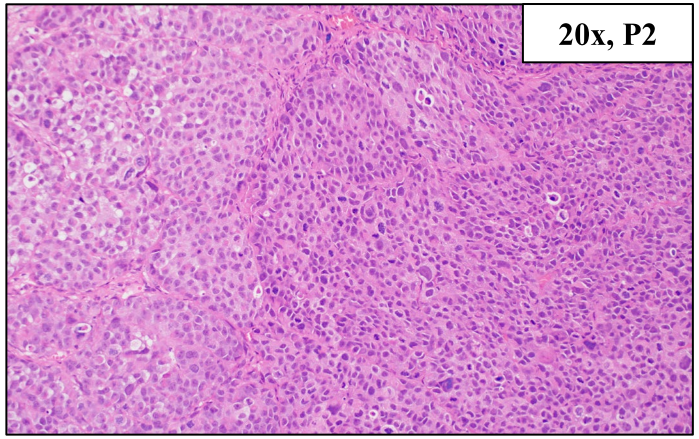


BRAF V600E, High Amp: TERT

### HCI-ASM084, Acral Melanoma

| Primary Tumor Location | PDX Tumor Location | Other Known Metastatic Locations |
|---|---|---|
| Left Thumb Skin | Left Lower Back or Right Anterior Thigh | Bilateral Lungs, Right Upper Arm, Left Lower Back, Right Anterior Thigh |

Clinical Tumor Histology

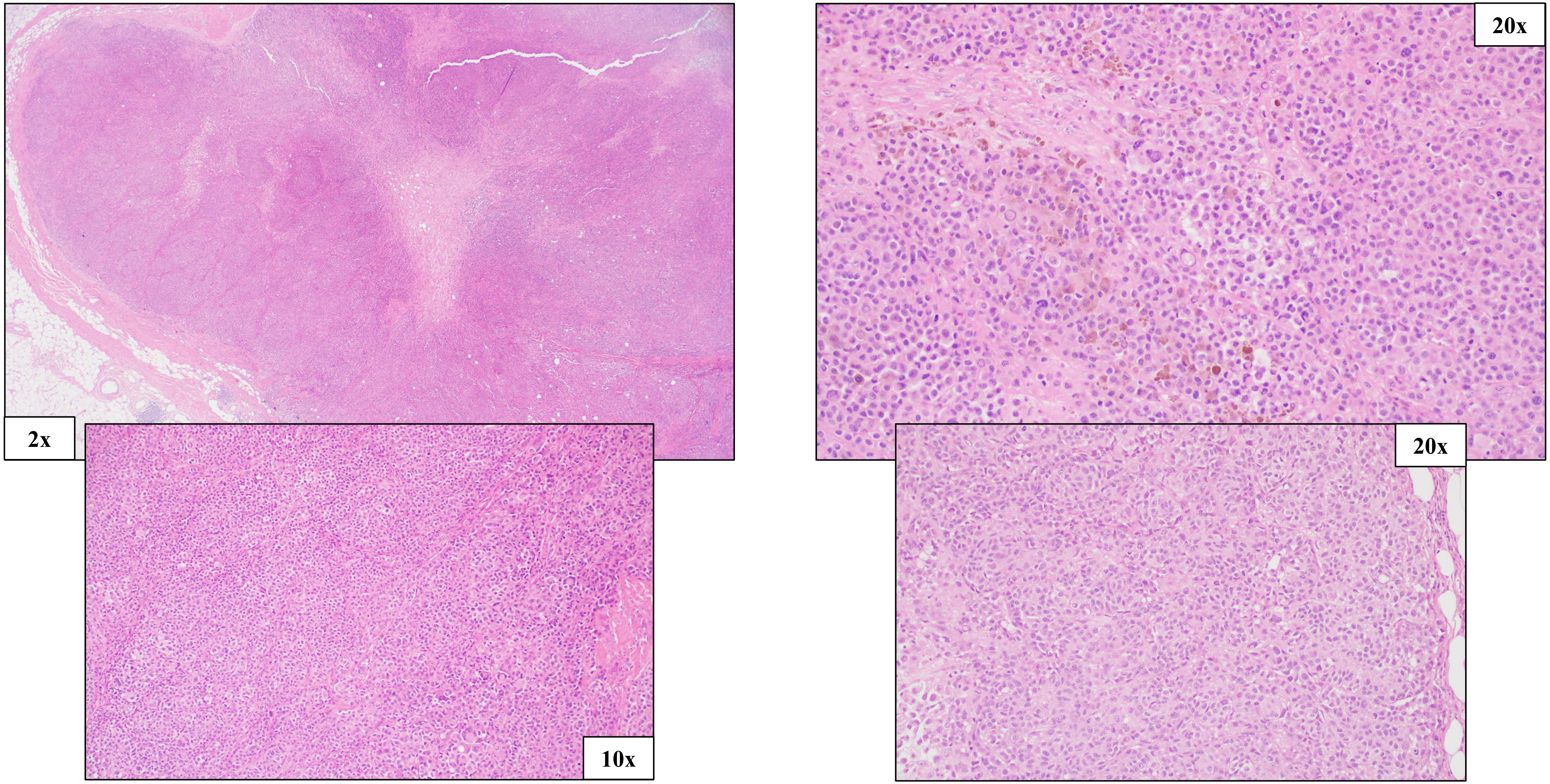


IHC Stains:

Not reported

BRAF G469A

### HCI-ASM084, Acral Melanoma

#### SNV (%VAF)

BRAF c.1406G>C, p.G469A (69%)

#### CNV

Amplification: TERT, EGFR, MET, BRAF
Loss: ARID1A, PTEN, SPRED1
Deep Deletion: CDKN2A

#### PDX Tumor Histology, Passages 2 and 4

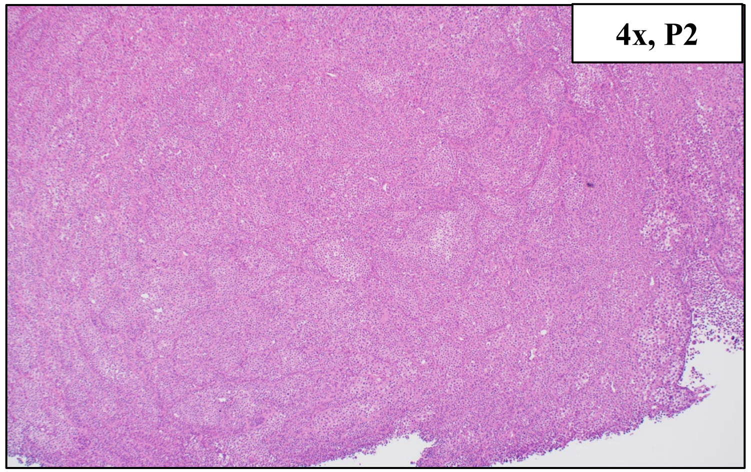


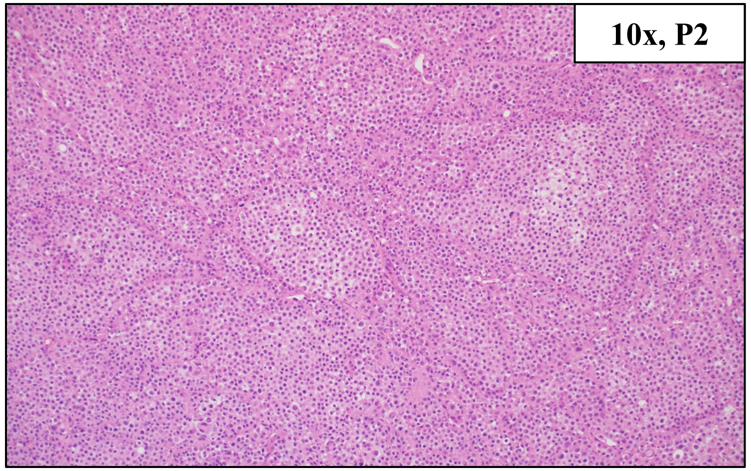


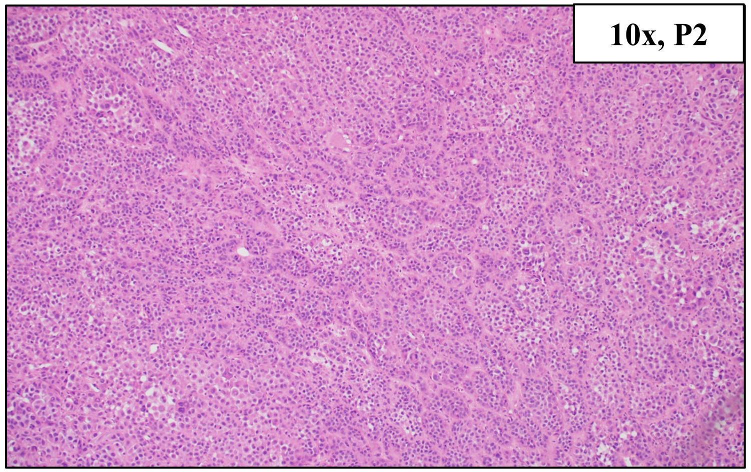


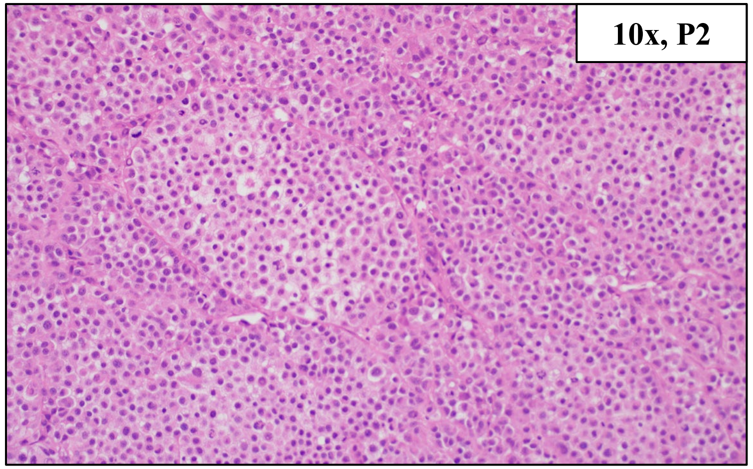


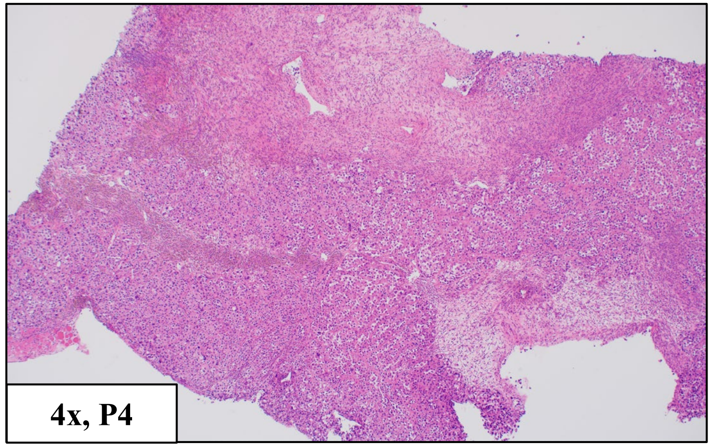


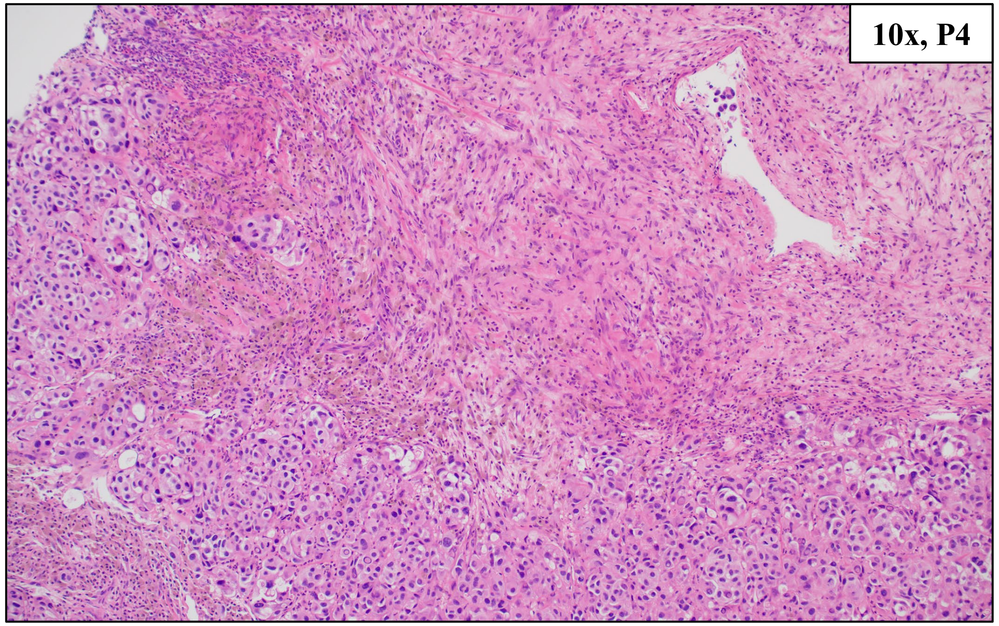


BRAF G469A

### HCI-AM085, Acral Melanoma

| Primary Tumor Location | PDX Tumor Location | Other Known Metastatic Locations |
| --- | --- | --- |
| Right Thumb Skin | Right Middle Lobe Lung | none |

Clinical Tumor Histology

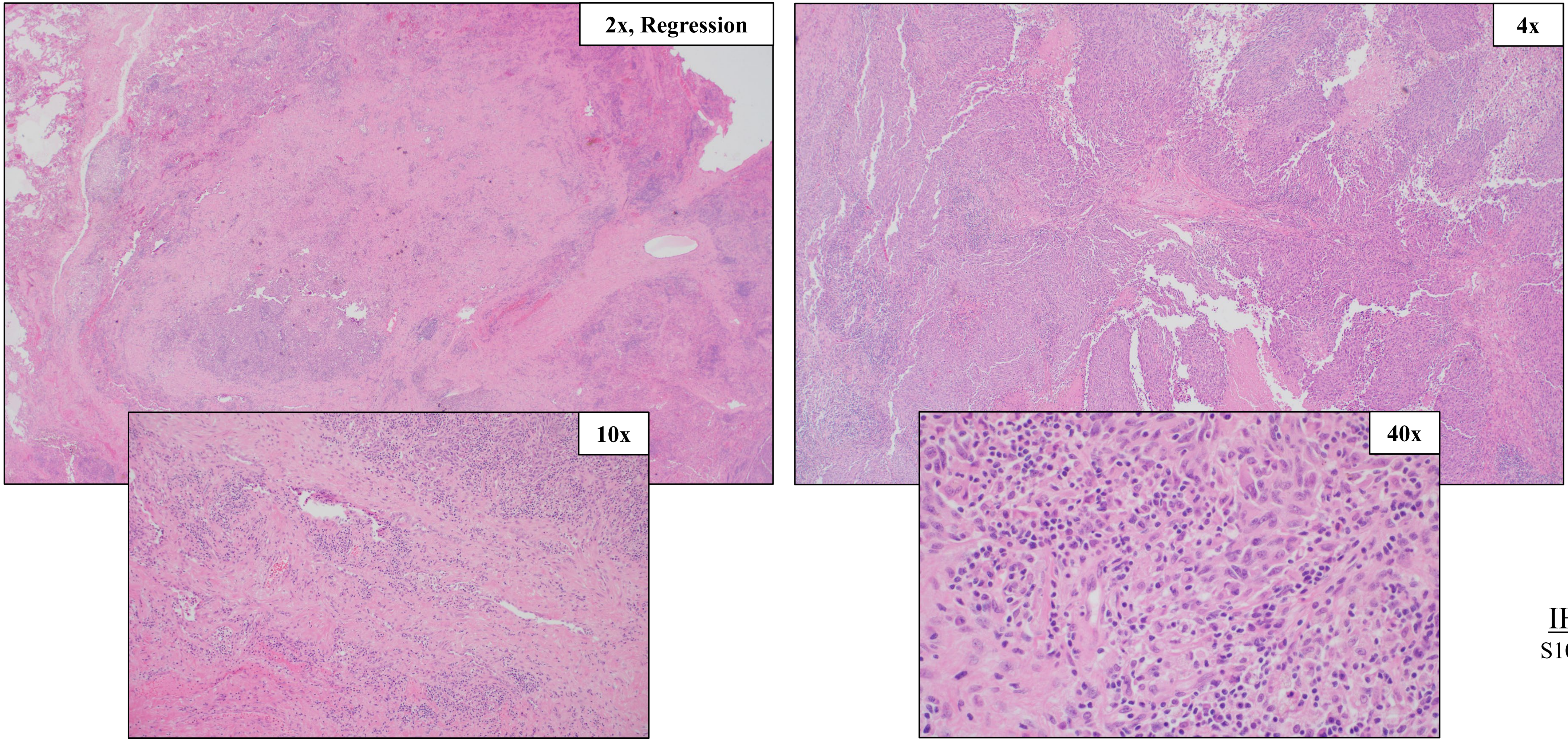


IHC Stains:
S100+, MelanA+

NF1 splice donor variant, High Amp: CDK4, MDM2, MC1R

### HCI-AM085, Acral Melanoma

<u>SNV (%VAF)</u>

NF1 Splice Donor Variant c.4763_4772+4del (99%)

<u>CNV</u>

Amplification ≥8 copies: CDK4, MDM2, MC1R
Amplification: MITF, TERT, MET, HRAS
Loss: ARID1B, PTEN, NF1

<u>PDX Tumor Histology, Passages 2 and 4</u>

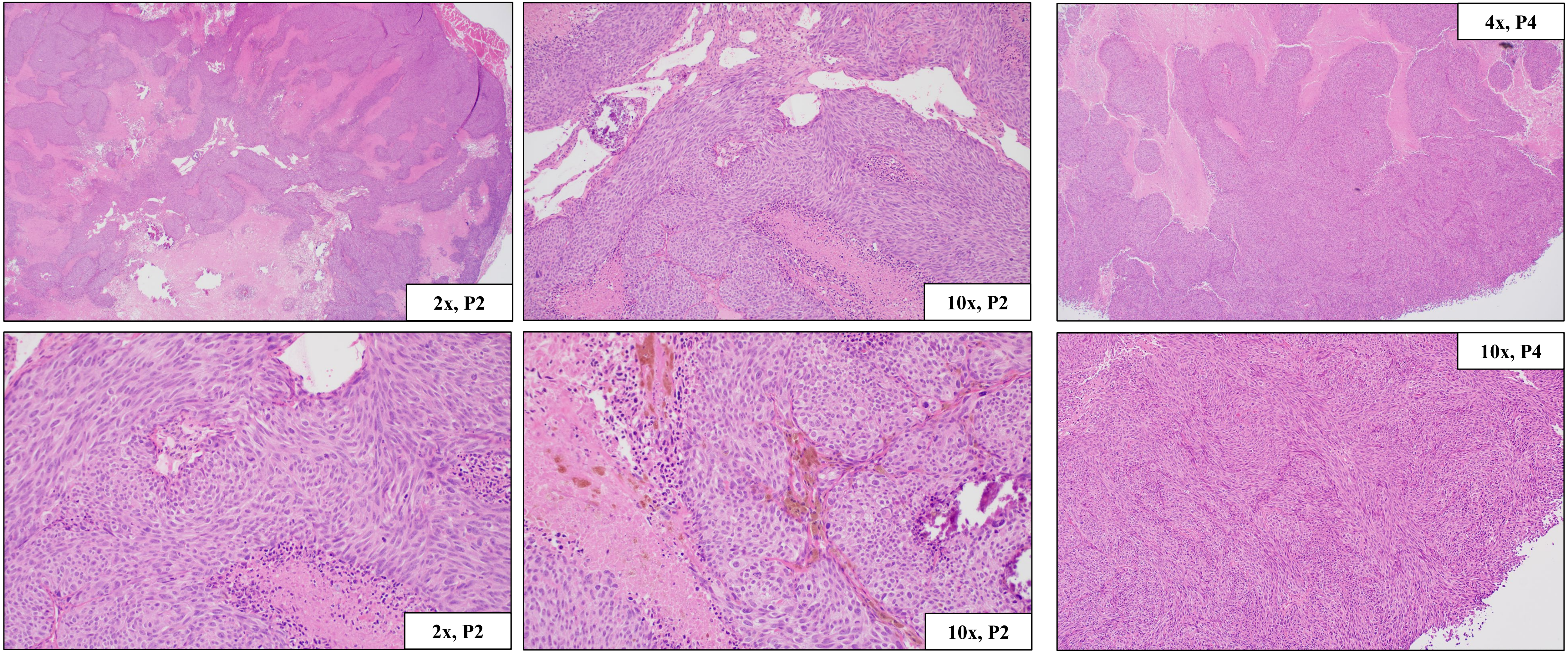


NF1 splice donor variant, High Amp: CDK4, MDM2, MC1R

### HCI-AM086, Acral Melanoma (same patient as HCI-AM087)

| Primary Tumor Location | PDX Tumor Location | Other Known Metastatic Locations |
| --- | --- | --- |
| Left Great Toe Skin | Left Groin Soft Tissue | Left Inguinal Lymph Nodes, Epidural Spine, C2 Vertebral Lamina, Bilateral Lungs |

Clinical Tumor Histology

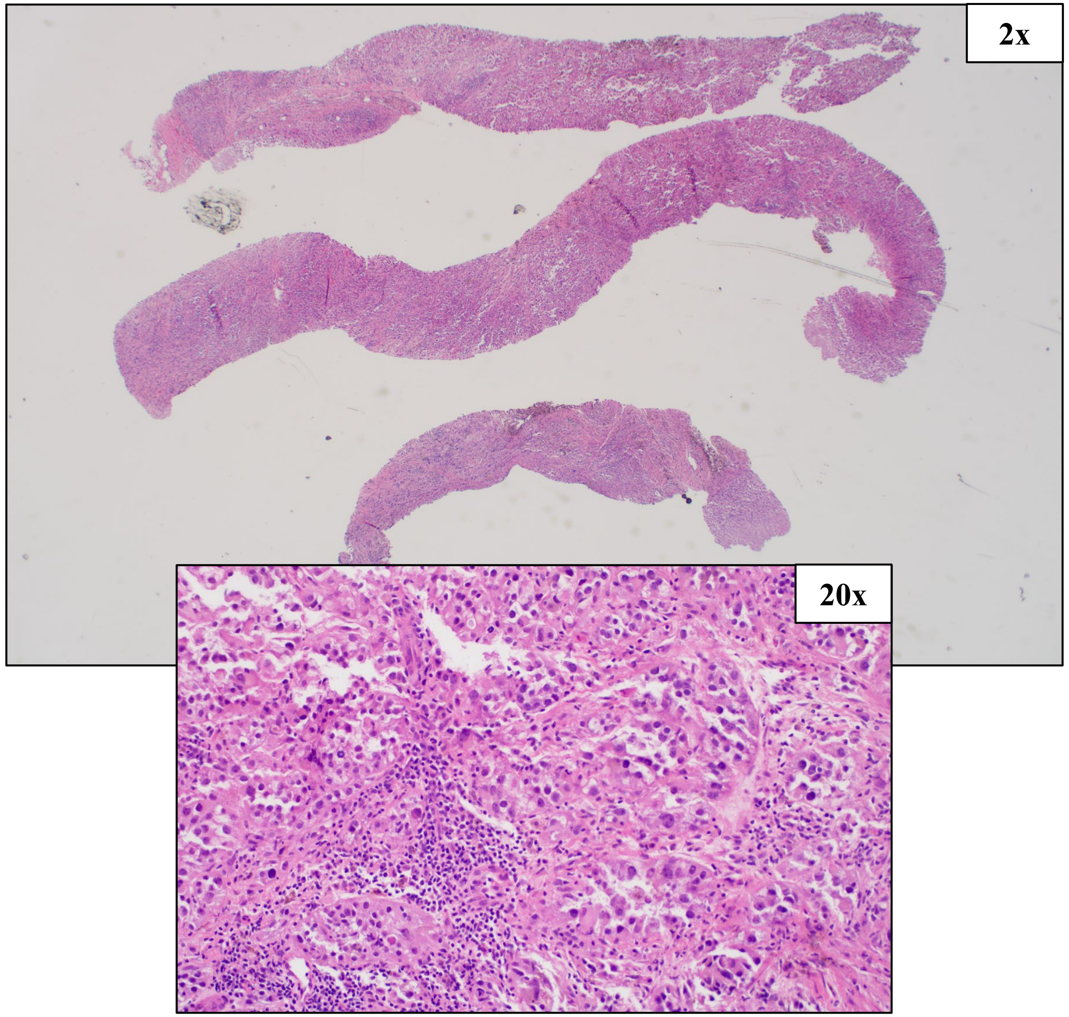


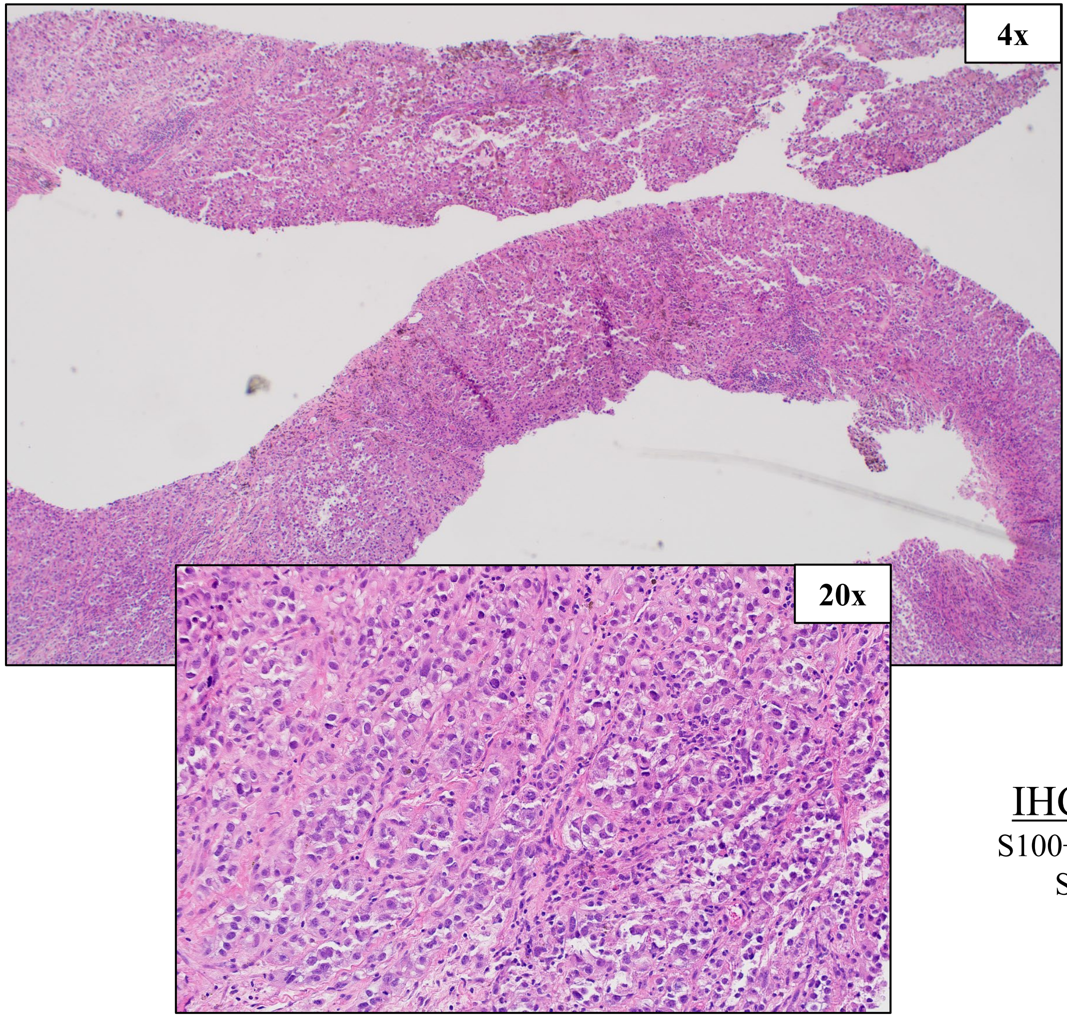


IHC Stains:
S100+, MelanA+, SOX10+

KRAS G12D, High Amp: CDK4, GAB2, MDM2

### HCI-AM086, Acral Melanoma (same patient as HCI-AM087)

#### SNV (%VAF)

KRAS c.35G>A, p.G12D (83%)

#### CNV

Amplification ≥8 copies: GAB2, CDK4, MDM2
Amplification: MITF, HRAS, KRAS
Loss: CDKN2A, ARID2

#### PDX Tumor Histology, Passages 2 and 4

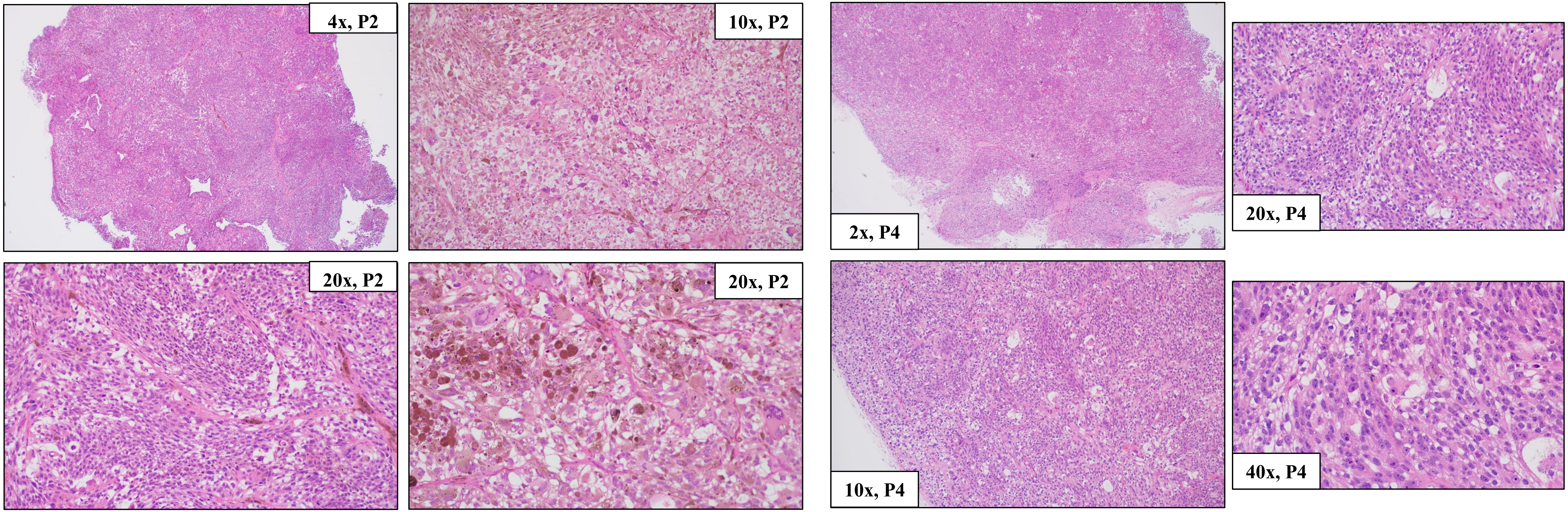


KRAS G12D, High Amp: CDK4, GAB2, MDM2

### HCI-AM087, Acral Melanoma (same patient as HCI-AM086)

| Primary Tumor Location | PDX Tumor Location | Other Known Metastatic Locations |
|---|---|---|
| Left Great Toe Skin | Left Groin Soft Tissue | Left Inguinal Lymph Nodes, Epidural Spine, C2 Vertebral Lamina, Bilateral Lungs |

Clinical Tumor Histology

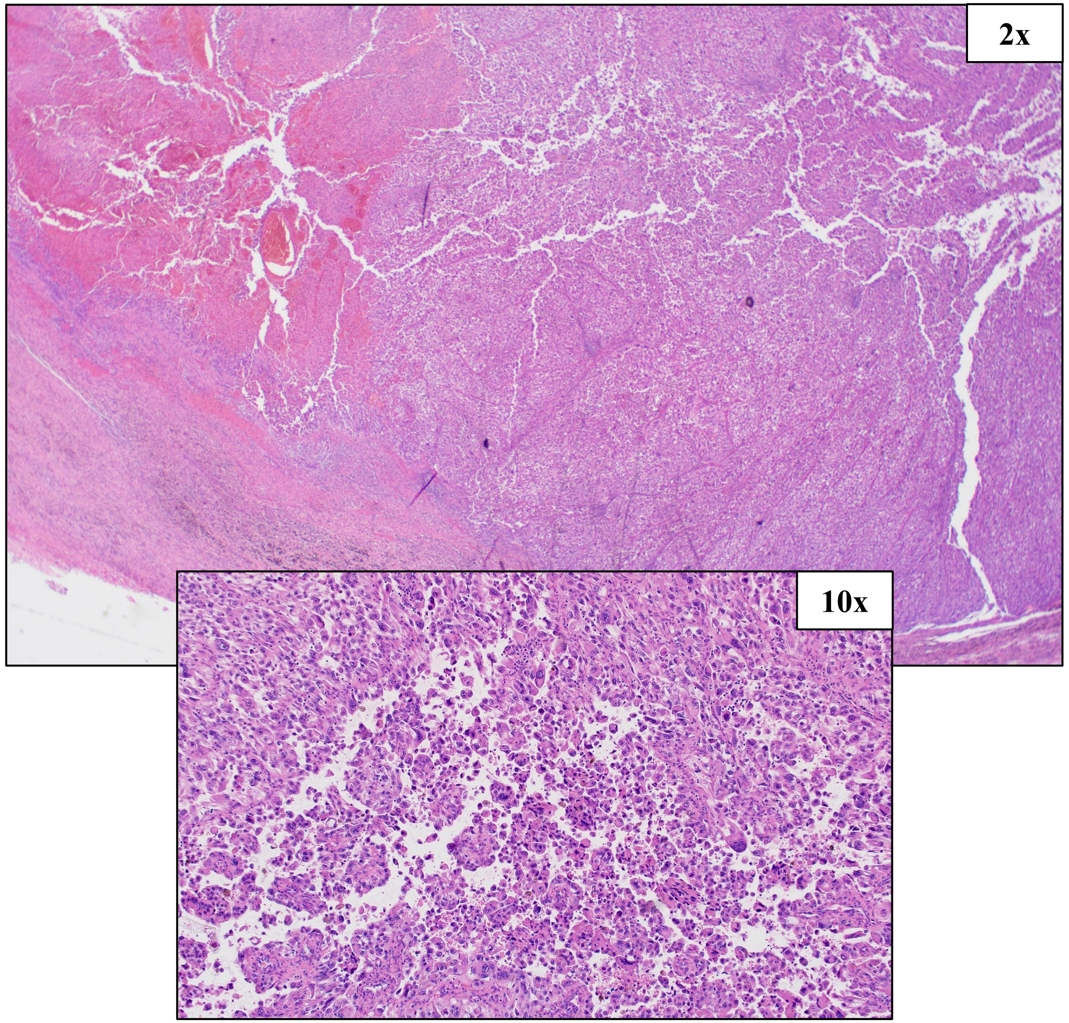


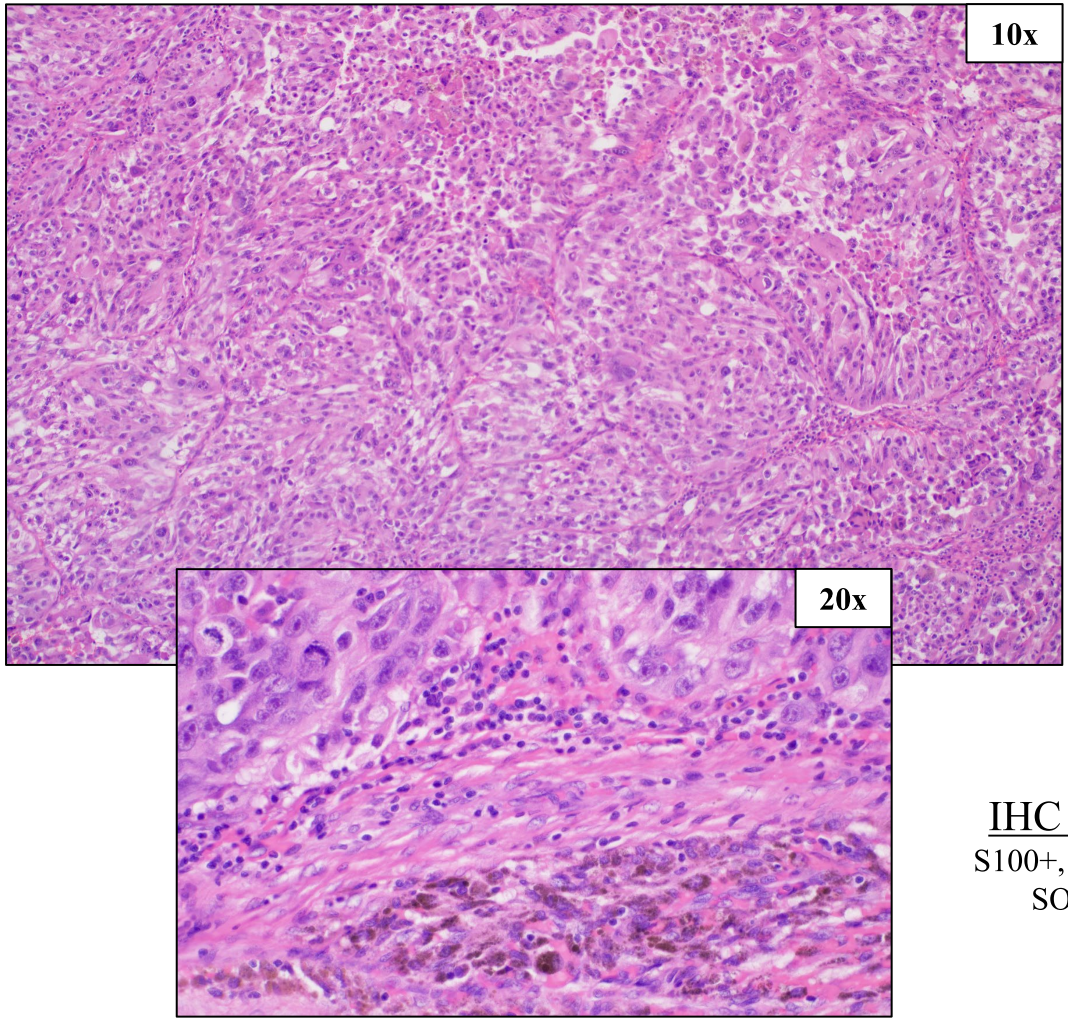


IHC Stains:
S100+, MelanA+, SOX10+

KRAS G12D, High Amp: CDK4, GAB2, MDM2

### HCI-AM087, Acral Melanoma (same patient as HCI-AM086)

PDX Tumor Histology, Passages 2 and 5

SNV (%VAF)

KRAS c.35G>A, p.G12D (74%)

CNV

Amplification ≥8 copies: GAB2, CDK4, MDM2
Amplification: MET, KRAS, BRCA1
Loss: PDGFRA, KIT, ARID1B, CDKN3A, ARID2

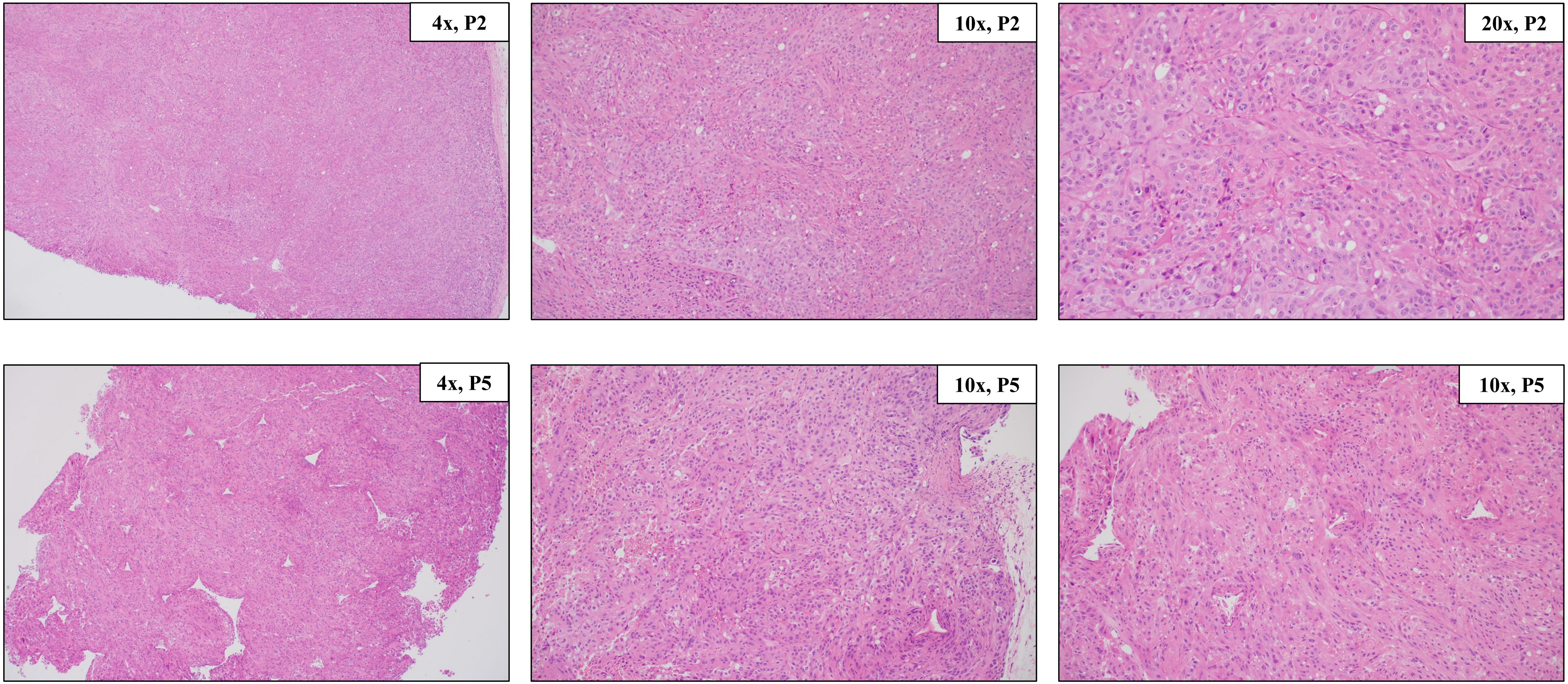


KRAS G12D, High Amp: CDK4, GAB2, MDM2

### HCI-AM088, Acral Melanoma

| Primary Tumor Location | PDX Tumor Location | Other Known Metastatic Locations |
|---|---|---|
| Right Foot Acral Skin | R Groin Lymph Node | Right Inguinal LN, R Humerus, T1-2 Vertebral Bodies, C1-C5 Epidural Soft Tissue, Brain |

Clinical Tumor Histology

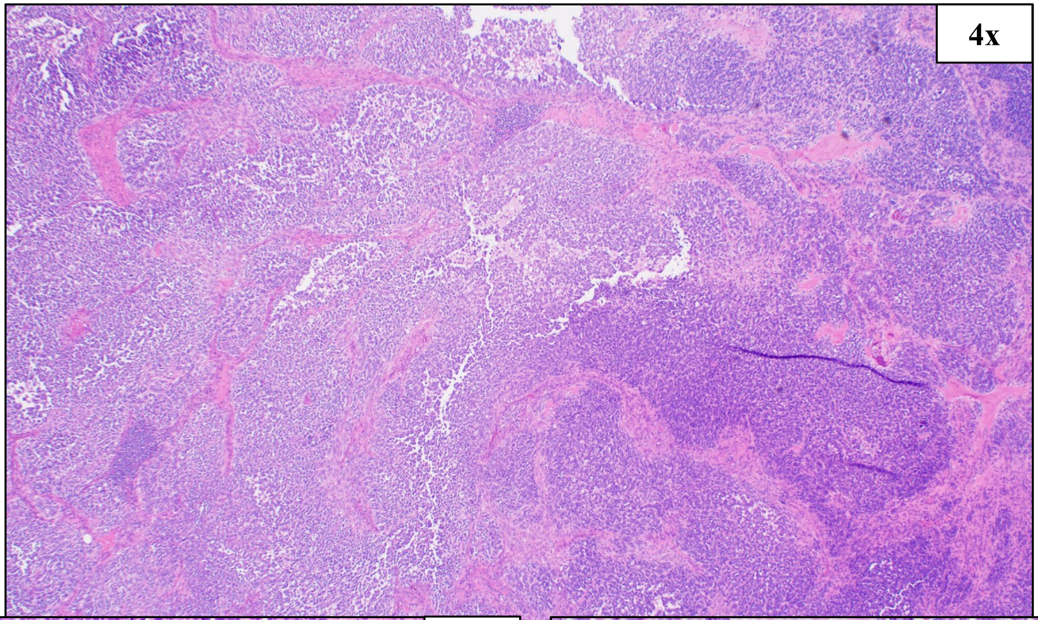


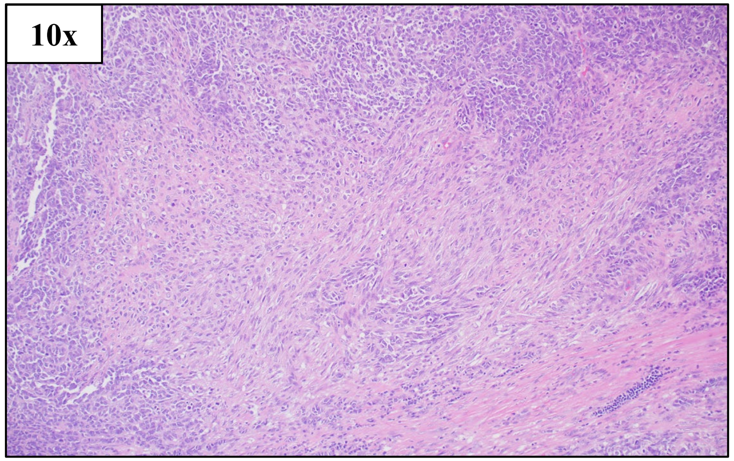


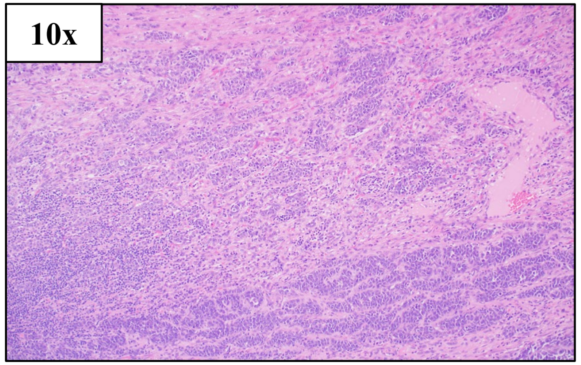


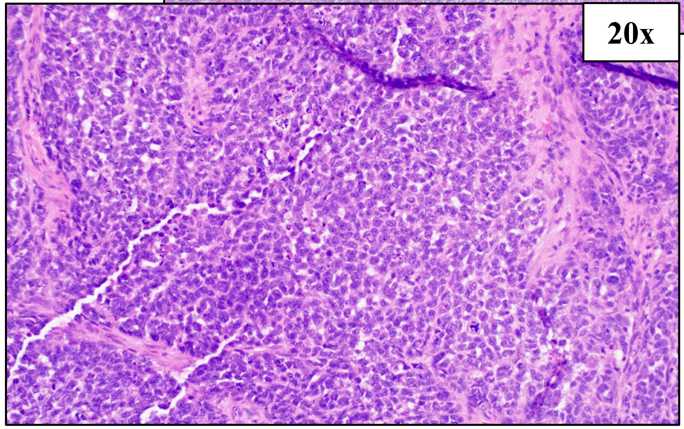


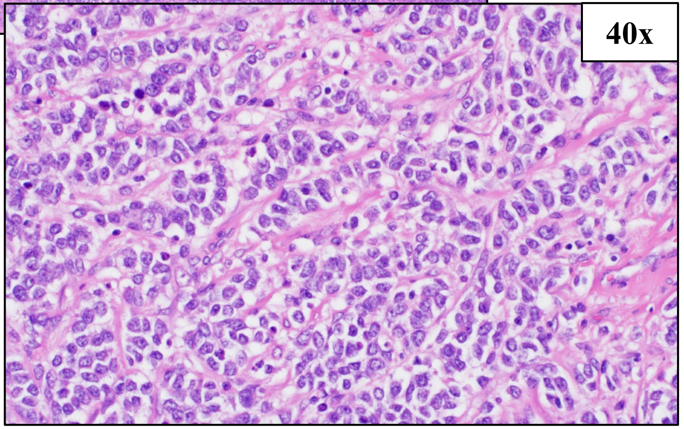


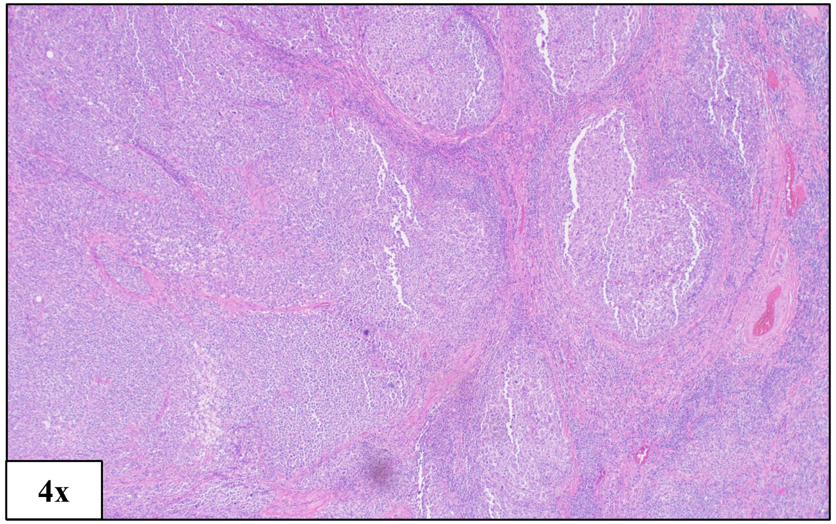


IHC Stains:

S100+/-, MelanA+

BRAF V600E

### HCI-AM088, Acral Melanoma

#### SNV (%VAF)

BRAF c.1799T>A p.V600E (50%)
PTEN c.697C>T p.R233* (97%)
PIK3CA c.1035T>A p.N345K (Clinical NGS)

#### CNV

Amplification: HRAS, MC1R
Loss: PTEN, BRCA2, RB1

PDX Tumor Histology, Passage 2 and 3

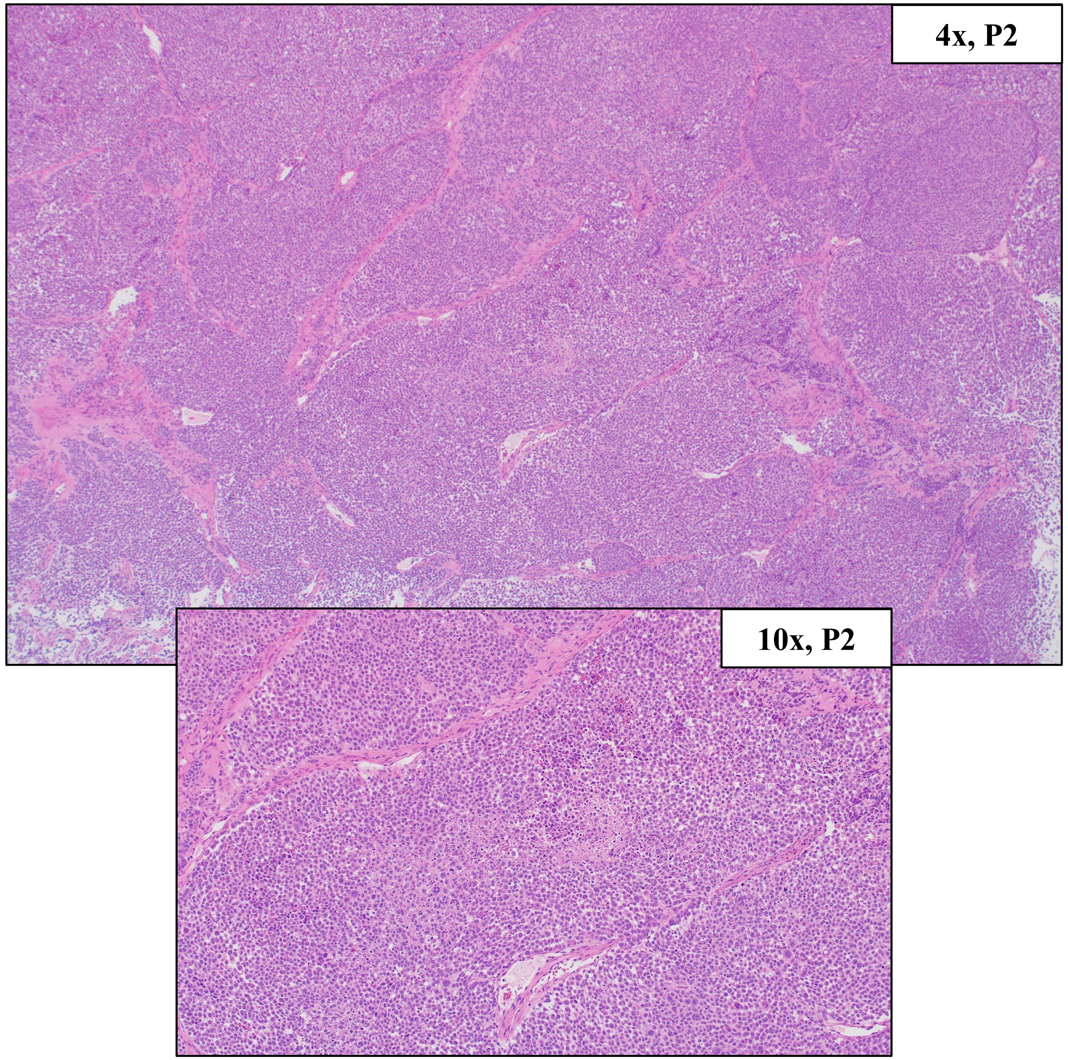


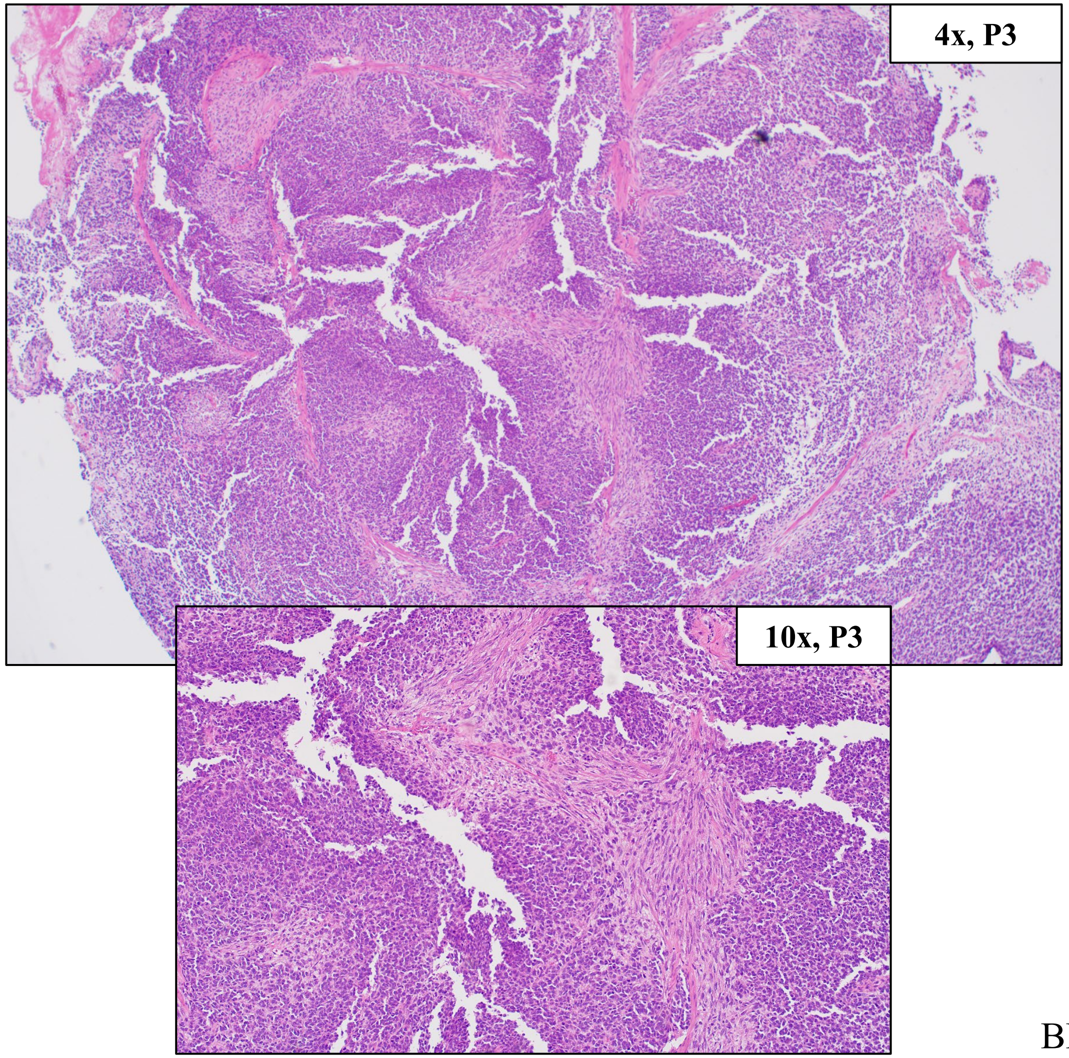


BRAF V600E

### HCI-AM090, Acral Melanoma

| Primary Tumor Location | PDX Tumor Location | Other Known Metastatic Locations |
| --- | --- | --- |
| Left 4th Dorsal Toe and Nail Bed | Primary Tumor | Left Groin Lymph Node |

Clinical Tumor Histology

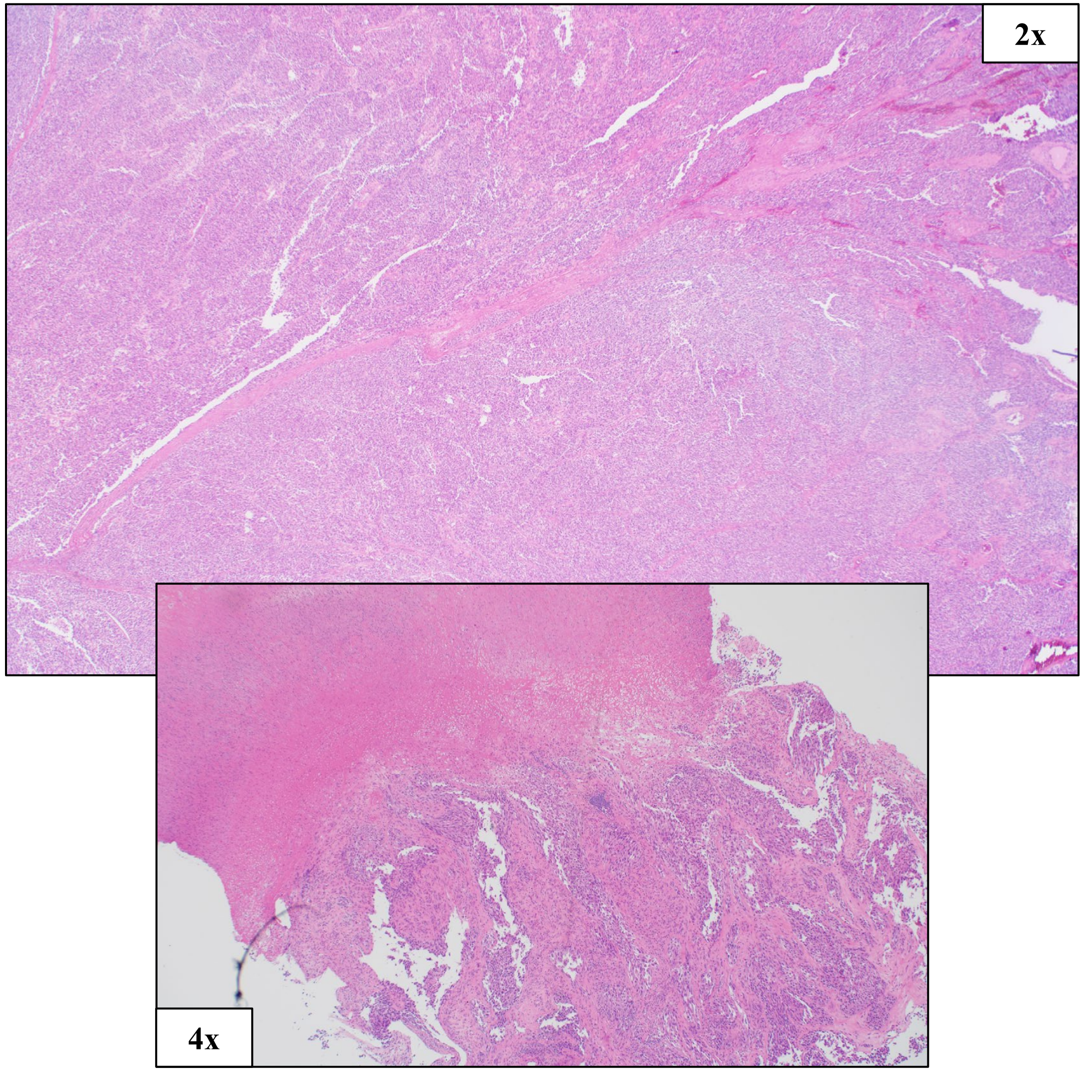


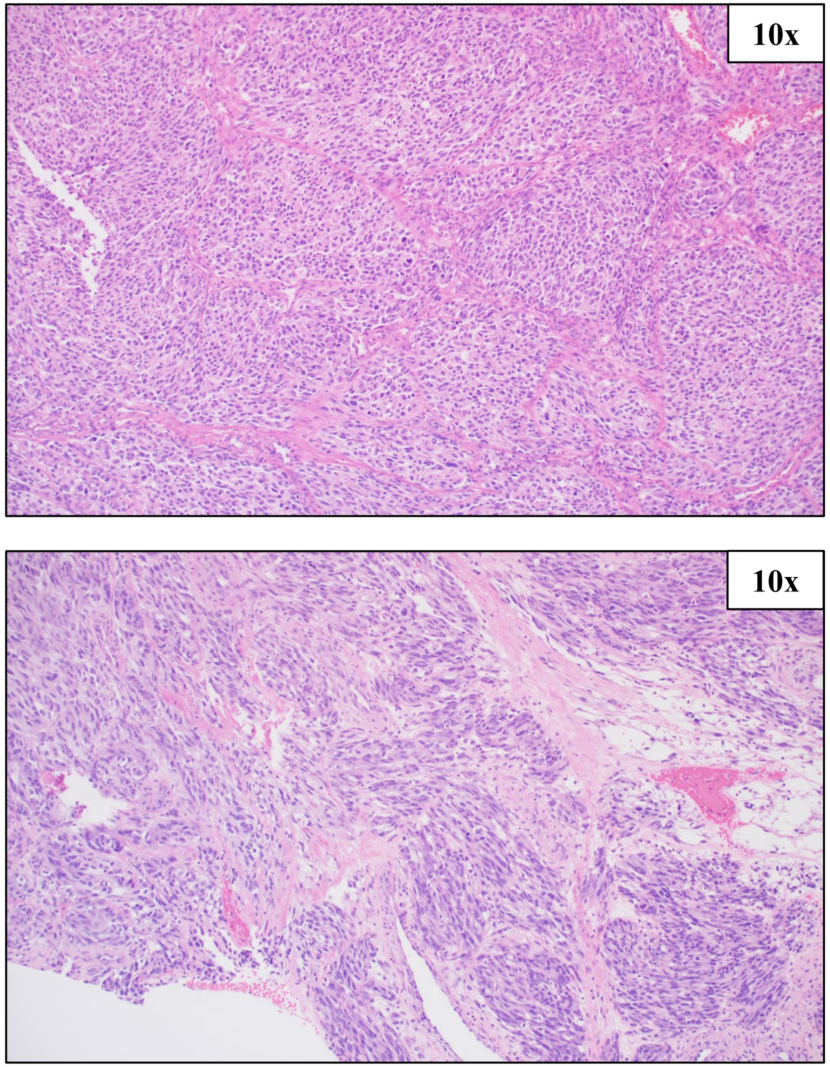


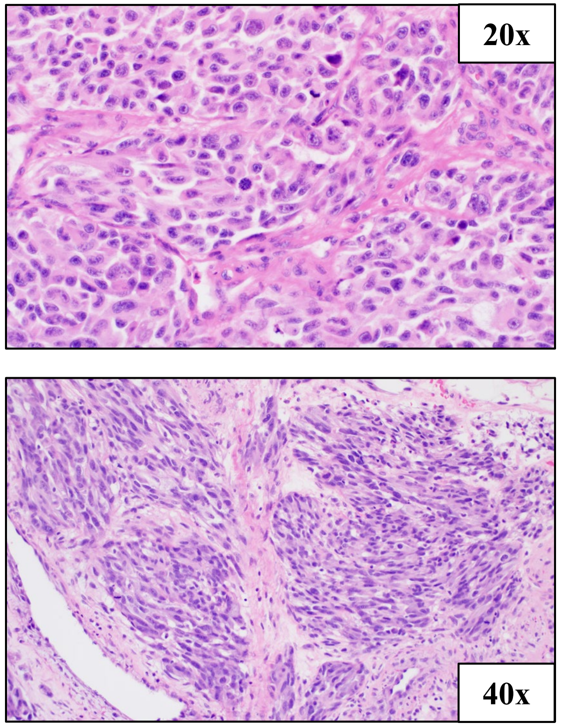


IHC Stains:

S100+, MART1+, SOX10+

High Amp: CRKL

### HCI-AM090, Acral Melanoma

#### SNV (%VAF)

MAP2K1 c.362G>C, p.C121S (66%)

#### CNV

Amplification ≥8 copies: CRKL
Amplification: TERT, HRAS, CCND1, CDK4
Loss: ARID1B, CDKN2A, PTEN, BRCA2, RB1, SPRED1

#### PDX Tumor Histology, Passages 1 and 3

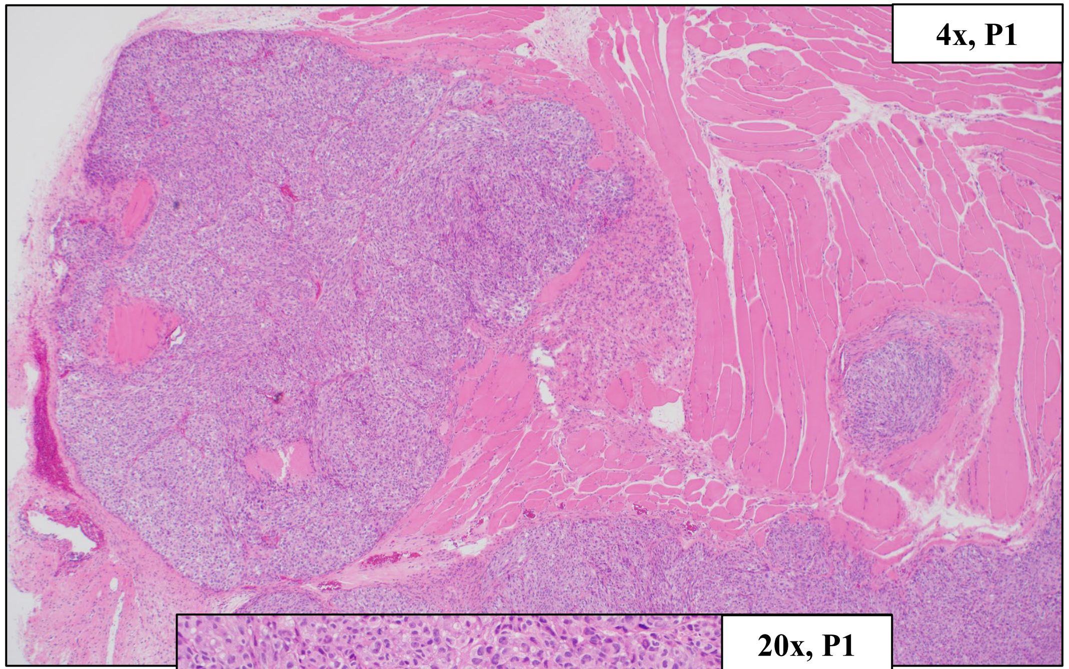


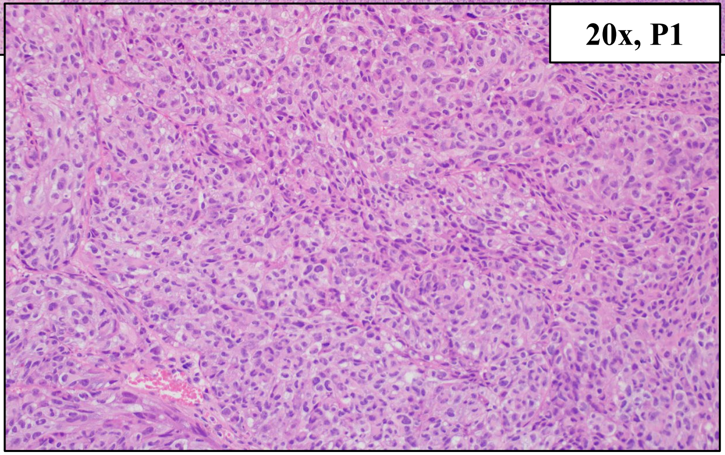


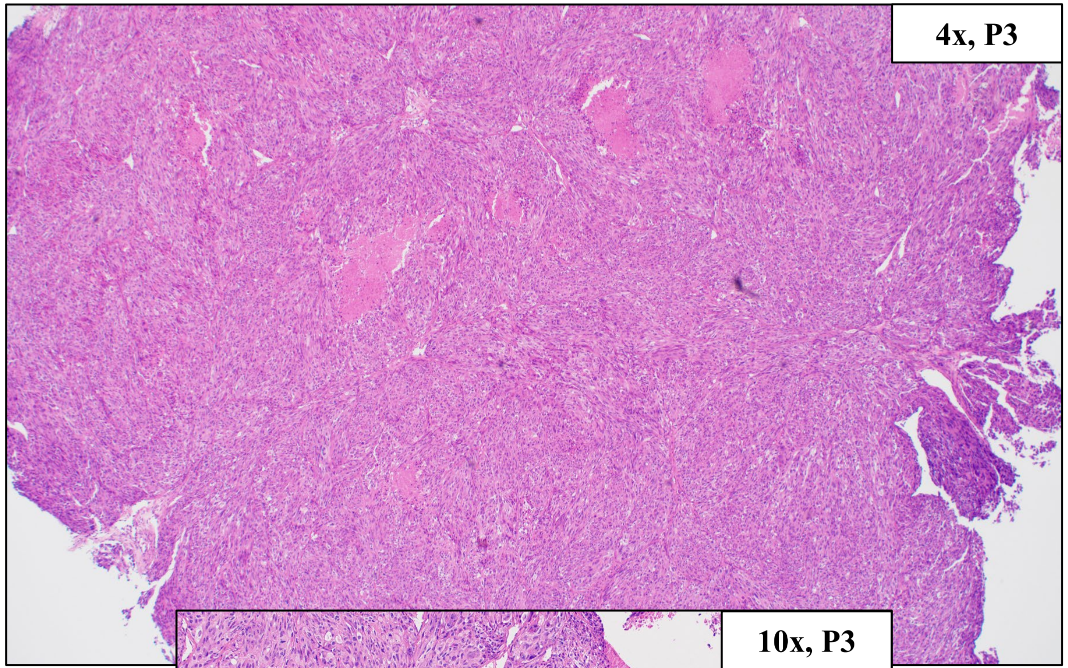


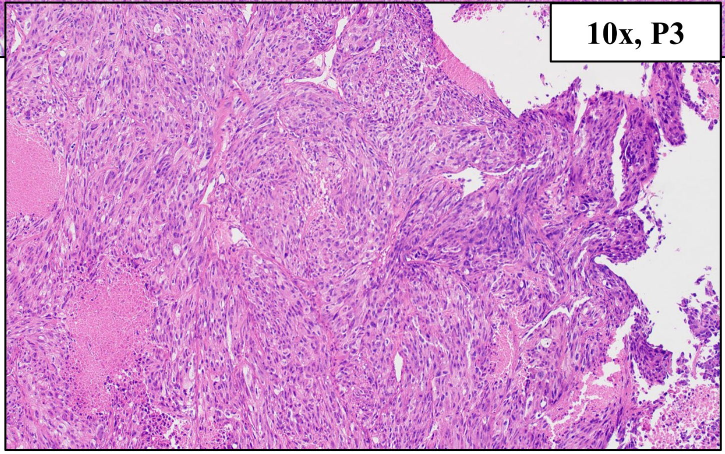


High Amp: CRKL

### HCI-ASM091, Acral Melanoma

| Primary Tumor Location | PDX Tumor Location | Other Known Metastatic Locations |
| --- | --- | --- |
| Left Thumbnail | Primary Tumor | None |

Clinical Tumor Histology

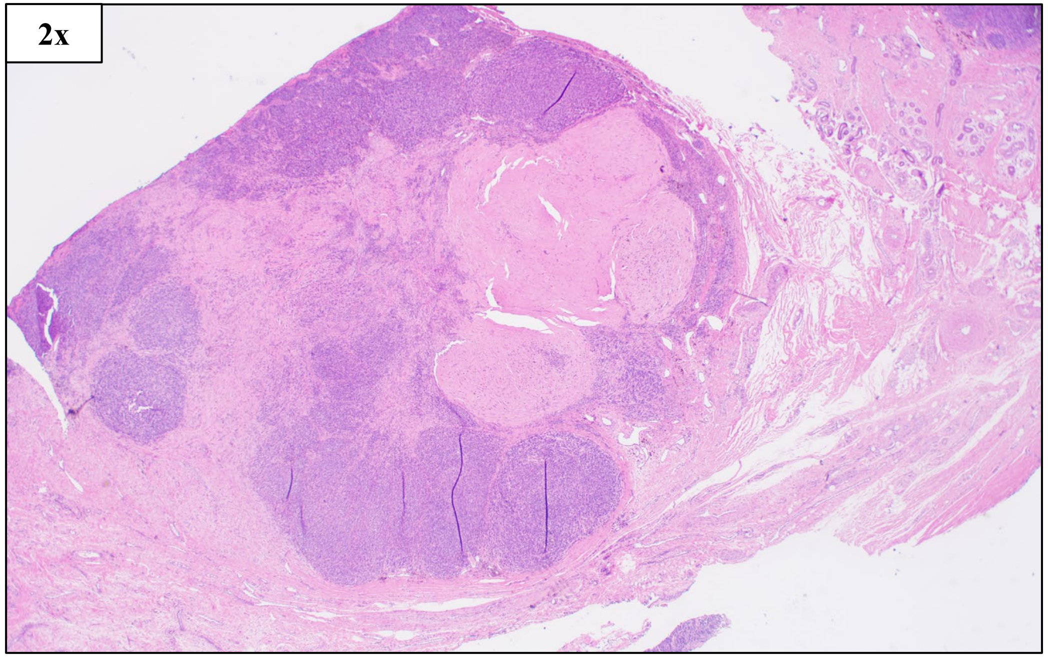


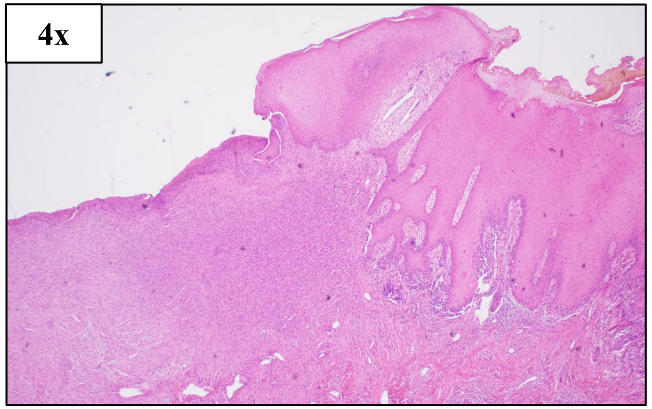


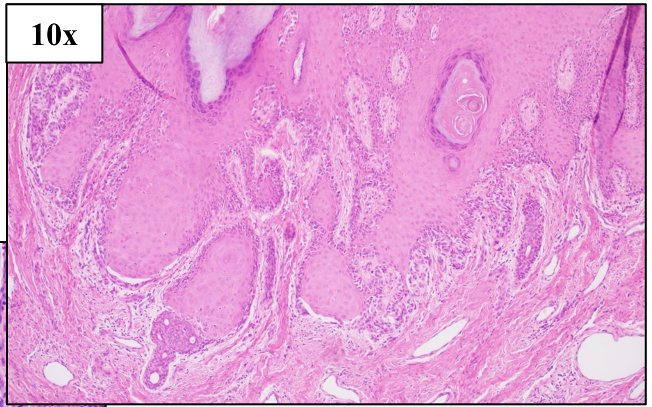


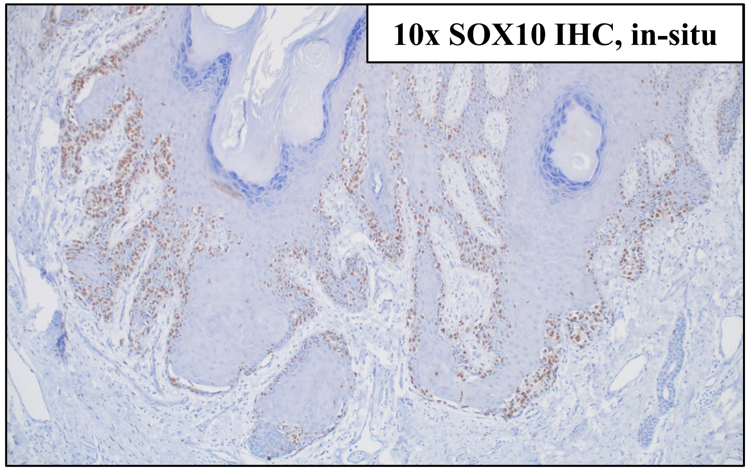


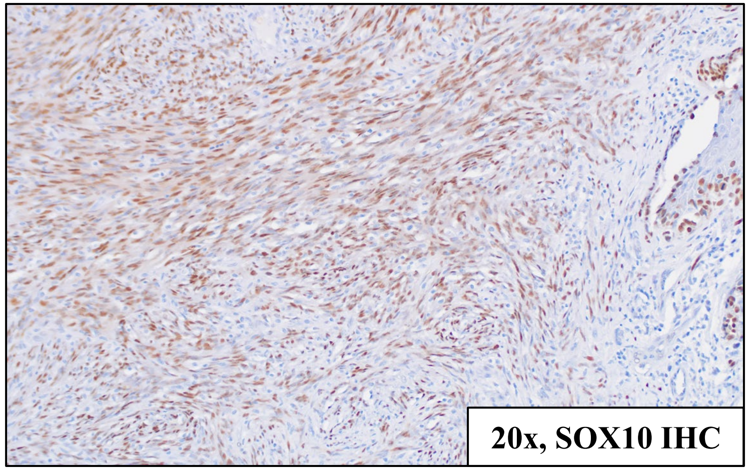


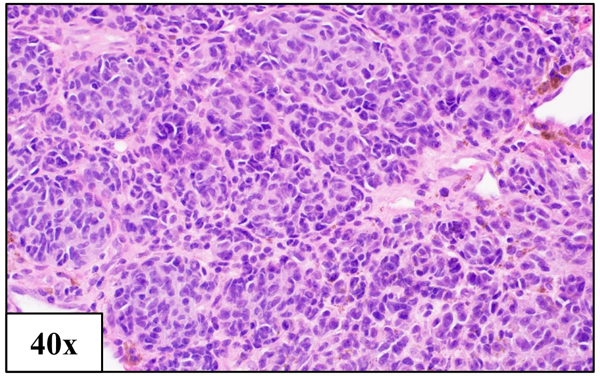


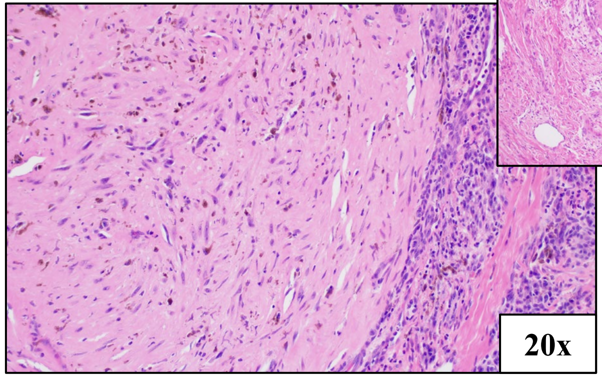


IHC Stains:
SOX10+

High Amp: GAB2

### HCI-ASM091, Acral Melanoma

#### SNV (%VAF)

No Tier 1 or 2 mutations

#### CNV

Amplification ≥8 copies: GAB2
Amplification: HRAS, CCND1, BRCA1
Loss: ARID1B
Deep deletion: CDKN2A

#### PDX Tumor Histology, Passages 1 and 2

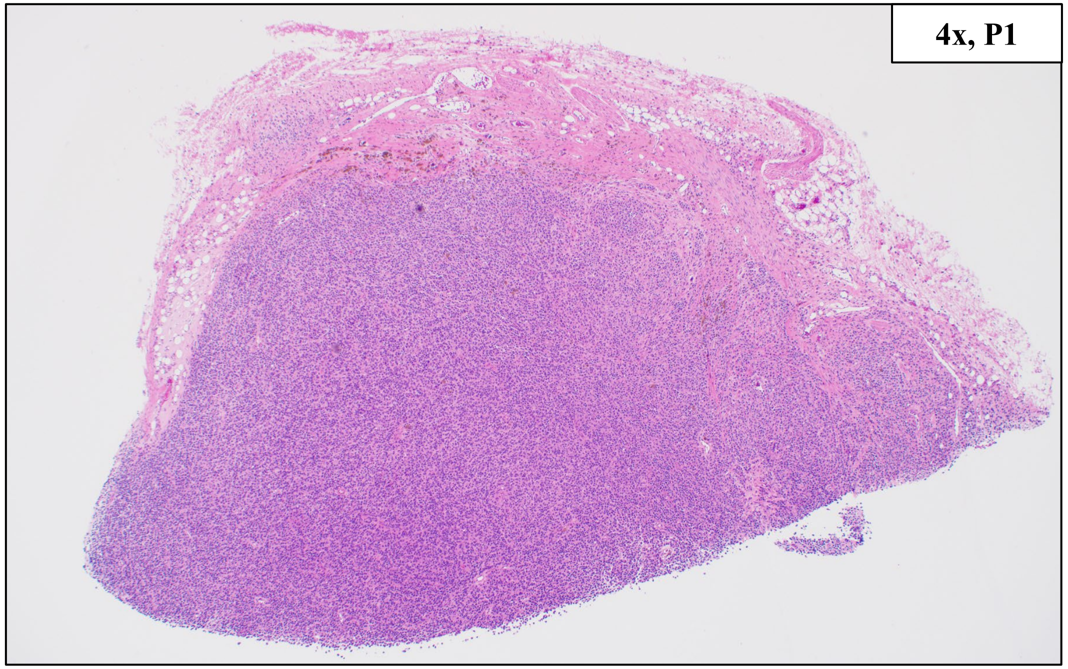


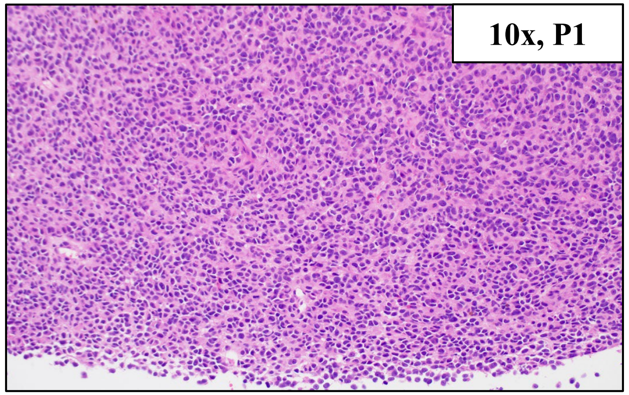


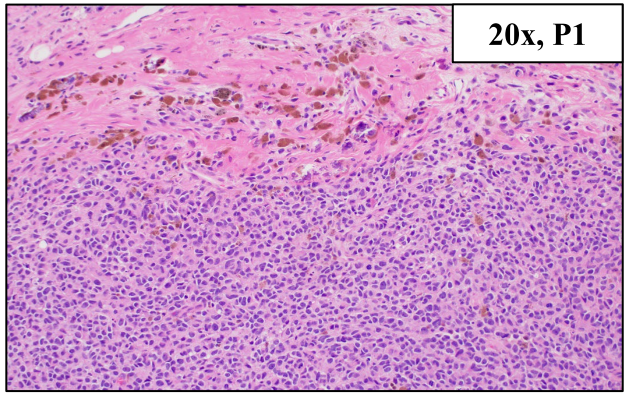


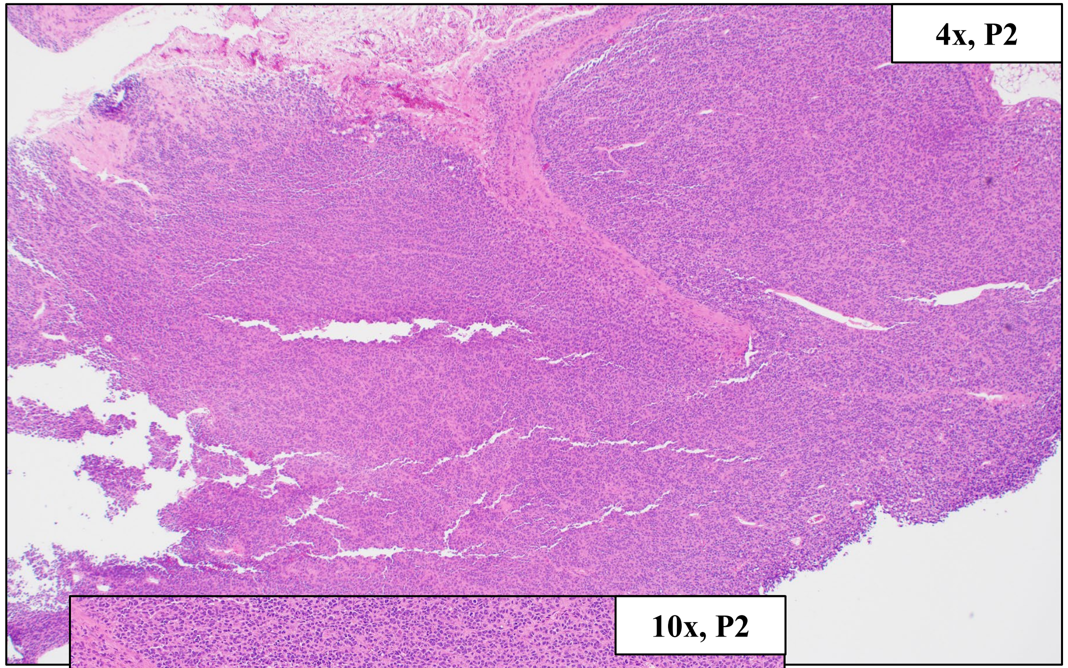


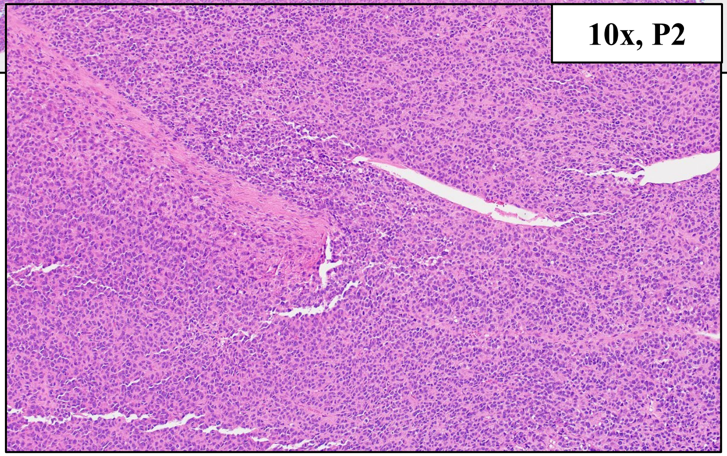


High Amp: GAB2

### HCI-AM092, Acral Melanoma

| Primary Tumor Location | PDX Tumor Location | Other Known Metastatic Locations |
| --- | --- | --- |
| Right Lateral Ankle at Acral Skin Junction | Primary Tumor | Right Inguinal and Obturator Lymph Nodes |

#### Clinical Tumor Histology

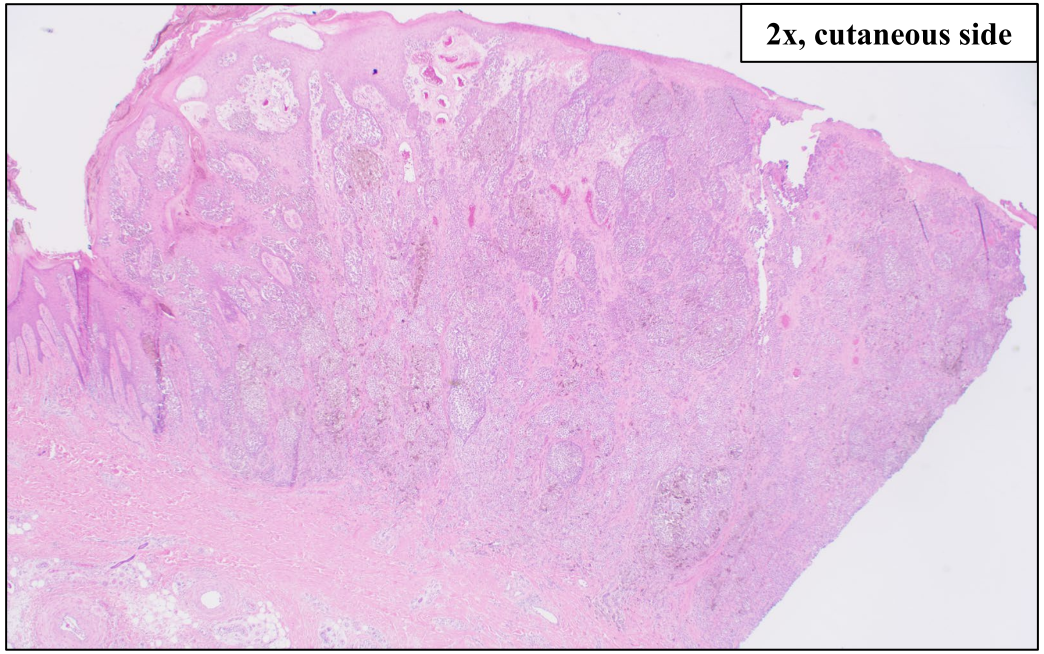
2x, cutaneous side

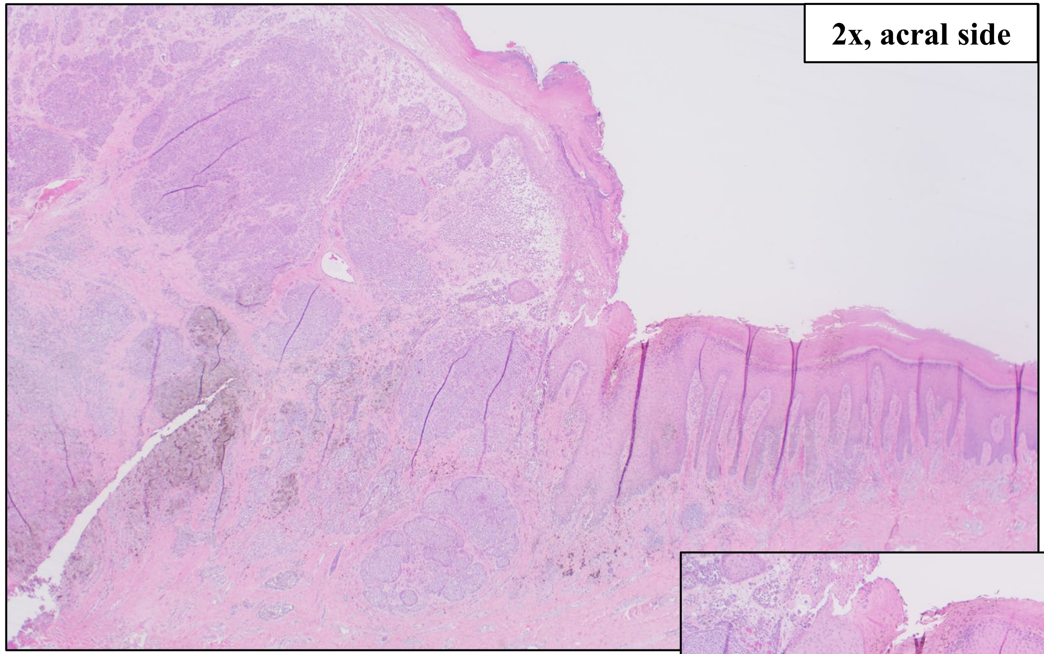
2x, acral side

**IHC Stains:**
SOX10+, MelanA+

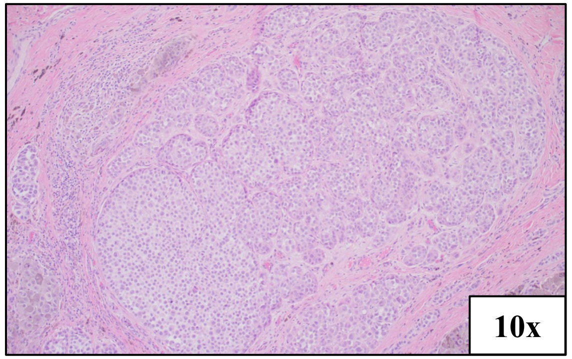
10x

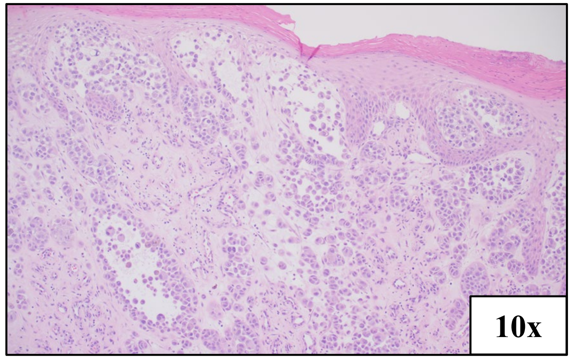
10x

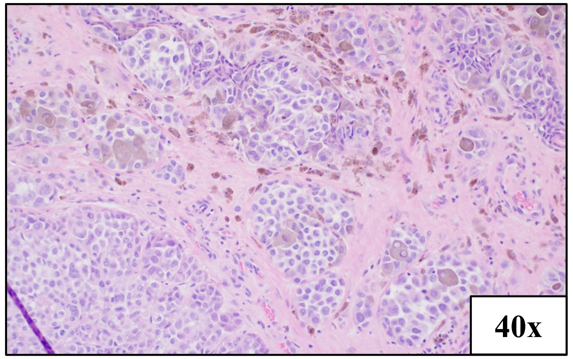
40x

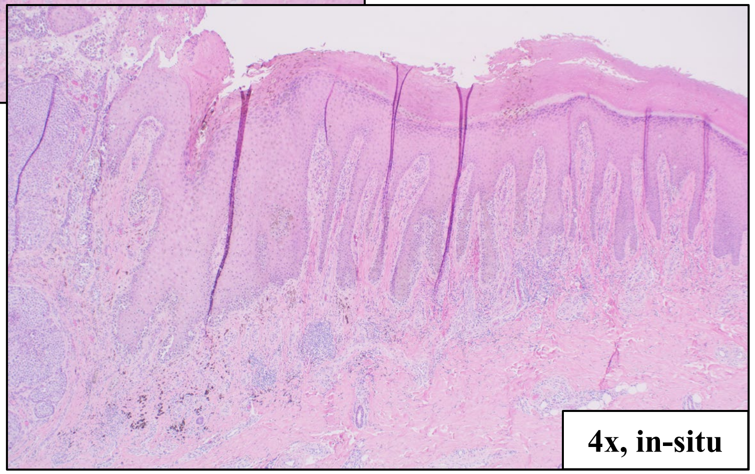
4x, in-situ

BRAF V600E

### HCI-AM092, Acral Melanoma

#### SNV (%VAF)

BRAF c.1799T>A, p.V660E (65%)
CDKN2A c.301G>T, p.G101W (98%)

#### CNV

Loss: ARID1B, CDKN2A, HRAS, TP53
Deep deletion: PTEN

#### PDX Tumor Histology

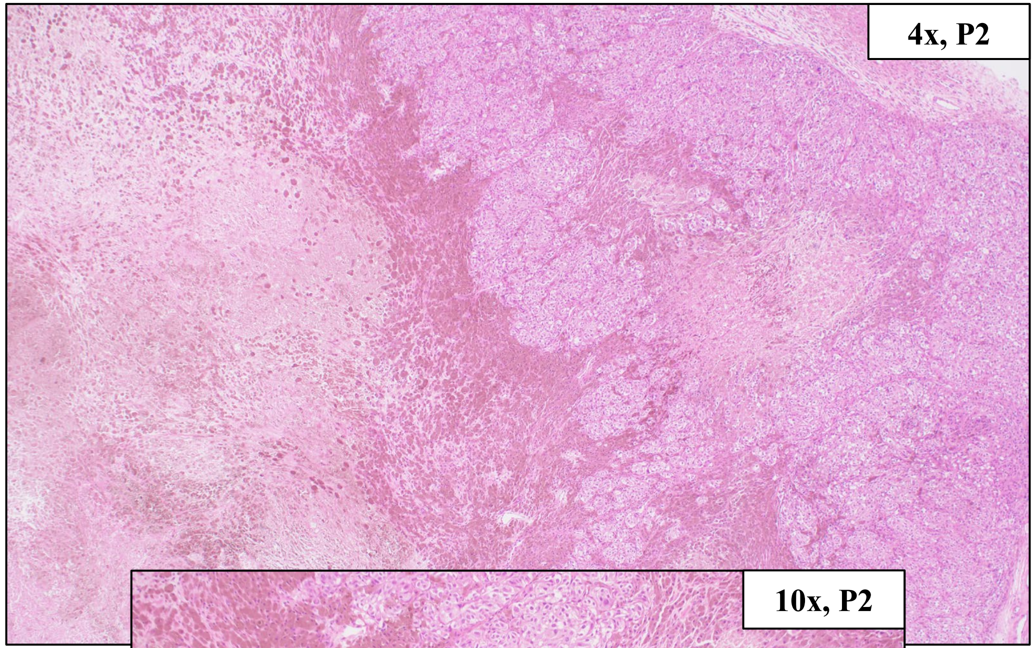


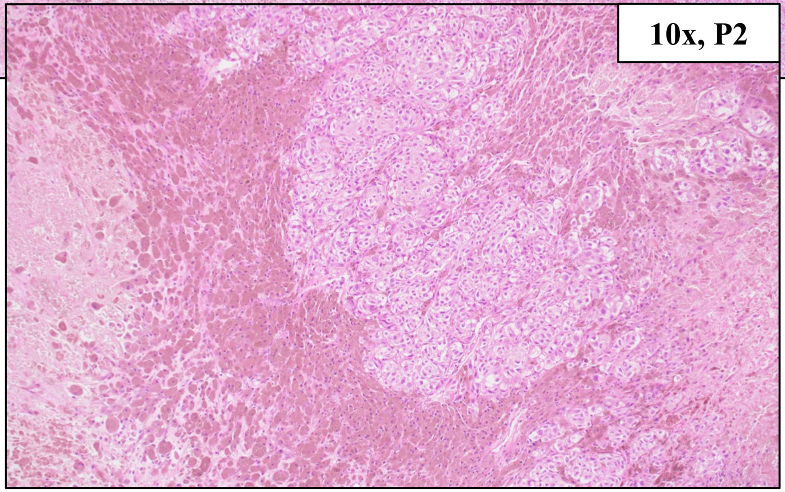


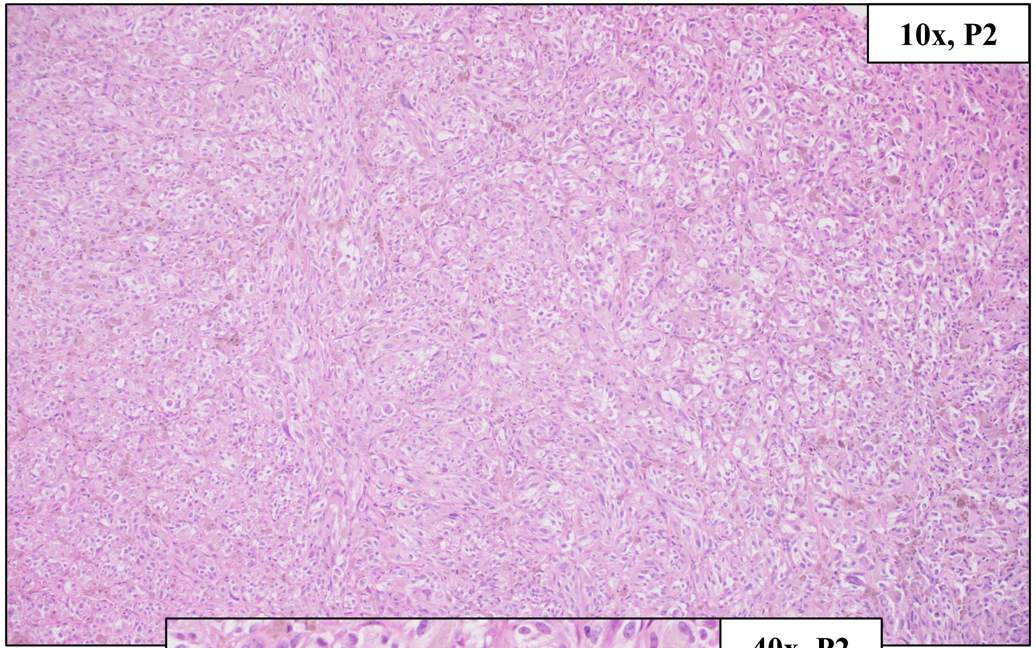


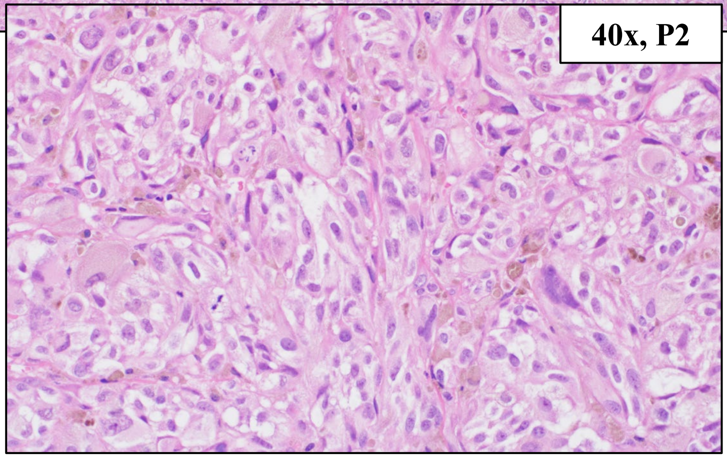


BRAF V600E

### HCI-AM093, Acral Melanoma

| Primary Tumor Location | PDX Tumor Location | Other Known Metastatic Locations |
|---|---|---|
| Left 5th Toe, Lateral Volar Skin | Primary Tumor | Left Medial Calf Skin and Lymph Nodes, Peritoneal Carcinomatosis |

Clinical Tumor Histology

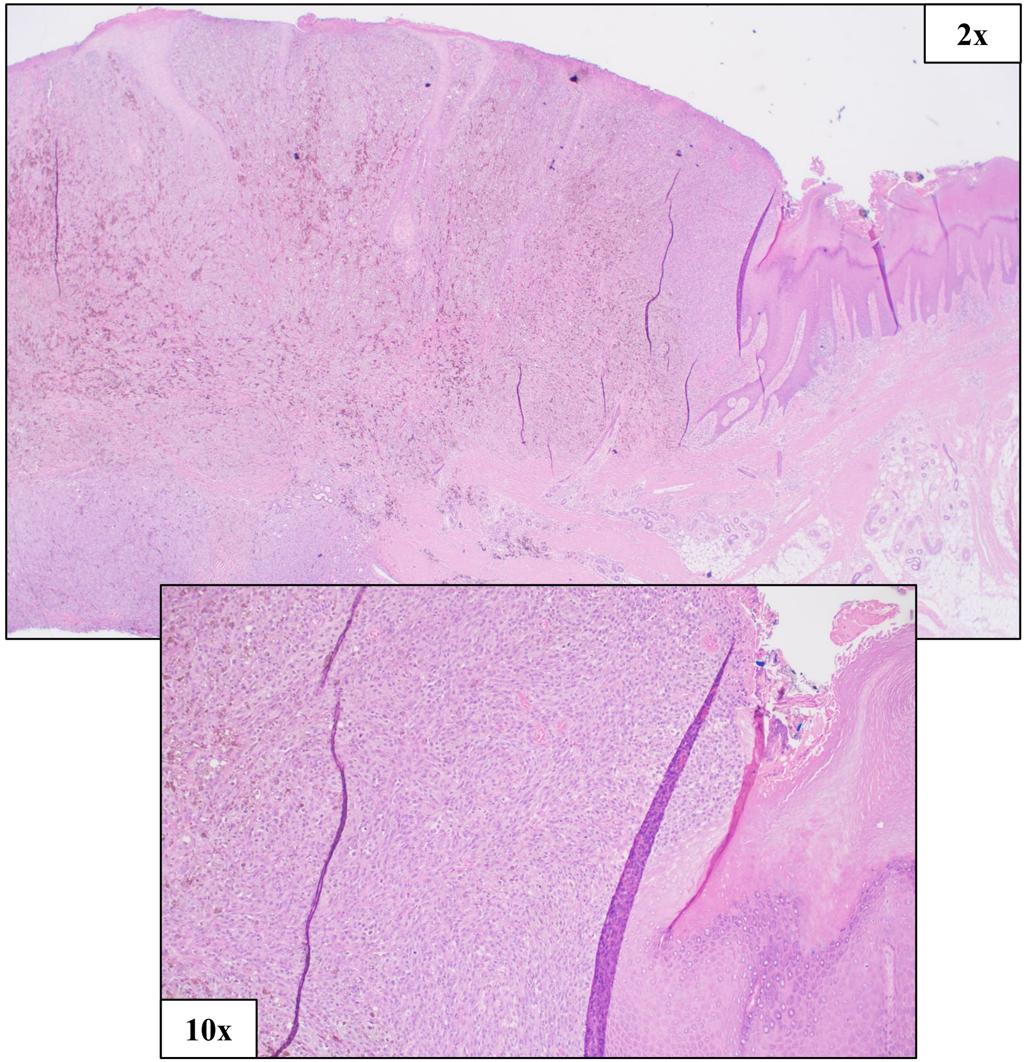


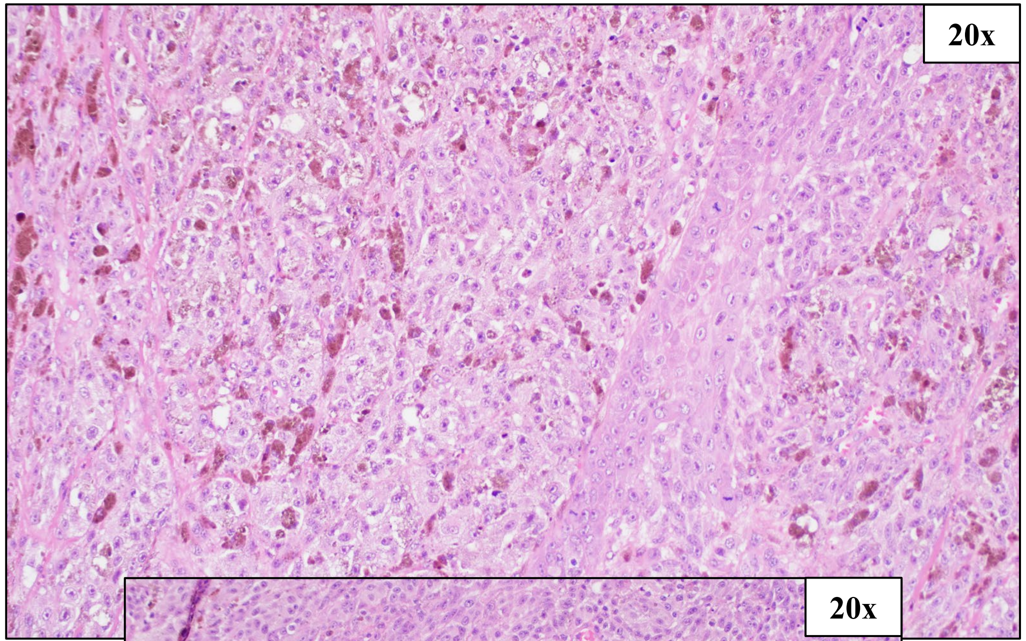


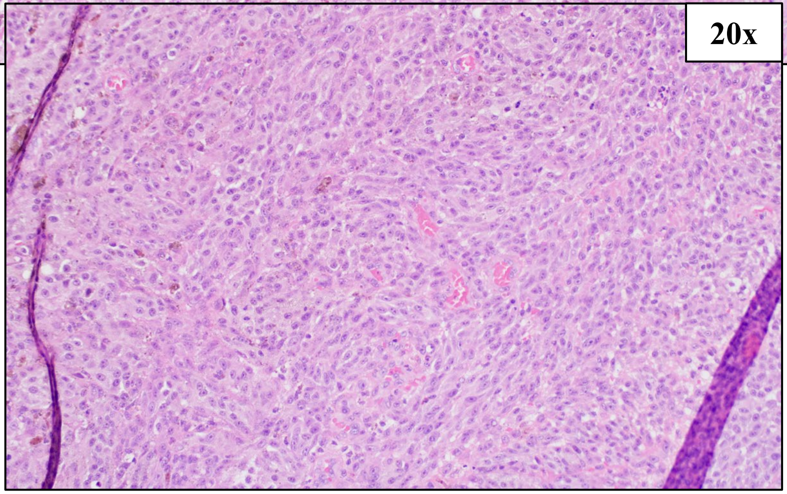


IHC Stains:
SOX10+

NRAS Q61R

### HCI-AM093, Acral Melanoma

#### SNV (%VAF)

NRAS c.182A>G, p.Q61R (99%)

#### CNV

Amplification: TERT, CRKL
Loss: MET, CDKN2A, PTEN, GAB2

#### PDX Tumor Histology

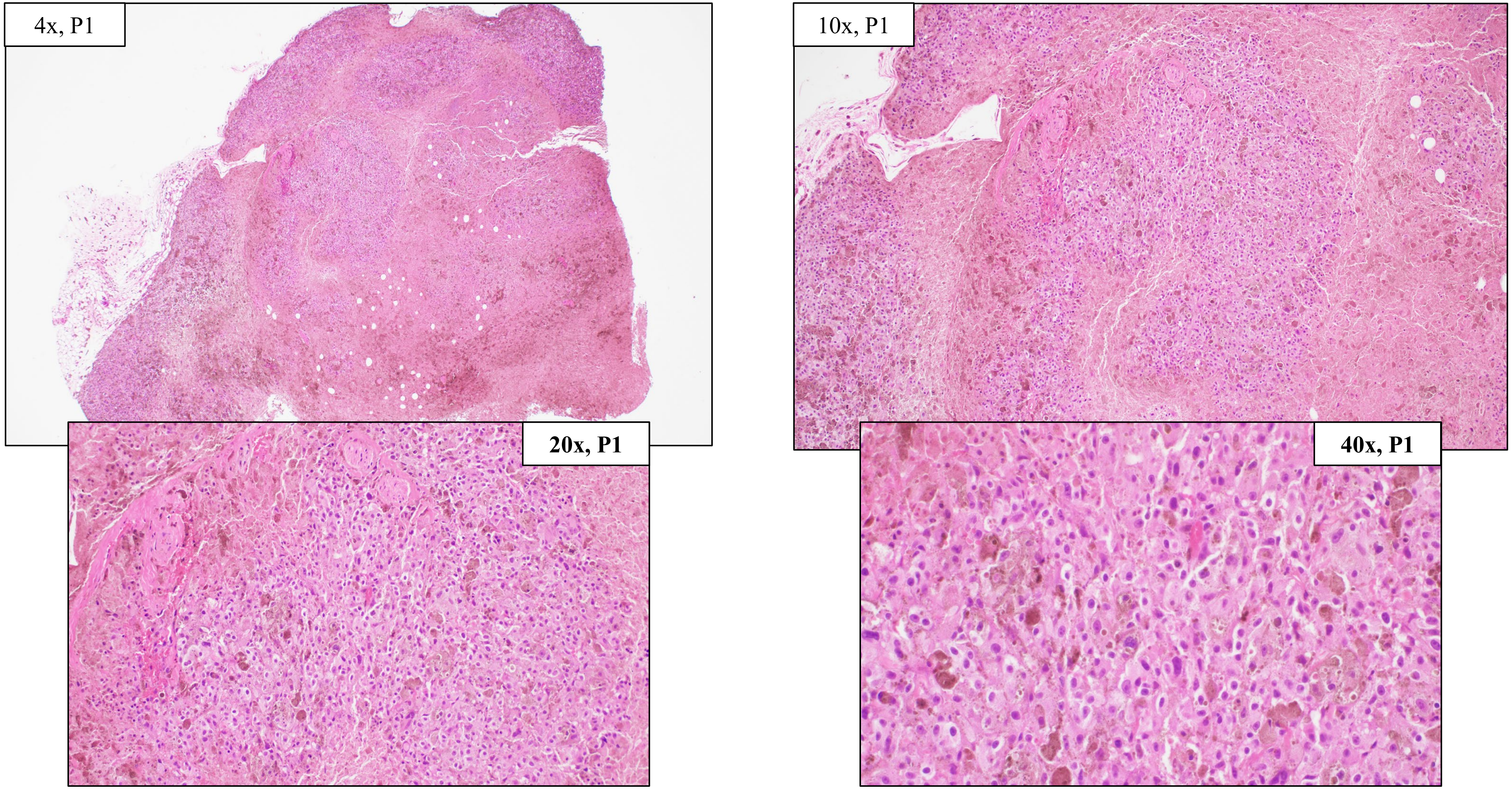


NRAS Q61R
